# Genetic diversity, distribution and domestication history of the neglected GGA^t^A^t^ genepool of wheat

**DOI:** 10.1101/2021.01.10.426084

**Authors:** Ekaterina D. Badaeva, Fedor A. Konovalov, Helmut Knüpffer, Agostino Fricano, Alevtina S. Ruban, Zakaria Kehel, Svyatoslav A. Zoshchuk, Sergei A. Surzhikov, Kerstin Neumann, Andreas Graner, Karl Hammer, Anna Filatenko, Amy Bogaard, Glynis Jones, Hakan Özkan, Benjamin Kilian

## Abstract

Wheat yields are stagnating around the world and new sources of genes for resistance or tolerances to abiotic traits are required. In this context, the tetraploid wheat wild relatives are among the key candidates for wheat improvement. Despite of its potential huge value for wheat breeding, the tetraploid GGA^t^A^t^ genepool is largely neglected. Understanding the population structure, native distribution range, intraspecific variation of the entire tetraploid GGA^t^A^t^ genepool and its domestication history would further its use for wheat improvement. We report the first comprehensive survey of genomic and cytogenetic diversity sampling the full breadth and depth of the tetraploid GGA^t^A^t^ genepool. We show that the extant GGA^t^A^t^ genepool consists of three distinct lineages. We provide detailed insights into the cytogenetic composition of GGA^t^A^t^ wheats, revealed group-, and population-specific markers and show that chromosomal rearrangements play an important role in intraspecific diversity of *T. araraticum*. We discuss the origin and domestication history of the GGA^t^A^t^ lineages in the context of state-of-the-art archaeobotanical finds. We shed new light on the complex evolutionary history of the GGA^t^A^t^ wheat genepool. We provide the basis for an increased use of the GGA^t^A^t^ wheat genepool for wheat improvement. The findings have implications for our understanding of the origins of agriculture in southwest Asia.

## Introduction

The domestication of plants since the Neolithic Age resulted in the crops that feed the world today. However, successive rounds of selection during the history of domestication led to a reduction in genetic diversity, which now limits the ability of the crops to further evolve (Tanksley and McCouch 1997; van Heerwaarden et al. 2010). This is exacerbated by the demand for high crop productivity under climate change. Crop wild relatives (CWR) represent a large pool of beneficial allelic variation and are urgently required to improve the elite genepools (Dempewolf et al. 2017; Kilian et al. 2021). Bread wheat and durum wheat are the staple crops for about 40% of the world’s population. But as wheat yields are stagnating around the world (Iizumi et al. 2017; Ray et al. 2013; Ray et al. 2012); new sources of genes for resistance or tolerances to abiotic traits such as drought and heat are required. In this context, the wheat wild relatives are among the key sources for bread wheat (*T. aestivum* L., 2n = 6x = 42, BBAADD) and durum wheat (*T. durum* Desf., 2n = 4x = 42, BBAA) improvement (Dante et al. 2013; Zhang et al. 2016).

However, in nature, no wild hexaploid wheat has ever been found. Only two wild tetraploid wheat species (2n = 4x = 28) were discovered, namely, (i) wild emmer wheat *T. dicoccoides* (Körn. ex Asch. et Graebn.) Körn. ex Schweinf. [syn. *T. turgidum* subsp. *dicoccoides* (Körn. ex Asch. & Graebn.) Thell.] and (ii) Armenian, or Araratian emmer *T. araraticum* Jakubz. (syn. *T. timopheevii* (Zhuk.) Zhuk. subsp. *armeniacum* (Jakubz.) van Slageren). Morphologically, both species are very similar but differ in their genome constitution (Zohary et al. 2012). *Triticum dicoccoides* has the genome formula BBAA and *T. araraticum* has GGA^t^A^t^ (Jiang and Gill 1994).

The wheat section Timopheevii mainly consists of wild tetraploid *Triticum araraticum* (GGA^t^A^t^), domesticated tetraploid *T. timopheevii* (Zhuk.) Zhuk. (Timopheev’s wheat, GGA^t^A^t^), and hexaploid *T. zhukovskyi* Menabde et Ericzjan (2n = 6x= 42, GGA^t^A^t^A^m^A^m^) (Dorofeev et al. 1979; Goncharov 2012).

Wild *T. araraticum* was first collected by M.G. Tumanyan and A.G. Araratyan during 1925–28 southeast of Erevan, Armenia (Nazarova 2007), soon after the discovery of domesticated *T. timopheevii* by P.M. Zhukovsky (Zhukovsky 1928) (Supplementary Material S1). Subsequently, *T. araraticum* was found in several other locations in Armenia and Azerbaijan (Dorofeev et al. 1979; Jakubziner 1933, 1959), as well as in Iran, Iraq and Turkey. Single herbarium specimens resembling *T. araraticum* have been sporadically recorded among *T. dicoccoides* accessions collected from the Fertile Crescent (Jakubziner 1932; Sachs 1953). However, only botanical expeditions from the University of California at Riverside (USA) to Turkey in 1965, to the Fertile Crescent in 1972–73 (Johnson and Hall 1967; Johnson and Waines 1977) and the Botanical Expedition of Kyoto University to the Northern Highlands of Mesopotamia in 1970 (Tanaka and Ishii 1973; Tanaka and Kawahara 1976) significantly expanded our understanding of the natural distribution of *T. araraticum*. More recently *T. araraticum* was found in north-western Syria (Valkoun et al. 1998). Especially in south-eastern Anatolia, Turkey, the distribution area of *T. araraticum* overlaps with the distribution range of *T. dicoccoides*. From the western to eastern Fertile Crescent, it is assumed that *T. araraticum* gradually substitutes *T. dicoccoides* (Johnson 1975), and *T. dicoccoides* is absent from Transcaucasia (Özkan et al. 2011). In most habitats, *T. araraticum* grows in patches and in mixed stands with other wild cereals (Troitzky 1932; Tumanyan 1930).

It is difficult to distinguish *T. araraticum* from *T. dicoccoides* by morphology under field conditions (Dagan and Zohary 1970; Tanaka and Sakamoto 1979). However, both species can easily be differentiated based on biochemical, immunological, cytological and molecular markers (Badaeva et al. 1994; Gill and Chen 1987; Jiang and Gill 1994; Kawahara and Tanaka 1977; Konarev et al. 1976; Lilienfeld and Kihara 1934). From an archaeobotanical perspective, both species can be reliably identified based on several characteristics of charred spikelets (Jones et al. 2000). In blind tests, it was possible to distinguish modern representatives of the two species with a c. 90% accuracy, on the basis of the primary keel of the glume, which arises just below the rachis disarticulation scar, and the prominent vein on the secondary keel (observable at the base of the glume, which is the part of spikelet most commonly preserved by charring in archaeological material) (Jones et al. 2000).

According to cytogenetic and molecular analyses, *T. araraticum* originated as a result of hybridization between *Aegilops speltoides* Tausch (2n = 2x= 14, SS) and *Triticum urartu* Thumanjan ex Gandilyan (2n = 2x= 14, AA) independently from *T. dicoccoides* (Dvořák et al. 1988; Rodríguez et al. 2000a,b). Similarity of the cytoplasmic genomes of *Ae. speltoides* and *T. araraticum* indicated that *Ae. speltoides* was the maternal parent of *T. araraticum* (Tsunewaki 1996). Hybrids between *T. araraticum* × *T. dicoccoides* were reported as sterile because of meiotic disturbances and gene interactions (Makushina 1938; Svetozarova 1939; Tanaka and Ishii 1973; Wagenaar 1961), although a few authors (Noda and Ge 1989; Sachs 1953; Tanaka and Ichikawa 1972; Tanaka and Kawahara 1976) reported relatively good chromosome pairing in the F_1_ hybrids in some *T. araraticum* × *T. dicoccoides* combinations.

*Triticum dicoccoides* is considered to be the older species than *T. araraticum* (Gornicki et al. 2014; Huang et al. 2002), which is supported by higher similarity of the S-G genomes compared to the S-B genomes (Jiang and Gill 1993; Kilian et al. 2007; Rodríguez et al. 2000a). Cytogenetic and molecular data showed that the speciation of *T. araraticum* was accompanied by complex species-specific translocations involving chromosomes 1G-6A^t^-4G and 3A^t^-4A^t^ (Chen and Gill 1984; Jiang and Gill 1993; Rodríguez et al. 2000b; Salina et al. 2006) as well as with mutations of the primary DNA structure causing the divergence of homoeologous chromosomes (e.g., chromosomes 3A-3A^t^) (Dobrovolskaya et al. 2009). These changes of karyotype structure are specific for the whole section Timopheevii as compared to the emmer wheat lineage (BBAA, BBAADD) (Badaeva et al. 1986; Hutchinson et al. 1982; Zhang et al. 2013).

Intraspecific diversity of *T. araraticum* was detected by karyotype analysis (Badaeva et al. 1990, 1994; Kawahara et al. 1996; Kawahara and Tanaka 1977) and using nuclear (Nave et al. 2021; Shcherban et al. 2016) and chloroplast DNA markers (Mori et al. 2009).

Most recent phylogenetic studies based on whole chloroplast genome sequences, genome-wide sequence information and enlarged taxon sampling provided increased resolution of the evolutionary history within the Triticeae tribe, thereby shedding also new light on the GGA^t^A^t^ wheat genepool (Bernhardt et al. 2017; Gornicki et al. 2014).

Research and pre-breeding activities have focused on *T. dicoccoides* because it gave rise to the economically most important wheats, *T. durum* Desf. and *T. aestivum* L. (Avni et al. 2017; El Haddad et al. 2021). However, *T. araraticum* and *T. timopheevii* have also contributed to bread wheat improvement. Important genes were transferred such as genes controlling resistance against stem rust, leaf rust, powdery mildew or wheat leaf blotch (Allard and Shands 1954; Brown-Guedira et al. 1996, 2003; Dyck 1992; McIntosh and Gyarfas 1971). Cytoplasmic male sterility (CMS) induced by *T. timopheevii* cytoplasm showed great potential for heterotic hybrid technology (Maan and Lucken 1972; Mikó et al. 2011; Würschum et al. 2017). However, despite of its potential huge value for bread and durum wheat improvement, only a comparatively small number of genes was transferred from *T. timopheevii* (even less from *T. araraticum*). Most of the gene contributions originate from only one line (D-357-1) bred by R. Allard at the University of Wisconsin in 1948 (Martynov et al. 2018).

Understanding the population structure and the intraspecific variation of the entire tetraploid GGA^t^A^t^ genepool would further its use for wheat improvement. In this study, we report the first comprehensive survey of cytogenetic and genomic diversity sampling the full breadth and depth of the tetraploid GGA^t^A^t^ genepool. We provide new insights into the genetic relationships among GGA^t^A^t^ wheats and its domestication, taking into account the state-of-the-art archaeobotanical finds.

## Materials and methods

### Germplasm

A comprehensive germplasm collection of tetraploid wheats was established comprising 862 genebank accessions (1–5 genotypes per accession number): 450 *T. araraticum,* 88 *T. timopheevii,* three *T. militinae* Zhuk. et Migusch., one *T. zhukovskyi,* 307 *T. dicoccoides,* eight *T. dicoccon,* and five *T. durum*. Additionally, the following materials were included: (i) four samples of *T. araraticum* collected by Dr. Nelli Hovhannisyan in Armenia; (ii) 17 samples of *T. araraticum* and 12 samples of *T. dicoccoides* collected by Dr. H. Özkan in Turkey; and (iii) 26 samples of *T. dicoccoides* collected by Drs. E. Badaeva, O.M. Raskina and A. Belyayev in Israel. Altogether, 921 accessions of wild and domesticated tetraploid wheats were examined (Supplementary Table S2).

A subset of 787 tetraploid wheat genotypes representing 765 genebank accessions was examined using Sequence-Specific Amplification Polymorphism (SSAP) markers. The subset included 360 genotypes of *T. araraticum*, 76 *T. timopheevii* (including two *T. militinae*), while 351 genotypes of *T. dicoccoides* were considered as an outgroup. Of them, 243 (67%) *T. araraticum,* 139 (39.6%), and nine *T. timopheevii* genotypes (including one *T. zhukovsky*) were shared with C-banding analysis. The whole collection was single-seed descended (SSD) at least twice under field conditions (2009–2012) and taxonomically re-identified in the field at IPK, Gatersleben in 2011 (Supplementary Table S2).

Based on the results of the SSAP analysis, a subset of 103 genotypes, including 37 *T. araraticum* collected from different geographic regions and representing all genetic groups, one *T. militinae*, 14 *T. timopheevii*, 38 *T. dicoccoides,* 9 *T. dicoccon,* and four *T. durum* was selected for a complementary analysis using Amplified Fragment Length Polymorphism (AFLP) markers to infer the population structure of GGA^t^A^t^ wheats (Supplementary Table S2). Forty-seven *T. araraticum* and *T. timopheevii* accessions (88.7%) were shared with SSAP and 39 (73.6%) with C-banding analyses.

Karyotype diversity of 370 *T. araraticum* accessions was assessed by C-banding and Fluorescence *in situ* hybridization (FISH) in comparison with 17 *T. timopheevii* and one *T. zhukovskyi* genotypes (Supplementary Table S2). According to the C-banding analysis, most *T. araraticum* accessions (353 of 370) were karyotypically uniform and were treated as single genotype each. Seventeen accessions were heterogeneous and consisted of two (13 accessions) or even three (four accessions) cytogenetically distinct genotypes, which were treated as different entities (genotypes). In order to infer the population structure of GGA^t^A^t^ wheats, 265 typical genotypes of *T. araraticum* were selected representing all karyotypic variants, seven *T. timopheevii,* one *T. zhukovskyi.* A total of 87 *T. araraticum* genotypes representing all geographic regions and chromosomal groups were selected for Fluorescence i*n situ* hybridization (FISH) analysis.

### Molecular analysis

#### SSAP analysis

Evolutionary relationships among the comprehensive collection of wild and domesticated tetraploid wheat taxa were first inferred based on polymorphic retrotransposon insertions. For this, the highly multiplex genome fingerprinting method SSAP was implemented based on polymorphic insertions of retrotransposon families *BARE-1* and *Jeli* spread across the wheat chromosomes. DNA was isolated from freeze-dried leaves of 787 SSD plants, using the Qiagen DNeasy Kit (Hilden, Germany). The SSAP protocol was based on Konovalov et al. (2010) with further optimizations for capillary-based fragment detection (Supplementary Material S3). In total, 656 polymorphic markers were generated for BBAA- and GGA^t^A^t^-genome wheats by amplification of multiple retrotransposon insertion sites. Data analysis was performed in SplitsTree 4.15.1 (Huson and Bryant 2006). NeighborNet planar graphs of Dice distances (Dice 1945) were constructed based on presence/absence of SSAP bands in the samples.

#### AFLP analysis

The AFLP protocol, as described by Zabeau and Vos (1993) was performed with minor modifications according to Altıntas□ et al. (2008) and Alsaleh et al. (2015). In total, six AFLP primer combinations were used to screen the collection of 103 lines (Supplementary Material S3). Neighbor-Joining (NJ) trees were computed based on Jaccard distances (Jaccard 1908; Perrier et al. 2003). NeighborNet planar graphs were generated based on Hamming distances (Huson and Bryant 2006). Genetic diversity parameters and genetic distances were calculated using Genalex 6.5 (Peakall and Smouse 2012).

### Cytogenetic analysis

#### C-banding

Chromosomal preparation and C-banding procedure followed the protocol published by Badaeva et al. (1994). The A^t^- and G-genome chromosomes were classified according to the nomenclature proposed by Badaeva et al. (1991) except for chromosomes 3A^t^ and 4A^t^. Based on meiotic analysis of the F_1_ *T. timopheevii* × *T. turgidum* hybrids (Rodríguez et al. 2000b) and considering karyotype structure of the synthetic wheat *T.* × *soveticum* (Zhebrak) (Mitrofanova et al. 2016), the chromosomes 3A^t^ and 4A^t^ were exchanged. Population structure of *T. araraticum* and the phylogenetic relationship with *T. timopheevii* were inferred based on chromosomal passports compiled for 265 genotypes representing all geographic regions and chromosomal groups (247 *T. araraticum*, 17 *T. timopheevii*, one *T. zhukovskyi*). Chromosomal passports were constructed by comparing the karyotype of the particular accession with the generalized idiogram of A^t^- and G-genome chromosomes (Fig. 1; Supplementary Material S3), as described for *T. dicoccon* Schrank (Badaeva et al. 2015b).

**Fig. 1.**
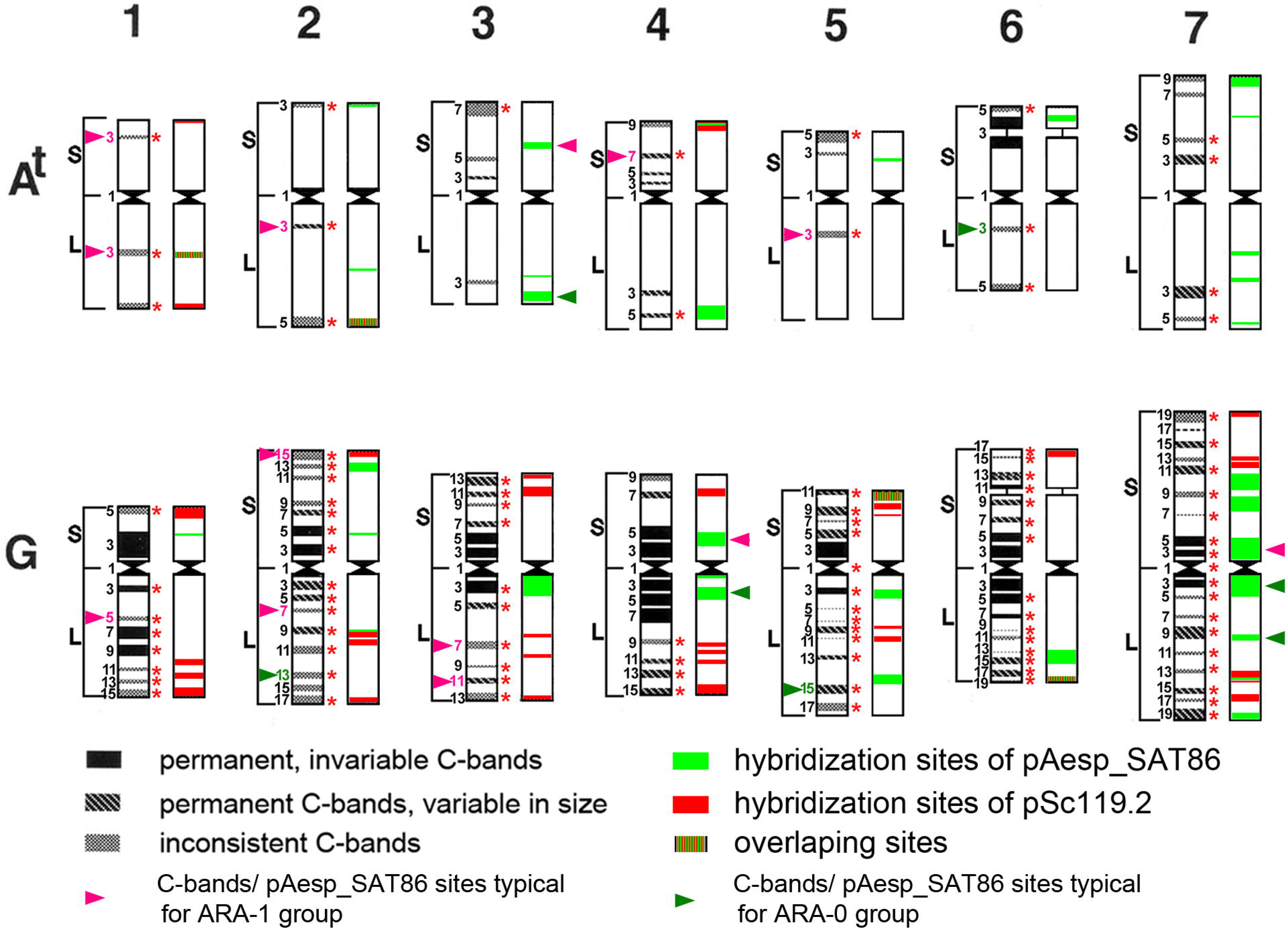
Generalized idiogram and nomenclature of the A^t^- and G-genome chromosomes. The C-banding pattern is shown on the left, the pSc119.2 (red) and pAesp_SAT86 (green) pattern on the right side of each chromosome. 1–7 – homoeologous groups; S – short arm, L – long arm. The numerals on the left-hand side designate putative positions of C-bands/FISH sites that can be detected on the chromosome arm; C-bands specific for the ARA-1 group are shown with pink numerals, C-bands specific for the ARA-0 group are indicated by green numerals. Red asterisks on the right-hand side indicate C-bands that were considered for the “chromosomal passport”.

### Fluorescence *in situ* hybridization (FISH)

Two polymorphic G-genome specific DNA probes, Spelt-1 (Salina et al. 1998) and Spelt-52 (Salina et al. 2004), were used for screening all 95 (87 *T. araraticum,* seven *T. timopheevii* and one *T. zhukovskyi*) genotypes considered for FISH analysis. Additionally, 26 of the 95 genotypes were analyzed using the probe pAesp_SAT86 (Badaeva et al. 2015a), which also showed differences between the accessions. The probes pSc119.2 (Bedbrook et al. 1980), GAA_n_ (Pedersen et al. 1996), and pTa-535 (Komuro et al. 2013) were subsequently hybridized to the same chromosomal spread to allow chromosome identification. Classification of pSc119.2- and pTa-535-labeled chromosomes followed the nomenclature of (Badaeva et al. 2016; Jiang and Gill 1993). Chromosomes hybridized with the GAA_10_ microsatellite sequence were classified according to nomenclature suggested for C-banded chromosomes.

## Results

### Genetic diversity and population structure of the GGA^t^A^t^ genepool

Simultaneous amplification of multiple retrotransposon insertion sites using eight Long Terminal Repeat (LTR) primer combinations generated 656 polymorphic Sequence-Specific Amplification Polymorphism (SSAP) markers: 255 markers were obtained for *Jeli* insertions and 401 markers for *BARE-1* insertions. According to our previous study (Konovalov et al. 2010), *Jeli* targets mostly the A genome, while *BARE-1* is distributed between A- and B/G-genome chromosomes. Altogether, 787 wheat genotypes representing 753 genebank accessions were considered for data analysis, after excluding apparently misidentified taxa and several cases of low-yield DNA extraction (Supplementary Table S2, column 6). Several major observations were made: (i) the extant GGA^t^A^t^ genepool consists of three distinct lineages (two *T. araraticum* lineages and one of *T. timopheevii*, TIM). Surprisingly, wild *T. araraticum* consists of two major genetic lineages, preliminarily designated as ‘ARA-0’ and ‘ARA-1’ (Fig. 2; Supplementary Figure S4; Supplementary Figure S5); (ii) while ARA-0 was found to be geographically widespread, ARA-1 was only found in south-eastern Turkey (Adiyaman, Kahramanmaraş, Gaziantep, Kilis) and in north-western Syria, where the distribution ranges of *T. araraticum* and of *T. dicoccoides* overlap (Fig. 3); (iii) as expected, *T. dicoccoides* is genetically more diverse, supporting a more recent origin of *T. araraticum* (Supplementary Figure S4; Supplementary Table S6); (iv) among the GGA^t^A^t^ lineages, ARA-0 harbors more genetic diversity (Supplementary Table S6); (v) differences between A^t^-and A^t^+G genome diversity patterns based on *Jeli* and *BARE-1* were discovered, respectively (Supplementary Table S6); and (vi) Nei’s genetic distance between lineages based on all 656 SSAP markers or considering only the *BARE-1* markers revealed that ARA-0 is phylogenetically more closely related to TIM than ARA-1. However, considering only the *Jeli* markers, ARA-1 was more closely related to TIM. *Triticum dicoccoides* (DIC) was genetically related most closely to the ARA-1 lineage (Supplementary Table S6).

**Fig. 2.**
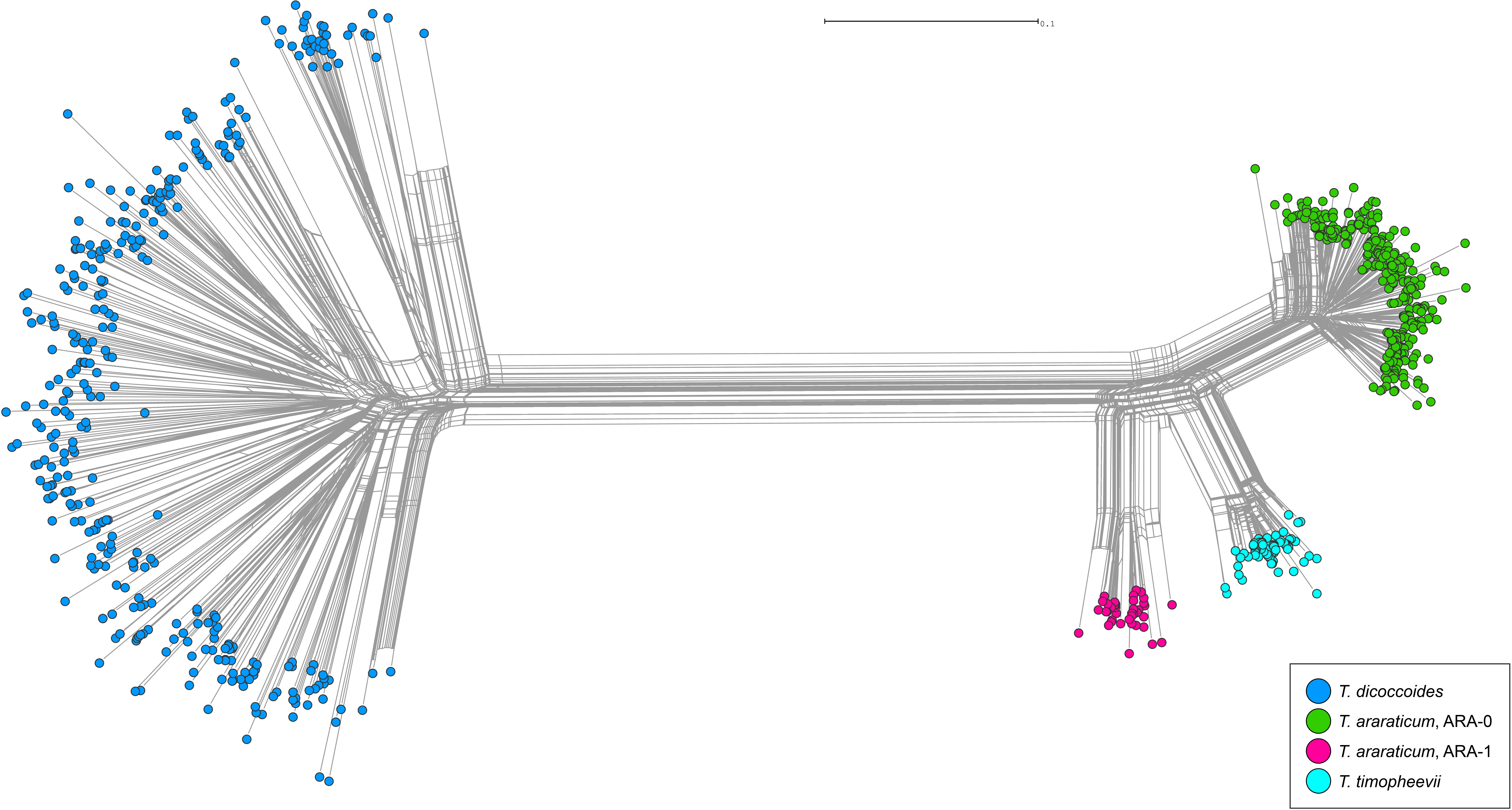
Genetic relationships between GGA^t^A^t^ and BBAA wheats. NeighborNet planar graph of Dice distances representing the diversity of 787 GGA^t^A^t^ and BBAA tetraploid wheat genotypes based on 656 SSAP markers.

**Fig. 3.**
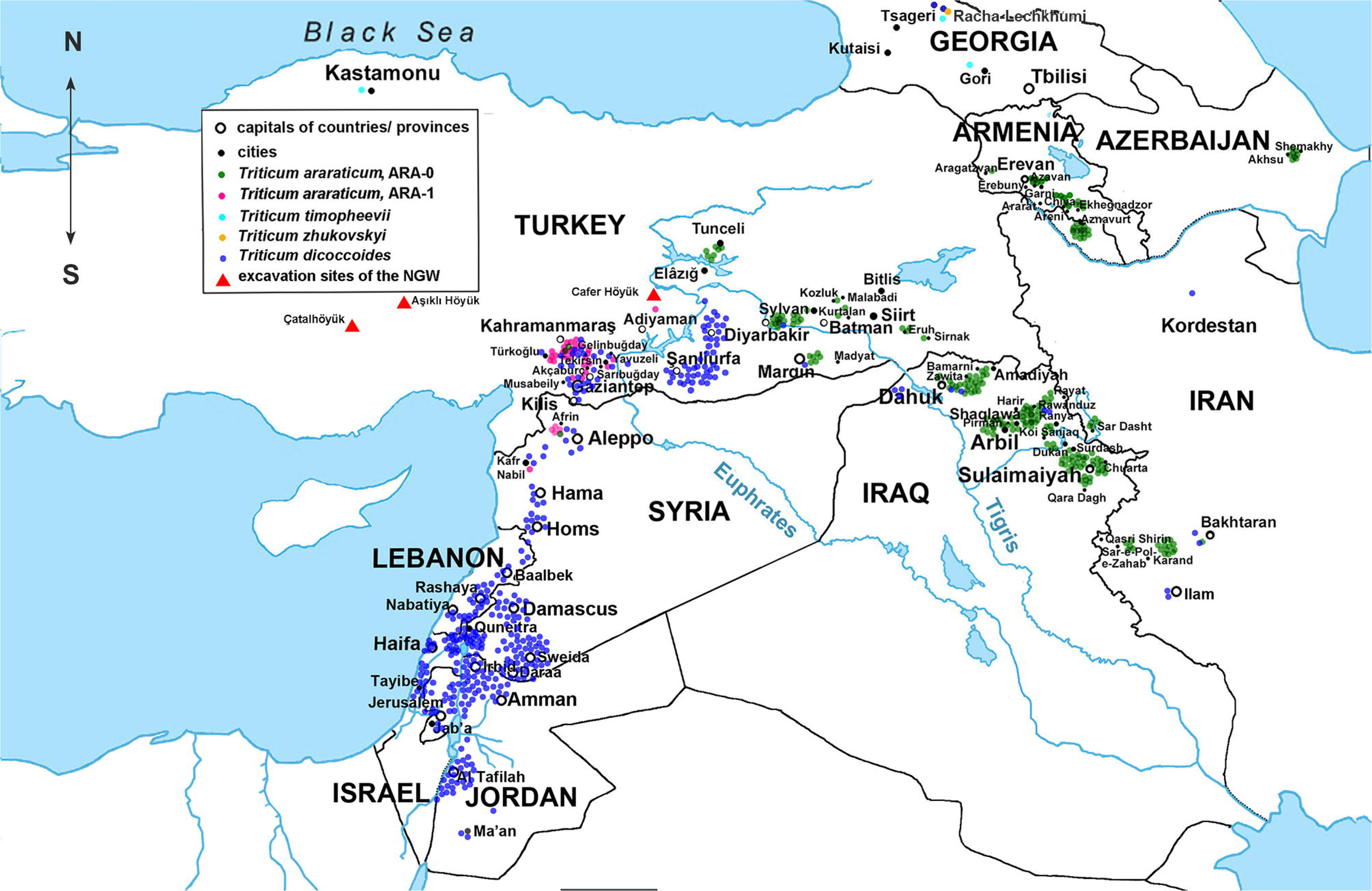
Natural geographical distribution of wild tetraploid *T. araraticum* and *T. dicoccoides*. Green dots correspond to collection sites of ARA-0 accessions, pink dots to ARA-1, and dark blue dots to *T. dicoccoides* (DIC). The collection sites of *T. timopheevii* and *T. zhukovskyi* are shown with turquoise and yellow dots, respectively. Key excavation sites in Turkey where NGW was identified are indicated with red triangles.

A carefully selected subset of 103 genotypeswas used for Amplified Fragment Length Polymorphism (AFLP) analysis (Supplementary Table S2, column 8). A screen using the six most promising AFLP primer combinations uncovered a total of 146 polymorphic markers across all genotypes. Major findings can be summarized as: (i) clustering of genotypes based on AFLP markers corroborates the existence of the three major and genetically distinct lineages of GGA^t^A^t^ wheats (Supplementary Figure S7). One lineage comprised all *T. timopheevii* (and its derived mutant *T. militinae* Zhuk. et Migusch.) genotypes. Importantly, the two lineages of *T. araraticum* and their distribution ranges were verified; (ii) the summary statistics highlight that, in contrast to the SSAP analysis, ARA-1 was more diverse than TIM for all parameters (Supplementary Table S8); (iii) ARA-1 was found to be genetically most closely related to TIM (Supplementary Table S8); (iv) ARA-1 was related most closely to DIC Supplementary Table S8); (v) potential hybridization signals between ARA-0 and ARA-1 lineages were identified (Supplementary Figure S7, purple split); (vi) ARA-1 lines collected around Kilis, Kahramanmara and Gaziantep in Turkey were genetically closest to TIM (Supplementary Figure S7, purple split); and (vii) shared splits between ARA-0 genotypes collected in Armenia and Azerbaijan with TIM (Supplementary Figure S7, purple split) indicate the potential contribution of ARA-0 to the formation of TIM or probably hybridization of ARA-0 with TIM. The genetically closest ARA-0 genotype to TIM was collected from the present Ararat province of Armenia.

In total, 248 genotypes of *T. araraticum* and 17 of *T. timopheevii* collected across the entire distribution range and representing all karyotypic variants (Supplementary Table S2, column 12) were selected to infer the population structure based on chromosomal passports (Badaeva et al. 2015b). The results based on 96 informative C-bands supported the molecular findings using *Jeli* markers and AFLP markers and were congruent with the AFLP marker results as the closest ARA-1 genotypes to TIM were collected near Gaziantep (61 km SE from Türko lu, SW of Karadağ), Turkey (Fig. 3; Supplementary Figure S9; Supplementary Table S10). Fluorescence *in situ* hybridization (FISH) using six DNA probes was carried out for 95 genotypes (Supplementary Table S2, column 13), which, in turn, corroborates the findings obtained using *Jeli* markers, AFLP and C-banding markers (Supplementary Table S11).

### Intraspecific genetic diversity of T. araraticum based on karyotype structure and C-banding patterns

Cytogenetic analysis of 391 *T. araraticum* genotypes in comparison with 17 *T. timopheevii* genotypes provided detailed insights into the genetic composition of GGA^t^A^t^ wheats. First, cytogenetic analysis highlighted significant differences between wild emmer *T. dicoccoides* and *T. araraticum/T. timopheevii* in karyotype structure and C-banding patterns (Fig. 4). Second, we revealed high diversity of the C-banding patterns and broad translocation polymorphisms within *T. araraticum* (Figs. 4, 5). The karyotype lacking chromosomal rearrangements was defind as ‘normal’ (N) and was the most frequent karyotype variant shared by *T. timopheevii* and *T. araraticum*. The ‘normal’ karyotype was found in 175 of 391 *T. araraticum* genotypes (44.6%) (more specifically: 155 of 342 ARA-0 = 45.32%; 20 of 49 ARA-1 = 40.81%). These 175 genotypes differed from each other only in the presence/absence or size of one to several C-bands. The ratio of karyotypically normal genotypes decreased from 86.7% in Azerbaijan (excluding Nakhichevan), to 60.0% in Turkey, 46.5% in Iraq, 30.6% in Armenia, 20.0% in Nakhichevan, Azerbaijan, 16.7% in Syria, to 9.1% in Iran (Supplementary Figure S12). Local populations differed in the ratio of normal/rearranged genotypes. For example, some populations from Dahuk, Iraq, possessed only karyotypically normal genotypes, while in others all genotypes possessed chromosomal rearrangements. Similarly, in Turkey the frequency of karyotypically normal genotypes varied from 100% (Mardin) to 0% (Kilis) (Supplementary Table S13). C-banding patterns of *T. araraticum* were highly polymorphic. Based on the presence or absence of particular C-bands (Figs. 1, 4), all genotypes were divided into two groups. The first, larger group (ARA-0) comprised of 342 genotypes (Supplementary Figures S14 - S19). The second group (ARA-1) included 49 genotypes (Supplementary Figure S18 *h1–h5; h6–h9*; Supplementary Figure S20). Interestingly, genebank accession TA1900 presumably collected 32 km S of Denizli, Dulkadiroğlu district, Kahramanmaraş province in the Taurus Mountain Range of Turkey in 1959 shared karyotypic features of *T. timopheevii* and ARA-0 (Fig. 4) and, probably, it is a natural or artificially produced hybrid of *T. timopheevii* with an unknown genotype of *T. araraticum*.

**Fig. 4.**
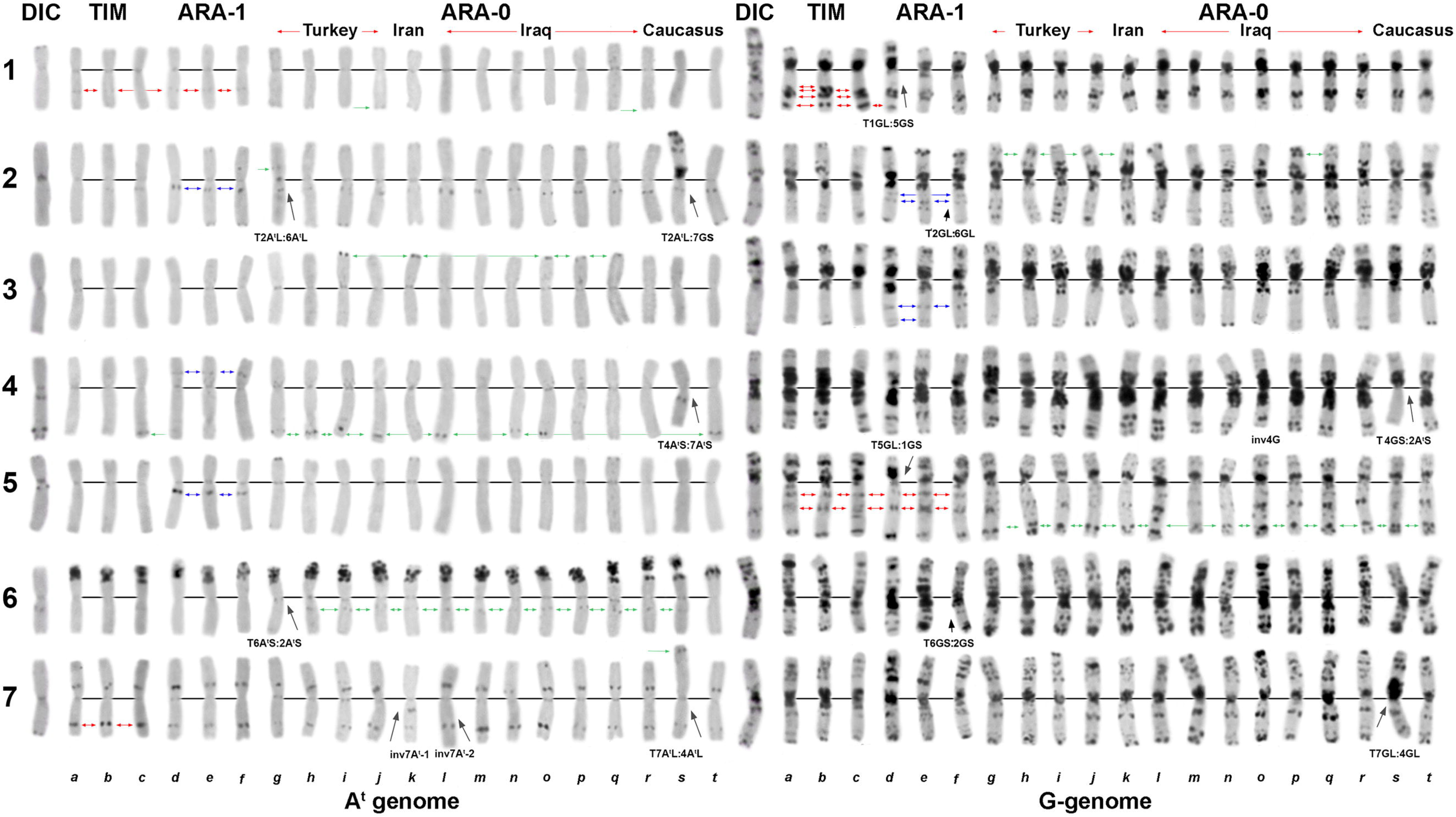
Comparison of the C-banding patterns of *T. dicoccoides* (DIC), *T. timopheevii* (TIM, *a–c,* normal karyotypes), *T. araraticum* ARA-1 (*d–f*) and ARA-0 (*g–t*). DIC (IG 117174, Gaziantep), *a –* KU-1818 (Georgia); *b –* PI 119442; *c –* TA1900; *d –* IG 116165; *e –* PI 654340; *f –* KU-1950; *g –* CItr 17677; *h –* KU-8917; *i –* KU-8909; *j –* KU-1933 (all from Turkey); *k –* CItr 17680 (Iran); *l –* PI 427381 (Erbil, Iraq); *m –* PI 538518*; n –* PI 427425 (Dahuk, Iraq); *o –* KU-8705; *p –* KU-8695 (Shaqlawa, Erbil, Iraq); *q –* KU-8451; *r* – KU-8774 (Sulaymaniyah, Iraq); *s –* TRI 11945 (Nakhichevan); *t –* KU-1901 (Armenia). 1–7 – homoeologous groups. C-bands typical for ARA-1 are indicated with blue arrows, for ARA-0 – with green arrows, and C-bands characteristic for *T. timopheevii –* with red arrows. Black arrows point to rearranged chromosomes in genotypes.

**Fig. 5.**
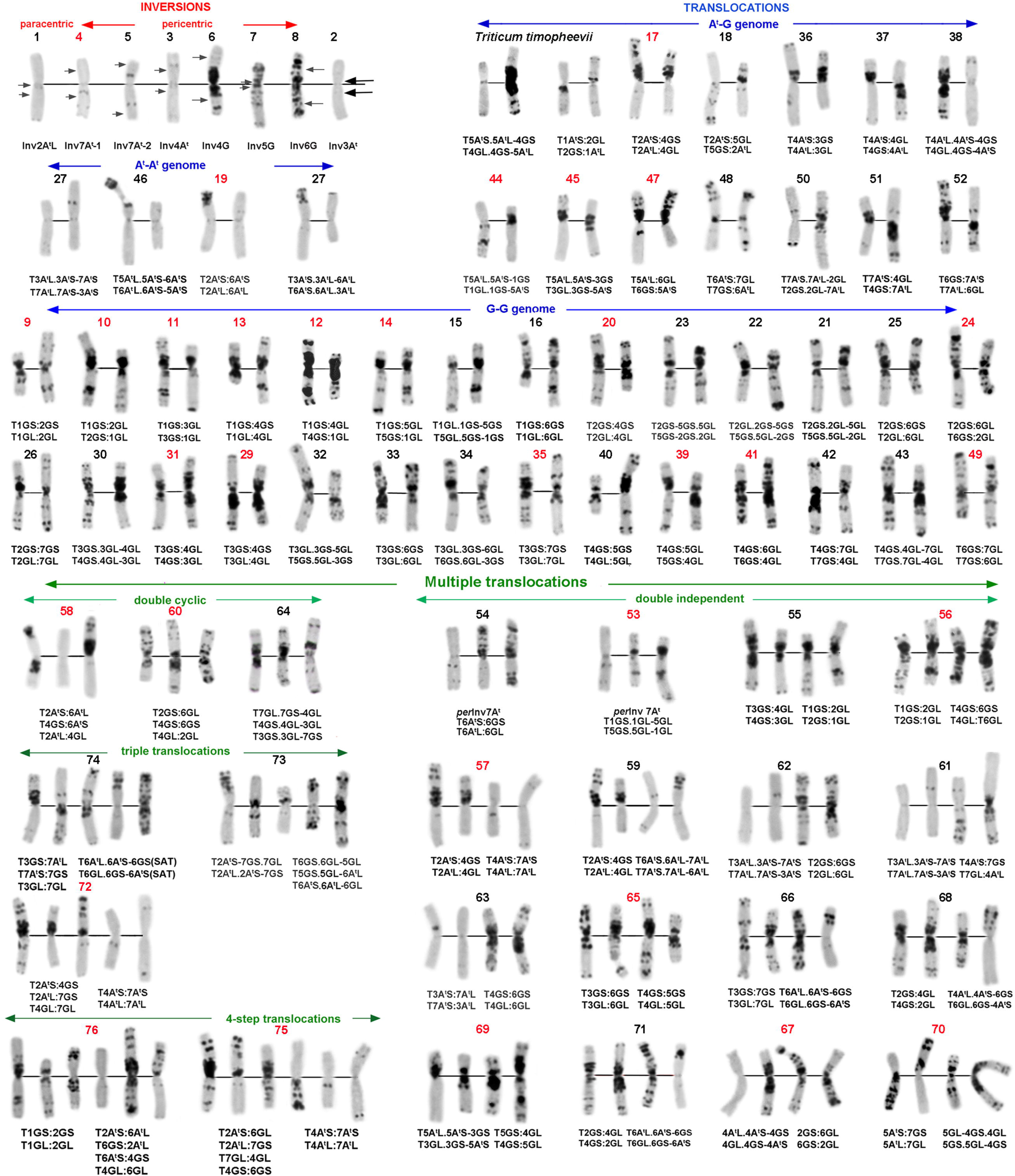
Chromosomal rearrangements identified in *T. araraticum*. The number of translocation variant corresponds to the number of the respective variant in Supplementary Table S13. Novel variants are designated with black numbers, and already known variants by red numbers.

Six chromosomes carried diagnostic C-bands for the ARA-1 lineage at the following positions: 1A^t^L3, 4A^t^S7, 5A^t^L3, 1GL5, 2GL7, 3GL7+L11 (Fig. 1). All these diagnostic C-bands were present in all ARA-1 genotypes, both with normal and rearranged karyotypes (Supplementary Figure S18, Supplementary Fig. S20). Some C-bands were also common for both ARA-0 and ARA-1 (e.g., 2A^t^L3 and 2GS15, Fig. 4), but their size was larger in ARA-1 genotypes.

Only few C-bands were characteristic for the ARA-0 lineage (Figs. 1, 4). Three C-bands appeared with a frequency of over 95%, and two of them, 6A^t^L3 and 5GL15, occurred only in the ARA-0 group. The third C-band, 2GL13, was detected in 97% of ARA-0, but also in few ARA-1 genotypes.

‘Region-specific’ C-bands (in terms of highest frequency) (Badaeva et al. 1994) were detected for ARA-0. For example, (i) the band 4A^t^L7 dominated in Turkey and Transcaucasi (Supplementary Figure S14, Supplementary Figure S19); (ii) a medium to large 3A^t^S7 band was frequently observed in Iran and Sulaymaniyah (Iraq); (ii) one distinct 5GL13 band was frequently detected in genotypes from Erbil, Iraq (Supplementary Figure S15, Supplementary Figure S16). The specificity of C-banding patterns in genotypes originating from the same geographic region was not only determined by single bands, but usually by a particular combination of C-bands on several chromosomes. These region-specific banding patterns were observed for both, normal and translocated forms. For example, the unique banding pattern of chromosome 7G, lacking the marker C-band 7GS11, but carrying large bands for 7GS13, 7GS17 and 7GS21 on the short arm, and 7GL13 and 7GL15 on the long arm was common in Transcaucasia (Supplementary Figure S14). The chromosome 6G lacking telomeric C-bands on the long arm was frequent in genotypes from Dahuk and Sulaymaniyah (both Iraq), and also occurred in ARA-1 genotypes from Kahramanmara, Turkey (Supplementary Figure S20).

### Chromosomal rearrangements play an important role in intraspecific diversity of T. araraticum

In total, 216 out of 391 (55.4%) *T. araraticum* accessions carried a translocated karyotype. Seventy-six variants of chromosomal rearrangements including single and multiple translocations, paracentric and pericentric inversions were identified (Figs. 5, 6). Novel rearrangements were represented by 44 variants, while 32 variants were described earlier (Badaeva et al. 1990, 1994; Kawahara et al. 1996; Kawahara and Tanaka 1977, 1981). One-hundred-forty-seven genotypes differed from the ‘normal’ karyotype by one, 45 genotypes by two (double translocations), 21 genotypes by three (triple translocations) and three genotypes – by four chromosomal rearrangements (quadruple translocations) (Supplementary Table S13).

**Fig. 6.**
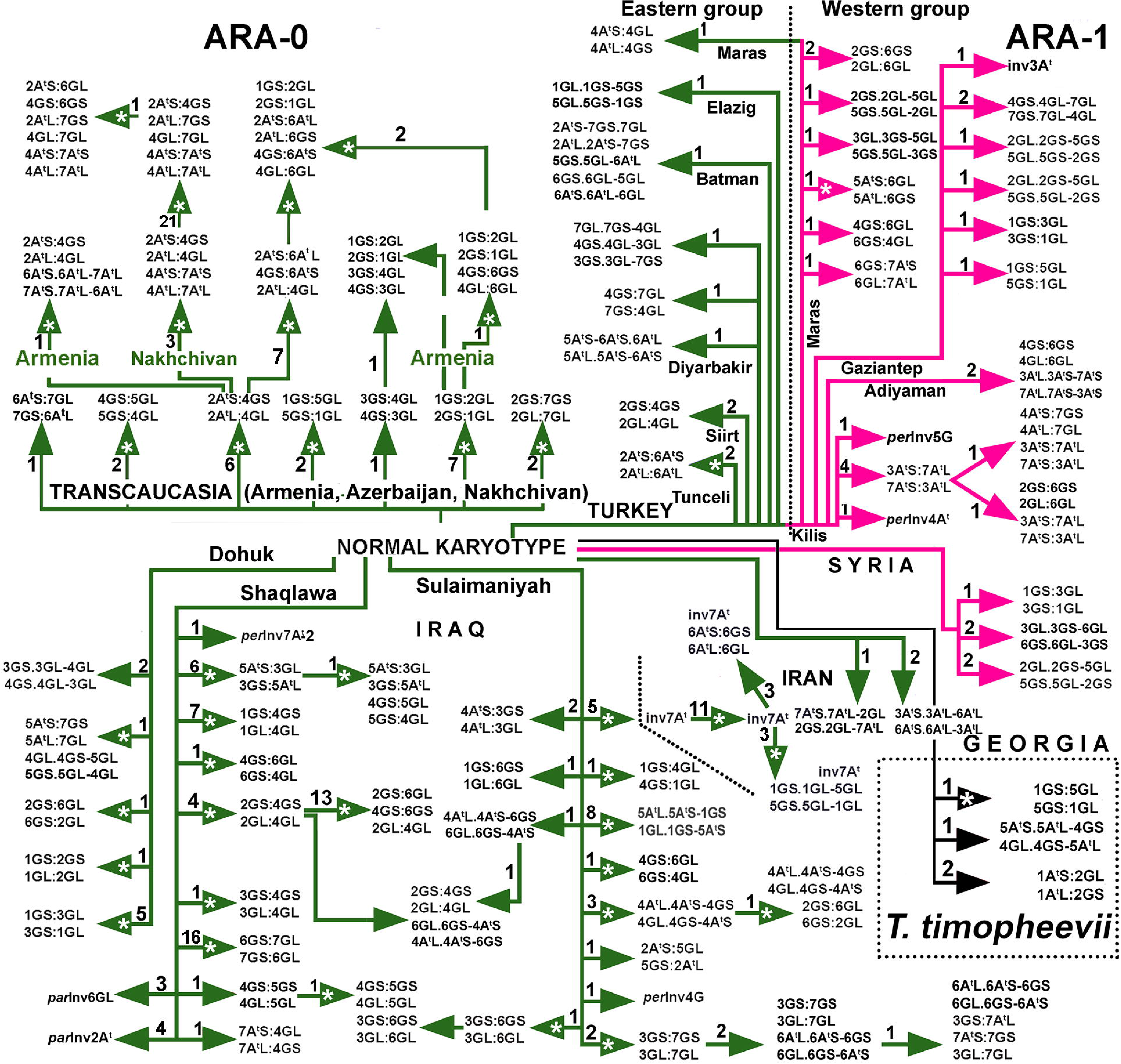
Intraspecific divergence of *T. araraticum* and *T. timopheevii*. Combinations of chromosome arms in rearranged chromosomes are designated. Line colors mark the different groups: ARA-0 (green), ARA-1 (pink) and *T. timopheevii* (black). Solid arrows designate novel rearrangements; arrows with asterisk designate previously described rearrangements (Badaeva et al. 1990, 1994). The numerals above/next to the arrows indicate the number of accessions carrying the respective translocation.

Altogether, we revealed 52 (33 novel) variants of single chromosomal rearrangements (Figs. 5, 6). They included paracentric (one variant) and pericentric inversions (seven variants) and 44 single translocations involving A^t^-A^t^, A^t^-G, or G-G-genome chromosomes. Double rearrangements were represented by 16 independent and three cyclic translocations; among them 10 were novel. Triple translocations were represented by three variants, two of which – T2A^t^:7G + T6A^t^:5G:6G and T3G:7G:7A^t^ + T6A^t^:6G – were found here for the first time. Both variants of quadruple translocations have been identified earlier in Transcaucasia (Badaeva et al. 1990, 1994).

A translocation between two chromosomes could give rise to different products depending on the breakpoint position and arm combination in rearranged chromosomes. For example, a centromeric translocation between 1G and 2G resulted in two translocation variants which were distinct in arm combinations (S:S *vs.* S:L). Three translocation variants involving chromosomes 3G and 4G differed from each other in arm combination and breakpoint position. To discriminate different translocation variants involving same chromosomes, they were designated as T1G:2G-1 and T1G:2G-2, etc. (Supplementary Table S13).

Most variants of chromosomal rearrangements were unique and identified in one or few genotypes, and only four variants were relatively frequent. These were a triple translocation T2A^t^:4G:7G + T4A^t^:7A^t^, *per*Inv7A^t^-1, T6G:7G, and T2G:4G:6G (21, 16, 12, and 13 genotypes, respectively). Taken together, these four variants accounted for approximately 16% of the whole materials we studied. Genotypes carrying the same rearrangement usually had similar C-banding patterns and were collected from the same, or closely located geographic regions. For example,

i. all genotypes with T2A^t^:4G:7G + T4A^t^:7A^t^ originated from Nakhichevan, Azerbaijan; (ii) T6G:7G and T2G:4G:6G were found in Erbil, Iraq; and (iii) *per*Inv7A^t^-1 was collected in Iran and in the neighboring region of Sulaymaniyah, Iraq. Other frequent translocations had a more restricted distribution and usually occurred in a single population. Only few genotypes carrying the same translocations were identified in spatially separated populations. These genotypes differed in their C-banding patterns, for example: (i) T2G:4G was found in four genotypes from Erbil, Iraq and in two genotypes from Siirt, Turkey;
ii. T1G:3G was identified in five ARA-0 genotypes from Dahuk, Iraq, one ARA-1 from Turkey and one ARA-1 genotype of a mixed accession IG 117895 collected in Syria; and (iii) T1G:5G was identified not only in three cytogenetically distinct *T. araraticum* ARA-0 genotypes from Armenia and Azerbaijan and ARA-1 from Turkey (Fig. 6; Supplementary Figure S14, Supplementary Figure S20), but also in *T. timopheevii* (Badaeva et al. 2016).

Most populations consisted of both genotypes with ‘normal’ karyotype and genotypes with one to several variants of chromosomal rearrangements. However, their ratio and spectra differed between regions (Fig. 6; Supplementary Figure S12; Supplementary Material S21).

### Cytogenetic diversity of GGA^t^A^t^ wheats assessed using FISH markers

To further investigate the intraspecific diversity of *T. araraticum* and to assess their phylogenetic relationships with *T. timopheevii*, we carried out FISH using six DNA probes. The probes pTa-535, pSc119.2 and GAA_n_ ensured chromosome identification (Badaeva et al. 2016) whereas Spelt-1, Spelt-52 and pAesp_SAT86 were used to estimate intra- and interspecific variation (Fig. 7).

**Fig. 7.**
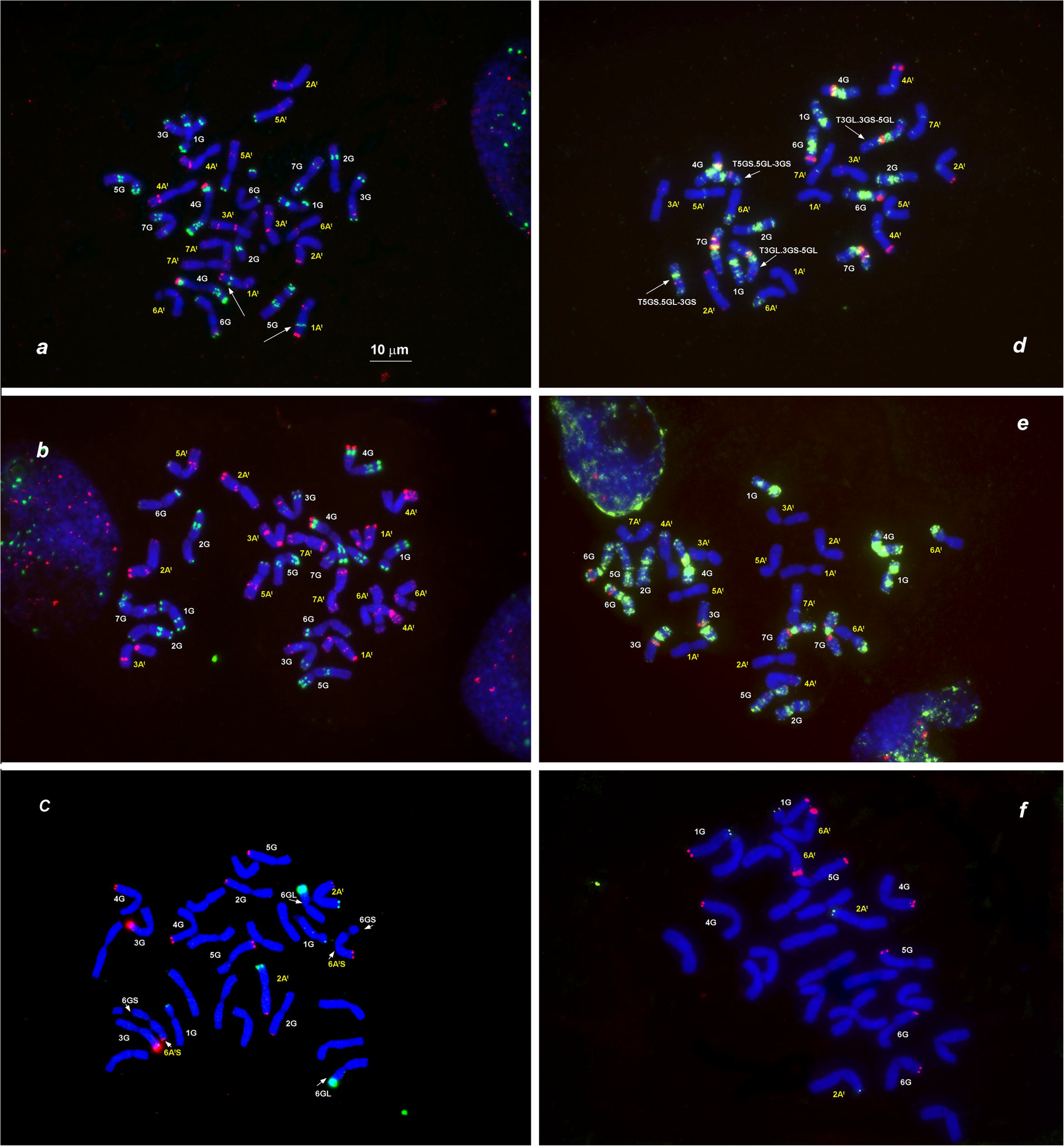
Distribution of different families of tandem repeats on chromosomes of *T. timopheevii* and *T. araraticum*. *Triticum timopheevii,* KU-107 (*a*), and *T. araraticum,* CItr 17680, ARA-0 (*b*), KU-8944, ARA-0 (*c*), KU-1984B, ARA-1 (*d*), PI 427364, ARA-0 (*e*), and 2630, ARA-1 (*f*). The following probe combinations were used: *a, b –* pSc119.2 (green) + pTa-535 (red); *d, e* – pAesp_SAT86 (red) + GAA_n_ (green); *c, f* – Spelt-1 (red) + Spelt-52 (green). The position of pSc119.2 site on 1A^t^ chromosome typical for *T. timopheevii* and ARA-1 is shown with an arrow (*a*). Translocated chromosomes (*c, d*) are arrowed. Chromosomes are designated according to genetic nomenclature; the A^t^-genome chromosomes are designated with yellow numerals and the G-genome chromosomes with white numerals. Scale bar, 10 μm.

The distribution of pTa-535 was monomorphic among *T. araraticum* and *T. timopheevii,* while the pSc119.2 site on 1A^t^L discriminated ARA-1 and TIM from ARA-0 (Fig. 7*a*).

The variability of pAesp_SAT86 hybridization patterns was analyzed in four *T. timopheevii* and 26 *T. araraticum* genotypes, of them seven were from ARA-1 and 19 from ARA-0 lineages (Supplementary Table S2). Distribution of the pAesp_SAT86 probe in all *T. timopheevii* genotypes was similar except for chromosome 7A^t^ in genotype K-38555, which was modified due to a paracentric inversion or insertion of an unknown chromosomal fragment. Labeling patterns, however, were highly polymorphic for *T. araraticum* (Fig. 7; Supplementary Figure S22). Large pAesp_SAT86 sites were found only on some G-genome chromosomes. The A^t^-genome chromosomes possessed several small, but genetically informative polymorphic sites. Some of these sites were lineage-specific. Most obvious differences were observed for 3A^t^, 4G and 7G chromosomes (Fig. 1). Thus, all TIM and ARA-1 genotypes carried the pAesp_SAT86 signal in the middle of 3A^t^S, while for ARA-0 it was located sub-terminally on the long arm. One large pAesp_SAT86 cluster was present on the long arm of 4G in ARA-0, but on the short arm in ARA-1 and TIM. Two large and adjacent pAesp_SAT86 clusters were detected on 7GS in ARA-1 and TIM, but they were split between opposite chromosome arms in all ARA-0 genotypes. Differences between ARA-0 and ARA-1 in pAesp_SAT86 cluster position on 4G and 7G could be caused by pericentric inversions. Some other pAesp_SAT86 sites were identified in either one of the three groups. ARA-1 exhibited the largest polymorphism of pAesp_SAT86 labeling patterns among all groups (Supplementary Figure S22).

FISH with Spelt-1 and Spelt-52 probes on chromosomes of 87 *T. araraticum* genotypes from different chromosomal groups and of different geographic origin, seven *T. timopheevii,* and one *T. zhukovskyi* genotypes revealed high intraspecific diversity of *T. araraticum* and low polymorphism in *T. timopheevii* (Supplementary Figure S23). The broadest spectra of labeling patterns were found in genotypes from Dahuk and Sulaymaniyah (Iraq) and in the ARA-1 group from Turkey, while material from Transcaucasia exhibited the lowest variation. The polymorphism was due to variation in the number of Spelt-1 and Spelt-52 sites, their size and chromosomal location (Supplementary Table S24). The Spelt-1 signals in various combinations appeared in sub-telomeric regions of either one or both arms of 2A^t^, 6A^t^, and all G-genome chromosomes. The Spelt-52 signals were observed in various combinations on 2A^t^S, 1GS, and 6GL chromosomes. Among them, two Spelt-1 sites (6GL and 7GL) and one Spelt-52 locus (2GS) were novel. The patterns of Spelt-1 and Spelt-52 repeats varied across geographic regions and between chromosomal groups (Supplementary Material S25; Supplementary Table S26).

## Discussion

We present the most comprehensive survey of cytogenetic and genomic diversity of GGA^t^A^t^ wheats. We describe the composition, distribution and characteristics of the GGA^t^A^t^ genepool. Building on our results, the latest published complementary genomics studies and state-of-the-art archaeobotanical evidence we revisit the domestication history of the GGA^t^A^t^ wheats. We arrived at the following four key findings:

### 1. The GGA^t^A^t^ genepool consists of three distinct lineages

We sampled the full breadth and depth of GGA^t^A^t^ wheat diversity and discovered a clear genetic and geographic differentiation among extant GGA^t^A^t^ wheats. Surprisingly, and supported by all marker types, three clearly distinct lineages were identified. The first lineage is comprised of all *T. timopheevii* genotypes (and the derived *T. militinae* and *T. zhukovskyi*; note that all *T. militinae* and all *T. zhukovskyi* accessions maintained *ex situ* in genebanks are each derived from only one original genotype). Interestingly, *T. araraticum* consists of two lineages that we preliminarily describe as ‘ARA-0’ and ‘ARA-1’. This finding is in contrast to Kimber and Feldman (1987) who concluded that *T. araraticum* does not contain cryptic species, molecularly distinct from those currently recognized. Based on passport data, ARA-0 was found across the whole predicted area of species distribution. ARA-1 was only detected in south-eastern Turkey and in neighboring north-western Syria. It is interesting to note that only in this part of the Fertile Crescent, the two wild tetraploid wheat species, *T. dicoccoides* and *T. araraticum*, grow in abundance in mixed stands.

Based on Figure 3, collecting gaps are evident for *T. araraticum*, and future collecting missions should focus on four specific regions: (i) between the Euphrates river in the west and the Elazığ – Silvan – Mardin transect region in the east. Interestingly, so far only *T. dicoccoides* was reported from this region; (ii) between Bitlis (Turkey) – Amadiyah (Iraq) in the west and the Armenian border in the east including north-western Iran; (iii) between Adıyaman – Silvan in the south and Tunceli in the north; and (iv) between Hama in the south and west/north-west of Aleppo in Syria.

We found only two *T. araraticum* populations which contained representatives of both ARA lineages: (i) 45 km south-east of Kahramanmaraş to Gaziantep, and (ii) 4 km north of St. Simeon on the road to Afrin in Syria. However, in both cases, the ARA-1 lineage was significantly more frequent than ARA-0. This suggests at least a certain level of taxon boundary between ARA-0 and ARA1 lineages and should be investigated in the future.

Independent support for the existence of two wild *T. araraticum* lineages and their distribution comes from Mori et al. (2009) based on 13 polymorphic chloroplast microsatellite markers (cpSSR) (Supplementary Table S2, column 15). The ‘plastogroup G-2’ was distributed in south-eastern Turkey and northern Syria and was closely related to *Triticum timopheevii* (Mori et al. 2009). However, Gornicki et al. (2014), based on whole chloroplast genome sequence information and sufficient taxon sampling (13 Triticeae species and 1127 accessions; 163 accessions in common with our study, Supplementary Table S2, column 17), provided increased resolution of the chloroplast genome phylogeny and showed that the *T. timopheevii* lineage possibly originated in northern Iraq (*and thus according to our data, belong to the ARA-0 lineage as no ARA-1 occurs in Iraq*). This was supported by Bernhardt et al. (2017), who, based on re-sequencing 194 individuals at the chloroplast locus *ndhF* (2232 bp) and on whole genome chloroplast sequences of 183 individuals representing 15 Triticeae genera, showed that some ARA-0 and TIM genotypes are most closely related. All GGA^t^A^t^ wheats re-sequenced by Bernhardt et al. (2017) were considered in our study (Supplementary Table S2, column 16). Haplotype analysis of the *Brittle rachis 1* (*BTR1-A*) gene in a set of 32 *T. araraticum* in comparison with two *T. timopheevii* accessions (Nave et al. 2021) also showed closer relationships of domesticated *T. timopheevii* to wild *T. araraticum* from Iraq. That is more, one of these accessions, TA102 (PI 538461, 1 km NE of Salahaddin) shared the same haplotype with *T. timopheevii* and it was assigned to ARA-0 group by our study (Supplementary Table S2, column 18).

It is important to note that our results (i.e., the characteristics, composition and geographical distribution of ARA-0 and ARA-1 lineages) are not in agreement with the latest comprehensive taxonomical classification of wheat by Dorofeev et al. (1979), who divided *T. araraticum* into two subspecies: subsp. *kurdistanicum* Dorof. et Migusch. and subsp. *araraticum* (Supplementary Table S2, column 2). We propose to re-classify the GGA^t^A^t^ genepool taxonomically in the future.

### 2. The karyotypic composition of GGA^t^A^t^ wheats is as complex as the phylogenetic history of the GGA^t^A^t^ genepool

Based on karyotype analyses, translocation spectra and distribution of DNA probes, *T. araraticum* populations from Dahuk and Sulaymaniyah (both Iraq) harbored the highest karyotypic diversity among all *T. araraticum* populations studied. We consider the region around Dahuk in Northern Iraq as the center of diversity of *T. araraticum*, and this is probably the region where *T. araraticum* originated. This is supported by Nave et al. (2021), who found the highest haplotype diversity among *T. araraticum* from Iraq, and by Bernhardt et al. (2017) and Gornicki et al. (2014) who traced chloroplast haplotypes from *Aegilops speltoides* growing in Iraq via *T. araraticum* (ARA-0) to *T. timopheevii* and *T. zhukovskyi*.

The karyotype ‘similar’ to those in ‘normal’ *T. timopheevii* was found in 44.6% of all *T. araraticum* genotypes. This is the group of candidates, in which the closest wild relative(s) to *T. timopheevii* is (are) expected. The frequency of the normal karyotype varied among countries and between populations (Supplementary Figure S12). It is interesting to note that the Samaxi-Akhsu population in Azerbaijan and some populations near Dahuk (Iraq) possessed mostly karyotypically normal genotypes. Diagnostic C-bands for the ARA-1 lineage, both with normal and rearranged karyotypes, were 1A^t^L3, 4A^t^S7, 5A^t^L3, 1GL5, 2GL7, 3GL7+L11 (Fig. 1). As expected, the number of C-bands characteristic for ARA-0 was smaller (due to the wide geographical distribution) and only two C-bands were lineage-specific and found in normal as well as translocated genotypes: 6A^t^L3 and 5GL15.

However, some FISH patterns suggested that *T. timopheevii* probably originated in Turkey and probably from ARA-1 (or, ARA-1 and TIM may have originated from a common ancestor, but then diverged). This is supported by the following observations: (i) TIM and ARA-1 carry the pSc119.2 signal in the middle of 1A^t^ long arm, while this site was absent from ARA-0; (ii) all ARA-0 and most ARA-1 possessed the Spelt-52 signal on 6GL, but it is absent in all TIM and five ARA-1 genotypes from Gaziantep-Kilis, Turkey. The distribution of Spelt-1 and Spelt-52 probes on chromosomes of these five genotypes was similar to, and in accession IG 116165 (ARA-1 from Gaziantep) almost identical with TIM; (iii) the pAesp_SAT86 patterns on chromosomes 3A^t^, 4G, and 7G are similar in TIM and ARA-1 but differed from ARA-0. Differences between ARA-1 and TIM based on FISH patterns of some other chromosomes as well as the results of C-banding and molecular analyses suggest that extant ARA-1 genotypes are not the direct progenitors of TIM but that the ARA-1 lineage is most closely related to it. Based on AFLP, C-banding, FISH and *Jeli* retrotransposon markers, TIM was genetically most closely related to ARA-1. Additional evidence for the close relationship between TIM and ARA-1 lineages comes from allelic variation at the *VRN-1* locus of genome A^t^ (Shcherban et al. 2016). This analysis revealed a 2.7 kb deletion in intron 1 of *VRN-A1* in three *T. timopheevii* and four *T. araraticum* accessions, which, according to our data, belong to the ARA-1 lineage. However, at *Vrn-G1*, TIM from Kastamonu in Turkey (PI 119442) shared the same haplotype (*Vrn1Ga*) with ARA-1 samples, while TIM from Georgia harbored haplotype *VRN-G1* as found in ARA-0. These results suggest multiple introgression events and incomplete lineage sorting as suggested by Bernhardt et al. (2017, 2020).

Regular chromosome pairing observed in the F_1_ hybrids of lines with ‘normal’ karyotypes (Kawahara et al. 1996), identified in our study as ARA-0 × ARA-1 (Supplementary Table S2, column 14), suggested that karyotypic differences between ARA-0 and ARA-1 lineages are not associated with structural chromosomal rearrangements such as large translocations or inversions.

The emergence or loss of most lineage-specific Giemsa C-bands (Fig. 3) or FISH loci (Supplementary Figure 22, Supplementary Figure 23) could be due to heterochromatin re-pattering: amplification, elimination or transposition of repetitive DNA sequences. Wide hybridization can also induce changes in C-banding and FISH patterns of *T. araraticum* chromosomes. Changes in pAesp_SAT86 hybridization patterns on 4G and 7G chromosomes, however, are likely to be caused by pericentric inversions, which are also frequent in common wheat (Qi et al. 2006). The role of inversions in inter- and intraspecific divergence is probably underestimated. In our case, it seems possible that divergence between ARA-1/TIM (two inversions) from ARA-0 (no inversion) was associated with at least two pericentric inversions. We did not find any genotype harboring both ARA-0 and ARA-1 specific FISH sites, although ARA-0 and ARA-1 genotypes co-existed in two populations in Turkey and Syria. However, based on FISH (Spelt-1 site on chromosome 6GL and 7GS, respectively), hybridization between certain ARA-1 and ARA-0 lines can be predicted.

Iran occupies a marginal part of the distribution range of *T. araraticum*. An abundance of the pericentric inversion of the 7A^t^ chromosome in the Iranian group indicates that it is derived from Iraq. The karyotypically ‘normal’ genotype was probably introduced to Transcaucasia via Western Azerbaijan (Iran). The low diversity of FISH patterns and the low C-banding polymorphism of *T. araraticum* from Transcaucasia indicate that *T. araraticum* was introduced as a single event. Interestingly, the AFLP data suggested some similarity between ARA-0 from Armenia and Azerbaijan and *T. timopheevii*.

We hypothesize that homoploid hybrid speciation (HHS) (Abbott et al. 2010; Nieto Feliner et al. 2017; Soltis and Soltis 2009) and incomplete lineage sorting may be the possible mechanisms explaining the origin of the ARA-1 lineage. Although this assumption was not experimentally supported, it is favored by some indirect evidence. ARA-1 grows in sympatry and in mixed populations with *T. dicoccoides* (Fig. 3) and is phylogenetically most closely related to *T. dicoccoides* (Supplementary Table S6; Supplementary Table S8). ARA-1 is morphologically more similar to *T. dicoccoides.* Thus, five of 10 misclassified *T. araraticum* accessions belonged to ARA-1 group (Supplementary Table S2), two of which, PI 656871 and IG 116176, were the mix of *T. dicoccoides* and ARA-1. Five misclassified ARA-0 accessions from USDA-ARS collection were from Siirt, Turkey, however in other gene bank three of these accessions were treated as *T. araraticum.* Relatively good chromosome pairing was observed in the F_1_ hybrids of some *T. araraticum x T. timopheevii* combinations (Tanaka and Ichikawa 1972), however, pollen fertility of such hybrids was very low (0.3 – 5.4%). ARA-1 could be derived from ancient hybridization of *T. timopheevii* × *T. dicoccoides*; or alternatively, ARA-1 and TIM could be derived from the hybridization ARA-0 × *T. dicoccoides*.

### 3. Does the *T. timopheevii* population found in western Georgia represent the last remnant of a widespread ancient cultivation area of GGA^t^A^t^ wheats?

Wild emmer *T. dicoccoides* belongs to the first cereals to be domesticated by humans in the Fertile Crescent and the evolution and domestication history of *T. dicoccoides* are relatively well studied (Badaeva et al. 2015b; Civáň et al. 2013; Özkan et al. 2011). Domesticated emmer *Triticum dicoccon* Schrank was a staple crop of Neolithic agriculture, was widely cultivated for over 10,000 years and harbored impressive genetic diversity (Nesbitt and Samuel 1996; Szabo and Hammer 1996; Zaharieva et al. 2010). The domestication of *T. dicoccoides* provided the key for durum wheat (Maccaferri et al. 2019) and bread wheat evolution (Pont et al. 2019). Much less is known about the domestication history of *T. timopheevii*. It is believed that *T. timopheevii* is the domesticated form of *T*. *araraticum* (Dorofeev et al. 1979; Jakubziner 1932). In contrast to *T. dicoccon*, *T. timopheevii* was, since its discovery, considered as a ‘*monomorphous narrowly endemic species*’ (Dekaprelevich and Menabde 1932) cultivated in few villages of western Georgia (Stoletova 1924-25; Zhukovsky 1928) (Supplementary Material S27). Dekaprelevich and Menabde (1932) noticed that the area of cultivation had probably been larger in the past. The last plants of *T. timopheevii in situ* were found by the expedition of the N.I. Vavilov Institute of Plant Genetic Resources (VIR, Russia) in 1983 near the village of Mekvena (Tskhaltubo, Georgia) and deposited in the VIR genebank under accession number K-56422 [E.V. Zuev, personal communication]. Today, the widespread view is that the cultivation area of *T. timopheevii* was restricted to Georgia in the (recent) past (Feldman 2001; Mitrofanova et al. 2016, Zohary et al. 2012).

However, hulled tetraploid wheat morphologically similar to *T. timopheevii* was identified at three Neolithic sites and one Bronze Age site in northern Greece and described by Jones et al. (Jones et al. 2000) as a **‘**New’ Glume Wheat (new glume wheat, NGW). The glume bases of these archaeological finds morphologically resemble *T. timopheevii* more than any other extant domesticated wheat (Jones et al. 2000). After these finds of NGW in Greece, this wheat was also identified at Neolithic and Bronze Age sites in Turkey, Bulgaria, Romania, Hungary, Slovakia, Austria, Italy, Poland, Germany and France (Bieniek 2002, 2007; Bogaard et al. 2007, 2013; Ergun 2018; Fairbairn et al. 2002; Fiorentino and Ulaş 2010; Fischer and Rösch 2004; Hajnalová 2007; Kenéz et al. 2015; Kohler-Schneider 2003; Kreuz and Boenke 2002; Perego 2017; Rottoli and Pessina 2007; Toulemonde et al. 2015; Ulaş and Fiorentino 2020; Valamoti and Kotsakis 2007). Earlier finds of an ‘unusual’ glume wheat in Serbia (Borojevic 1991) and Turkey (de Moulins 1997) have subsequently been recognized as NGW (Kenéz et al. 2015; Jones et al. 2000; Kroll 2016). Criteria were also established for distinguishing the grains of NGW (Kohler-Schneider 2003).

At some sites, NGW appeared as a minor component and may have been part of the accompanying weed flora of cereal fields (Kenéz et al. 2014; Ulaş and Fiorentino 2020). In other cases, it was probably cultivated in a mix with einkorn (Jones et al. 2000; Kohler-Schneider 2003) and/or emmer. The recovery of large quantities in storage deposits of whole spikelets at CCatalhoCyuCk in Turkey, caryopses and spikelet bases at the early Bronze Age settlement of Clermont-Ferrand in France, and rich deposits including whole spikelets at bronze age sites in Italy, demonstrated that, at least in some places, NGW was a major crop in itself (Bogaard et al. 2013, 2017; Ergun 2018; Kenéz et al. 2014; Perego 2017; Toulemonde et al. 2015).

Based on intensive archaeobotanical investigations at CCatalhoCyuCk in central Anatolia, for example, NGW was the predominant hulled wheat, overtaking emmer wheat around 6500 cal BC and remaining dominant until the site’s abandonment c. 5500 cal BC. The finds suggested that this wheat was a distinct crop, processed, stored, and presumably grown, separately from other glume wheats. NGW formed part of a diverse plant food assemblage at Neolithic CCatalhoCyuCk, including six cereals, five pulses and a range of fruits, nuts and other plants, which enabled this early farming community to persist for 1500 years (c. 7100 to 5500 cal BC) (Bogaard et al. 2013, 2017).

Recently, polymerase chain reactions specific for the wheat B and G genomes, and extraction procedures optimized for retrieval of DNA fragments from heat-damaged charred material, have been used to identify archaeological finds of NGW (Czajkowska et al. 2020). DNA sequences from the G genome were detected in two of these samples, the first comprising grain from the mid 7^th^ millennium BC at Çatalhöyük in Turkey, and the second made up of glume bases from the later 5^th^ millennium BC site of Miechowice 4 in Poland. These results provide evidence that NGW is indeed a cultivated member of the GGA^t^A^t^ genepool (Czajkowska et al. 2020). As NGW is a recognized wheat type across a broad geographic area in prehistory, dating back to the 9^th^ millennium BC in SW Asia, this indicates that *T. timopheevii* (*sensu lato, s.l. = domesticated GGA^t^A^t^ wheat in general*), was domesticated from *T. araraticum* during early agriculture, and was widely cultivated in the prehistoric past (Czajkowska et al. 2020).

This raises the question of whether the few populations of *T. timopheevii* (*sensu stricto*, *s.str.*) found in western Georgia were the last remnants of a wider GGA^t^A^t^ wheat cultivation or whether the *T. timopheevii* of Georgia was a local domestication independent of the domestication of *T. araraticum* in SW Asia. To answer this question, sequence information for NGW, the Georgian *T. timopheevii*, and the two lineages of *T. araraticum* (ARA-0, ARA-1) need to be compared.

Vavilov (1935) suggested that *T. timopheevii* of western Georgia was probably originally introduced from north-eastern Turkey. As cited by Dorofeev et al. (1979), Menabde and Ericzjan (1942) associated the origin of *T. timopheevii* with the region of the ancient kingdom of Urartu, whence immigrant ancestors of modern-day Georgians introduced it into western Georgia. Certainly, the possibility of introduction of Timofeev’s wheat into Georgia from the south should not be rejected (Dorofeev et al. 1979). Is the domestication history of *T. timopheevii s.str.* connected with other endemic wheats of Georgia, such as *T. karamyschevii* Nevski (*T. georgicum* Dekapr. et Menabde or *T. paleocolchicum* Menabde) and *T. carthlicum* Nevski, which were cultivated by Mingrelians in Western Georgia (Jorjadze et al. 2014; Mosulishvili et al. 2017)?

### 1. 4. Was the cultivation range of *T. timopheevii* (*s.str.*) wider in the recent past?

We screened all available passport data and found three cases, which could potentially help to reconstruct the recent past cultivation range of *T. timopheevii s.str*: Interestingly, two *T. timopheevii* accessions maintained in two *ex situ* genebanks are reported to originate from Turkey (Supplementary Table S2) [https://www.genesys-pgr.org/]: (i) ATRI 3433 (TRI 3433) conserved in the Federal *ex situ* Genebank of Germany hosted at the Institute of Plant Genetics and Crop Plant Research, IPK, Gatersleben. This *T. timopheevii* line was most likely collected by E. Baur in Turkey in 1926 (Schiemann 1934); and (ii) PI 119442 identified among a barley sample obtained from a market in Araç, near Kastamonu, Turkey (Fig. 3) in 1936 by Westover and Wellmann, and maintained at the National Plant Germplasm System, USDA-ARS, USA. Both accessions harbor the ‘normal’ karyotype of *T. timopheevii*, both were characterized in our studies and confirmed as a typical *T. timopheevii*. Additionally, the accession TA1900, presumably collected 32 km south of Denizli near Kahramanmaraş in Turkey on the 14th of August 1959 and maintained in the wheat germplasm collection of the Wheat Genetics Resource Center, Kansas State University, U.S.A., is interesting because it shared karyotypic features of *T. timopheevii* and the ARA-0 lineage (Fig. 4; Supplementary Figure S19, *i31*). However, we are not fully convinced that this line is a true natural hybrid. Based on the passport data, this accession could potentially have escaped from an experimental field or a breeding station, or received introgression(s) during *ex situ* maintenance (Zencirci et al. 2018). Assuming that the passport data is correct, we could speculate that *T. timopheevii* may have been cultivated in Turkey during the first half of the 20th century. However, is this realistic option?

We believe not. As reported by Stoletova (1924-25), Dekaprelevich and Menabde (1929, 1932), Menabde (1948), Dekaprelevich (1954), *T. timopheevii s.str.* was part of the spring landrace Zanduri (mixture of *T. timopheevii s.str.* and *T. monococcum*) and well adapted to the historical provinces Lechkhumi and Racha of Georgia (Supplementary Material S28). The Zanduri landrace was cultivated in the ‘*humid and moderately cool climate zone 400–800*’ m above the sea level (Dorofeev et al. 1979). Martynov et al. (2018) reported that *T. timopheevii* potentially has a ‘*low potential for plasticity*’ and is not drought tolerant. Climate at origin based on bioclimatic variables (Fick et al. 2017; R Core Team 2017) clearly differs between the regions of Western Georgia where the Zanduri landrace grew till the recent past, and both Kastamonu and Kahramanmara regions in Turkey (Supplementary Material S28). We speculate that the three *T. timopheevii* accessions which were collected in Turkey were probably introduced from Transcaucasia or elsewhere and may have been left over from unsuccessful cultivation or breeding experiments of *T. timopheevii s.str.* in the recent historical past. Also, based on botanical records, *T. timopheevii (s.str.)* has only been identified in Georgia, but not in Turkey or elsewhere (Davis 1965-1988; Hanelt 2001). From this we conclude that the cultivation range of *T. timopheevii* (*s str.*) was not wider in the recent past.

## Conclusions

The evolutionary history of the GGA^t^A^t^ wheat genepool is complex. However, some pieces of the puzzle are clearly recognizable: the region around Dahuk in northern Iraq can be considered the center of origin, but also the center of diversity of *T. araraticum*.

The origin of *T. timopheevii s.str.* remains unclear, but we speculate that it was probably introduced from Turkey, on the grounds that wild *T. araraticum* does not grow in Georgia and that *T. timopheevii s.str.* is more closely related to *T. araraticum* from Turkey or northern Iraq than to the Transcaucasian types.

Based on bioclimatic variables, we predict that *T. timopheevii s.str.* is maladapted to the climate outside Western Georgia. If this speculation is correct, it suggests a sister-group relationship between (i) the Georgian *T. timopheevii* (*s.str.*) and both *T. araraticum* lineages (ARA-0, ARA-1), but also between (ii) *T. timopheevii s.str.* and the prehistoric SW Asian *T. timopheevii s.l*.

The oldest known records of prehistoric *T. timopheevii s.l.* are of Turkish origin: (i) ALıklı Höyük in Cappadocia, and (ii) Cafer Höyük (just inside the Fertile Crescent and potentially older than ALıklı Höyük but less thoroughly dated) (Cauvin et al. 2011; Quade et al. 2018), though more prehistoric finds may be identified in the future. Emphasis for further archaeobotanical research should be given to the central part of the Fertile Crescent, including northern Iraq because this is probably the region where wild *T. araraticum* originated.

The unexpected discovery of the ARA-1 lineage is exciting and may have implications for our understanding of the origins of agriculture in southwest Asia. Was ARA-1 involved in the formation of *T. timopheevii s.l.* (= NGW)?

The distribution area of the ARA-1 lineage requires our attention. It is interesting to note that the ARA-1 lineage (GGA^t^A^t^) and *T. dicoccoides* (BBAA) grow in sympatry and in mixed populations only in a peculiar geographical area in the Northern Levant. This specific area is located between Kahramanmara in the north, Gaziantep in the east, Aleppo in the south and the eastern foothills of the Nur Dağlari mountains in the west. It is interesting that the closest extant wild relatives of domesticated einkorn, barley and emmer were also collected here: (i) the beta race of wild einkorn (Kilian et al. 2007) and the closest wild relatives to einkorn *btr1* type (Pourkheirandish et al. 2018); (ii) the closest wild barley to *btr2* barley (Pourkheirandish et al. 2015) and to *btr1b* barley (Civáň and Brown 2017). This specific region was among the areas predicted with high probability as potential refugia for wild barley during the Last Glacial Maximum (LGM) by Jakob et al. (2014); and (iii) wild emmer subgroup II of the central-eastern race (Özkan et al. 2011). On the other hand, some genetic research has suggested that domesticated emmer wheat and barley received substantial genetic input from other regions of the Fertile Crescent, resulting in hybridized populations of different wild lineages indicating a mosaic of ancestry from multiple source populations (Civáň et al. 2013; Oliveira et al. 2020; Poets et al. 2015). It will be interesting to see whether further genetic and archaeobotanical research on *T. araraticum* lineages and *T. timopheevii s.l.* can help to resolve this issue.

Finally, and fortunately, 1294 accessions of *T. araraticum* and 590 accessions of *T. timopheevii s.str.* are stored in 24 and 37 *ex situ* genebank repositories, respectively (Knüpffer 2009). Our study provides the basis for a more efficient use of *T. araraticum* and *T. timopheevii* materials for crop improvement.

## Acknowledgements

We thank the following for providing seeds, DNA and passport data: N. I. Vavilov Institute of Plant Genetic Resources (VIR, Russia), the Federal *ex situ* Genebank of Germany, Gatersleben (IPK, Germany), the International Center for Agricultural Research in the Dry Areas (ICARDA), the Max Planck Institute for Plant Breeding Research (MPIPZ, Germany), Kyoto University (National Bioresource Project, NBRP, Japan), the John Innes Centre (JIC, UK), Wheat Genetics Resource Centre, Kansas State University (WGRC, USA), Centre for Genetic Resources (WUR, CGN, Netherlands), the Australian Winter Cereal Collection Tamworth (AWCC Australia), the National Plant Germplasm System (USDA-ARS, USA), Eastern Cereal and Oilseed Research Centre, Agriculture and Agri-Food Canada (Canada), Botanical Institute Tbilisi (Georgia), Tbilisi Agricultural Institute (Georgia), Institute of Cytology and Genetics (Siberian Branch of the Russian Academy of Sciences, Russia), and the Russian State Agrarian University Moscow, Timiryazev Agricultural Academy (Russia). We also thank Drs. P.A. Gandilyan, M. Nazarova and I.G. Gukasyan (all Botanical Institute, Erevan, Armenia), Drs. I.D.O. Mustafaev and N.Kh. Aminov (both Institute of Genetic Resources of Azerbaijan, Baku, Azerbaijan) for providing materials from their collection missions. We thank the following for excellent technical assistance: Sigi Effgen (MPIPZ), Christiane Kehler, Birgit Dubsky, Heike Harms, Ute Krajewski, Marita Nix, Kerstin Wolf, Jürgen Marlow, Michael Grau, Peter Schreiber (all IPK), the ‘Experimental Fields and Nurseries’ and ‘Genome Diversity’ groups at IPK. We thank Dr. E.V. Zuev for providing us the data on collection sites of *T. araraticum* and *T. timopheevii* accessions from VIR collection. We are greatly indebted to Maarten Koornneef and George Willcox for discussions and support. We thank two anonymous reviewers, who helped us to improve an earlier version of this manuscript.

## Electronic supplementary material

The online version of this article (https://doi.org/xxx) contains supplementary material, which is available to authorized users.

## Supplementary information

**Supplementary information** accompanies this paper at http xxx

## Declarations

### Availability of data and material

All data sets supporting the conclusions of this article are available in the Electronic supplementary material and from the corresponding author Ekaterina D. Badaeva. The acquisition of plant material used in this study complies with institutional, national, and international guidelines.

### Code availability

Not applicable

### Authors’ contributions

**EB** conceived, designed and conducted the cytogenetic experiments and analyzed data, wrote the first manuscript version.

**FK** conducted the SSAP experiments and analyzed the data.

**HK** verified genebank information concerning the origin of the material, translated texts from Russian into English, and contributed to the discussions and editing of the manuscript.

**AF**, supported the molecular data analysis.

**AR** contributed to the FISH analysis.

**ZK** conducted the simulation of regional agro-ecological variation based on bioclimatic variables.

**SZ** contributed to the work on mapping of Spelt-1 and Spelt-52 probes on *T. timopheevii* and *T. araraticum* genotypes.

**SS** designed and synthesized oligo-probes for FISH analyses.

**KN** contributed to data analysis, supported the field trials and edited the manuscript.

**AG** contributed to the discussions and editing of the manuscript.

**AF** phenotyped and taxonomically re-identified the collection, contributed to the discussions and edited the manuscript.

**KH** phenotyped and taxonomically re-identified the collection, contributed to the discussions and edited the manuscript.

**AB**contributed to the discussions and editing of the manuscript.

**GJ** contributed to the discussions and editing of the manuscript.

**HÖ** conducted the AFLP experiments, analyzed the data, contributed to the discussions and edited the manuscript.

**BK** conceived the project, established the germplasm collection, analyzed data, interpreted results, wrote and edited the manuscript.

### Ethics approval

The authors declare that the experiments comply with the current laws of Germany.

### Consent to participate

Not applicable

### Consent for publication

Not applicable

### Open Access

This article is licensed under a Creative Commons Attribution 4.0 International License, which permits use, sharing, adaptation, distribution and reproduction in any medium or format, as long as you give appropriate credit to the original author(s) and the source, provide a link to the Creative Commons licence, and indicate if changes were made. The images or other third party material in this article are included in the article’s Creative Commons licence, unless indicated otherwise in a credit line to the material. If material is not included in the article’s Creative Commons licence and your intended use is not permitted by statutory regulation or exceeds the permitted use, you will need to obtain permission directly from the copyright holder. To view a copy of this licence, visit http://creativecommons.org/licenses/by/4.0/.

