## Supplementary material for "Genetic diversity, distribution and domestication history of the neglected GGA^t^A^t^ genepool of wheat": Supplementary information.docx

**Supplementary information** accompanies this paper at http xxx

**Supplementary Material S1.docx:** Regarding the first finds of *T. timopheevii;*

**Supplementary Table S2.xlsx: List of materials investigated.** *Column 1: Species (according to GRIS identifiers); taxonomically misclassified accessions are highlighted in red, accessions that do not correspond to their original accession numbers are highlighted in dark blue; Column 2: Subspecies of T. araraticumb(when information is available); Column 3: Accession number; Column 4: Seed/ DNA sources; Column 5: Other identifiers; Column 6: SSAP code; Column 7: SSAP-NNet group; Column 8: AFLP-code; Column 9: AFLP-NNet group; Column 10: group of GGAtAt wheat defined visually based on C-banding-analysis; Column 11: Translocation type - codes of chromosomal rearrangements identified in T. araraticum; Column 12: PASSP - “chromosome passport” codes; Column 13: FISH - based on combination of probes used in FISH analysis; Column 14: Accession number considered by Kawahara et al. (1996): translocation type according to Kawahara et al. (1996); Column 15: Haplotypes of T. araraticum/ T. dicoccoides accessions analyzed by Mori et al. (2009); Column 16: Accessions considered by Bernhardt et al. (2017); Column 17: Haplotypes of accessions according to Gornicki (2014); Column 18: Accession number considered by Nave (2021); Country of origin; Column 19: Collection site.*

**Supplementary Material S3.docx:** I. Details about the SSAP protocol and analysis; II. Details about the AFLP protocol and analysis; III. Details about the C-banding and FISH protocols and analyses;

**Supplementary Figure S4.tif: NeighborNet phylogenetic networks constructed for BBAA- and GGA^t^A^t^-genome wheat accessions based on Dice genetic distances and SSAP markers.** **(a)** combined set of 656 SSAP markers (*BARE-1* and *Jeli* retrotransposon insertions); **(b)** based on 401 markers of *BARE-1* retrotransposon insertions only; **(c)** based on 255 markers of *Jeli* retrotransposon insertions only. According to our previous data (Konovalov et al. 2010) *Jeli* represents mostly the A genome, while *BARE-1* is distributed between A- and B/G-genome chromosomes.

**Supplementary Figure S5.tif: NeighborNet phylogenetic networks constructed for GGA^t^A^t^-genome wheat accessions only based on Dice genetic distances and SSAP markers**. **(a)** combined set of 656 SSAP markers (*BARE-1* and *Jeli* retrotransposon insertions); **(b)** based on 401 markers of *BARE-1* retrotransposon insertions only; **(c)** based on 255 markers of *Jeli* retrotransposon insertions only. Note: the difference in distances between TIM, ARA-0 and ARA-1 groups revealed by different retrotransposon types is possibly reflecting different patterns of A^t^ and G genome diversity generated during evolution. According to our previous data (Konovalov et al. 2010) *Jeli* represents mostly the A genome, while *BARE-1* is distributed between A- and B/G-genome chromosomes.

**Supplementary Table S6.docx:** *a- Summary statistics of genetic variation between four tetraploid wheat groups based on 401 BARE-1, 255 Jeli and the 656 markers combined; b -* *Nei's genetic distance between four tetraploid wheat groups based on 401 BARE-1, 255 Jeli and the 656 markers combined;*

**Supplementary Figure S7.png: Phylogenetic graphs constructed based on 146 polymorphic AFLP markers.** **(a)** Neighbor-Joining (NJ) tree for 103 genotypes based on Jaccard distances [Jaccard 1908, Perrier et al. 2003]; **(b)** NeighborNet planar graph based on Hamming distances for 59 genotypes. The following splits are highlighted: red split separating TIM; blue split separating DIC; green split separating ARA-0; creamy split separating all ARA-1; purple split – separating lines 2677 (IG 117891 from Syria; ARA-1) and 2707 (TRI 17419 from Iraq; ARA-0); light blue split – separating all TIM and four ARA-0 lines from others. **(c)** NeighborNet planar graph based on Hamming distances for 52 genotypes. The following splits are highlighted: red split – separating ARA-1; blue split – separating TIM; green split – separating ARA-0; purple split – separating lines 2677 (IG 117891 from Syria; ARA-1) and 2707 (TRI 17419 from Iraq; ARA-0); brown split – separating all TIM and four ARA-0 lines collected in Armenia and Azerbaijan; light blue split – separating all TIM, five ARA-1 and four ARA-0 lines from all other ARA-1 and all other ARA-0 lines; creamy split – separating TIM and all ARA-0 lines but line 2677 (IG 117891 from Syria; ARA-1) from all lines. See Supplementary Table S2 for more information.

**Supplementary Table S8.docx:** *a - Summary statistics of genetic variation between 103 Triticum genotypes based on 146 polymorphic AFLP markers; b - Nei's genetic distance between 103 Triticum genotypes based on 146 polymorphic AFLP markers;*

**Supplementary Figure S9.docx:** *Neighbor-Joining (NJ) tree for 265 genotypes of T. araraticum and T. timopheevii;*

**Supplementary Table S10.docx:** *a - Summary statistics of genetic variation between 265 genotypes based on 96 polymorphic C-bands considered for chromosomal passports construction.; b - Nei's genetic distance between 265 genotypes based on 96 polymorphic C-bands considered for chromosomal passports construction;*

**Supplementary Table S11.docx:** *a - Summary statistics of genetic variation between 95 genotypes based on polymorphic Spelt-1 and Spelt-52 markers; b - Nei's genetic distance between 95 genotypes based on polymorphic Spelt-1 and Spelt-52 markers;*

**Supplementary Figure S12.tif: The ratios of chromosomal rearrangements among geographic populations of *T. araraticum****.* The circle size is proportional to the number of accessions in respective region.

**Supplementary Table S13.docx:** *Variants of chromosomal rearrangements in T. araraticum;*

**Supplementary Figure S14.tif: Polymorphism of the C-banding patterns of *T. araraticum* accessions from Transcaucasia.** Armenia: *a1 –* KU-1901; *a2 –* TRI 11345; *a3 –* PI 427301; *a4 –* TRI 11358; *a5 –* TRI 11564*a; a6* – K-61654-n; *a7 –* TRI 11564*c; a8* – K-61656 (2G:7G); *a9* – K-61657*b; a10 –* TRI 16599; *a11* – K-59940a; *a12 –* TRI 11564*b; a13 –* TRI 11945*a; a14* – K-61654-T; *a15 –* TA 2893; *a16* – K-59940b. Azerbaijan, Nakhchivan: *b1 –* TRI 7389*b; b2 –* TRI 7389*a; b3 –* TRI 16598; *b4 –* TRI 6912; *b5 –* TRI 7388; *b6 –* TRI 11362; *b7 –* PI 418579; *b8 –* TRI 16597; *b9 –* TRI 11356. Azerbaijan: *b10 –* i-082532; *b11 –* PI 427302; *b12 –* TRI 16602; *b13* – K-61658*a*; *b14* – K-61658*b*. 1A^t^ – 7G – chromosomes; rearranged chromosomes are indicated.

**Supplementary Figure S15.tif: Polymorphism of the C-banding patterns of *T. araraticum* accessions from Iraq (I)**. Dahuk: *c1 –* KU-8877 (13.4 km W from Amadiyah to Bamarni); *c2 –* KU-8884 (21.9 km W from Amadiyah to Dahuk); *c3 –* KU-8907 (22.4 km W from Amadiyah to Bamarni); *c4 –* KU-8822; *c5 –* KU-8824A (15.3 km ENE from Dahuk to Amadiyah); *c6 –* MG 29410 (origin unknown); *c7* – PI 427417; *c8 –* PI 538518; *c9 –* PI 427351; *c10 –* PI 427353 (3–4 km E of Suara-Tuka, Mazorka Gorge); *c11 –* KU-8829; *c12 –* TA 972 (4.4 km NW from Amadiyah); *c13 –* PI 427347; *c14 –* PI 538493 (5.5 km N of Dahuk); *c15 –* PI 427329 (6 km E of Suara Tuka); *c16 –* PI 427420; *c17 –* PI 427425; *c18 –* PI 538498; *c19 –* PI 538499; *c20 –* PI 538500; *c21 –* TRI 11508 (3–4 km E of Suara-Tuka, Mazorka Gorge); Erbil: *d1 –* PI 427311; *d2 –* PI 503292; *d3 –* PI 427380; *d4 –* PI 427381; *d5 –* PI 538470; *d6 –* PI 427376; *d7 –* PI 538461 (1 km NE of Salahaddin); *d8 –* TRI 11507; *d9 –* PI 427386 (2 km NE of Salahaddin). 1A^t^ – 7G – chromosomes; rearranged chromosomes are indicated.

**Supplementary Figure S16.tif: Polymorphism of the C-banding patterns of *T. araraticum* accessions from Erbil, Iraq (II)**. *e1 –* PI 427346; *e2 –* PI 538445 (13 km W of Shaqlawa); *e3 –* KU-8713 (19.1 W of Shaqlawa, SW slope of Pirman Dagh); *e4 –* KU-8715 (17.9 km W from Shaqlawa to Arbil, NE slope of Pirman Dagh) ; *e5 –* KU-8739 from collection of H. Özkan and *e6 –* from the gene bank of Kyoto University, *e7 –* KU-8763; *e8 –* KU-8758; *e9 –* KU-8754 (all from the site “2 km S from the position of 38.9 km from Rowanduz to Rayat”); *e10 –* PI 427406; *e11 –* 538490; *e12 –* PI 427343; *e13 –* PI 538492 (21 km S of Harir); *e14 –* KU-8774; *e15 –* PI 427392; *e16 –* PI 427398; *e17 –* PI 427390; *e18 –* PI 427327; *e19* – PI 427329; *e20* – PI 427334*b; e21 –* PI 427334*a* (4 km NE of Shaqlawa); *e22 –* PI 427399; *e23 –* PI 427402*; e24 –* PI 427403; *e25 –* PI 427341; *e26 –* PI 427339 (7 km NE of Shaqlawa); *e27 –* PI 352265; *e28 –* TA 934; *e29 –* TRI 17418; *e30 –* TRI 16600 (origin is unknown). 1A^t^ – 7G – chromosomes; rearranged chromosomes are indicated.

**Supplementary Figure S17.tif***: Polymorphism of the C-banding patterns of T. araraticum accessions from Sulaimaniyah, Iraq (III).* **Polymorphism of the C-banding patterns of *T. araraticum* accessions from Sulaimaniyah, Iraq (III)**. *f1 –* KU-8451; *f2* – KU-8452; *f3 –* PI 538514; *f4 –* TA 33 (13.2 km S of Sulaimaniyah; *f5 –* KU-8561; *f6 –* KU-8567; *f7* – PI 538513 (14 km S of Sulaimaniyah); *f8 –* PI 427361; *f9 –* PI 427360 (16 km E of Sulaimaniyah); *f10 –* PI 538511 (19 km E of Sulaimaniyah); *f11 –* PI 427355; *f12 –* PI 538505; *f13 –* PI 538504; *f14 –* PI 427435 (41 km NW of Sulaimaniyah); *f15 –* PI 538506; *f16 –* PI 427357*a*; *f17 –* PI 536508; *f18* – PI 427359 (43 km NW of Sulaimaniyah); *f19 –* PI 538456 (44 km NE of Sulaimaniyah); *f20 –* KU-8680; *f21 –* KU-8682; *f22 –* KU-8686 (53 km NW of Sulaimaniyah); *f23 –* KU-8640; *f24 –* KU-8601; *f25 –* KU-8602; *f26 –* KU-8619 (52.4 km SW of Sulaimaniyah); *f27* – KU-8672 (58.3 km SW from Sulaymaniyah to Surdash); *f28 –* KU-8705; *f29* – KU-8706; *f30 –* KU-8710; *f31 –* KU-8709 (Ranja). 1A^t^ – 7G – chromosomes; rearranged chromosomes are indicated.

**Supplementary Figure S18.tif: Polymorphism of the C-banding patterns of *T. araraticum* accessions from Iran** **(*g1–g18*) and Syria** **(*h1–h9*).** Accession codes: *g1 –* IG 113296; *g2* – IG 113297; *g3 –* IG 113298; *g4 –* 113295; *g5–* IG 113294 (from Sar Dasht); *g6 –* KU-1946; *g7 –* KU-8944; *g8 –* KU-8945 (12.2 km NW from Karand to Qasri Shirin); *g9 –* Amyn (unknown); *g10 –* PI 427366; *g11 –* TRI 11509; *g12 –* CItr 17680; *g13 –* PI 427369; *g14 –* PI 427304b; *g15 –* TA 05; *g16 –* PI 427365 (47 km NW of Eslamabad (Shahabad) to Sar-e-Pol-e-Zahab); *g17 –* KU-8947; *g18 –* KU-8948 (15.1 km NW from Karand to Qasri Shirin). Accessions from Syria: *h1–h3 –* three lines of cytogenetically uniform accession IG 119456; *h4* and *h5 –* IG 117891; *h6 –* IG 117895N; *h7 –* IG 117895c; *h8 –* IG 117895b; *h9 –* IG 117895a. A^t^ – G – genomes; 1–7 – homoeologous groups; rearranged chromosomes are indicated.

**Supplementary Figure S19.tif: Polymorphism of the C-banding patterns of *T. araraticum* from Turkey** (**ARA-0**). Kahramanmaraş *(i1, i2),* Diyarbakır (*i3–i15*), Batman (*i16*), Elazığ (*i17–i18*), Tunceli (*i19, i20*), Mardin (*i21–i25*), Siirt (*i26–i30*) and Denizli (*i31*): *i1 –* KU-1986; *i2 –* KU-1943 (45 km SE of Maras); *i3 –* KU-1923; *i4 –* KU-1930; *i5 –* KU-1935; *i6 –* KU-1936; *i7 –* KU-1925; *i8 –* KU-1939; *i9 –* KU-1938 (12 km E of Silvan); *i10 –* KU-8917; *i11 –* KU-1923; *i12 –* KU-8918; *i13 –* KU-8924; *i14 –* KU-8925 (17.3 km E from Silvan to Bitlis); *i15 –* IG 46426 (18 km W Kurtalan); *i16 –* IG-46427 (Batman, 5 km E Kozluk junction); *i17 –* TRI 17028 (Elazığ); *i18 –* KU-8938 (39.9 km N from Elazığ to Hozat); *i19 –* CItr 17677; *i20 –* TA 976 (17 km S of Tunceli on road from Elazığ to Erzincan); *i21 –* KU-8908; *i22 –* KU-8910; *i23 –* KU-8912; *i24 –* KU-8913; *i25 –* KU-8909 (26.3 km NE from Mardin to Midyat); *i26 –* PI 560872; *i27 –* PI 560697; *i28 –* PI 560877; *i29 –* PI 560873; *i30 –* PI 596290 (Siirt); *i31 –* TA1900 (32 km S of Denizli in the Taurus Mountain Range). 1A^t^ – 7G – chromosomes; rearranged chromosomes are indicated.

**Supplementary Figure S20.tif: Polymorphism of the C-banding patterns of *T. araraticum* from Turkey** (**ARA-1**). Kahramanmaraş: *j1 –* KU-1950; *j2 –* KU-1960; *j3 –* KU-1962b; *j4 –* KU-1964; *j5 –* KU-1965; *j6 –* KU-1984B; *j7 –* KU-1982; *j8 –* TA 1008; *j9 –* PI 658873; *j10 –* 3127; *j11 –* 3128; *j12* – *3130; j13* – TA 1475; Gaziantep: *k1 –* IG 116164; *k2 –* IG 116165; *k3 –* PI 564340; *k4 –* IG 116168; *k5 –* IG 116176; *k6 –* IG 116177; *k7 –* IG 46434; *k8 –* IG 116166; *k9 –* PI 658869b; Kilis: *l1 –* IG 116169; *l2 –* 2631; *l3 –* 2633; *l4 –* 2634; *l5 –* 2639; *l6 –* 2641; *l7 –* 2644; *l8 –* 2643; *l9 –* 3030. 1A^t^ – 7G – chromosomes; rearranged chromosomes are indicated.

**Supplementary Material S21.docx**: *Region-specific chromosomal rearrangements;*

**Supplementary Figure S22.tif: Comparison of the distribution of pAesp_SAT86 probe on chromosomes of *T. dicoccoides* (DIC), *T. timopheevii* TIM (T1–T4), *T.* araraticum ARA-1 (A1–A7) and ARA-0 (A8–A24).** DIC – KU-8836A (Iraq, SSW of Rowanduz); T1 *–* K-38555; T2 – K-29558; T3 – Budashkina; T4 *–* KU-107; A1 *–* IG 116164; A2 *–* IG 116169; A3 – KU-1982; A4 *–* IG-1984B; A5 *–* TA 1008; A6 *–* IG 119456; A7 – 2634; A8 *–* KU-1986; A9 – KU-8926; A10 *–* PI 427398; A11 – PI 538516; A12 *–* TRI 11507; A13 *–* KU-8907; A14 – KU-8824A; A15 – PI 427417; A16 – KU-8595; A17 – KU-8602; A18 – KU 8619; A19 – CItr 16780; A20 – IG 113295; A21 – K-59940; A22 – K-61658; A23 – K-61654; A24 – 58667. Countries of origin of *T. araraticum* accessions are shown above the accession codes. Rearranged chromosomes are designated and indicated with arrows; red lines show position of pAesp_SAT86 sites typical for ARA-1 or for ARA-1 + TIM groups; green lines indicate position of pAesp_SAT86 sites typical for ARA-0 group; yellow lines – pAesp_SAT86 sites typical for *T. timopheevii* only.

**Supplementary Figure S23.tif: Distribution of Spelt-1 (red) and Spelt-52 (green) probes on chromosomes of *T. timopheevii* (T) and *T. araraticum* representing ARA-1 (A1, A2, A4–A10) and ARA-0 (A3, A11–A29) groups**. Accession numbers: T – KU-1819; A1 – IG 117891; A2 – IG 117895C; A3 – IG 117895A; A4 – IG 119456 (Syria); A5 – IG 116165; A6 – IG 116170; A7 – KU-1984B; A8 – #2633; A9 – 32634; A10 – KU-1860; A11 *–* KU-8910; A12 – KU-1930; A13 – KU-8939 (Turkey); A14 – PI 427346; A15 – KU-8907; A16 – PI 538518; A17 – TRI 17419; A18 – KU-8774; A19 – PI 538516; A20 – PI 547380; A21 – PI 538505; A22 – PI 427364; A23 – KU-8682; A24 – KU-8451 (Iraq); A25 – KU-8944; A26 – CItr 17680; A27 – IG 113294 (Iran); A28 – TRI 11356; A29 – K-59940 (Transcaucasia)*.* 2A^t^, 6A^t^, 1G – 7G chromosomes carrying Spelt-1 and/or Spelt-52 signals arranged according to genetic nomenclature. *T. timopheevii* and ARA-1 genotypes are shown with pink numbers, ARA-0 genotypes with green numbers. Arms of rearranged chromosomes are designated according to their origin.

**Supplementary Table S24.docx: Distribution of Spelt-1 and Spelt-52 loci over chromosomes of 87 *T. araraticum,* seven *T. timopheevii* and one *T. zhukovskyi* genotypes.** Accession number; Number of Spelt-1 signals (per haploid genome); position of Spelt-1 loci on the A^t^ or G-genome chromosomes; position of Spelt-52 loci on the A^t^ or G-genome chromosomes; Chromosomal group. The signal size at the respective chromosomal position is assessed according to signal intensity: 1 – very weak; 2 – weak, 3 – medium, 4 – large, 5 – very large. Empty boxes indicate the absence of signal at the respective position.

**Supplementary Material S25.docx:** *Additional findings of the FISH analysis;*

**Supplementary Table S26.docx: Distribution of Spelt-1 and Spelt-52 loci in geographic/chromosomal groups in comparison with *T. timopheevii.*** Region/ Country of origin; # – number of genotypes per region/ country; frequencies of occurrence of Spelt-1 and Spelt-52 signals in particular positions are given in absolute values. Group – chromosomal group of genotypes in particular region defined by C-banding analysis. The frequencies of Spelt-1 and Spelt-52 loci for total *T. araraticum* (ARA) in comparison with *T. timopheevii* (TIM) groups are given in percentage.

**Supplementary Material S27.docx:** *Information extracted and translated from Dorofeev et al. (1979), regarding the cultivation area and geographical distribution of Triticum timopheevii, and the "Zanduri" wheat complex. Translation by H. Knüpffer and colleagues, February 2020;*

**Supplementary Material SM28.docx:** *Was the cultivation range of T. timopheevii (s.str.) wider in the recent past?*
