## Supplementary material for "Genetic diversity, distribution and domestication history of the neglected GGA^t^A^t^ genepool of wheat": Supplementary_Figure S4_and S5.docx

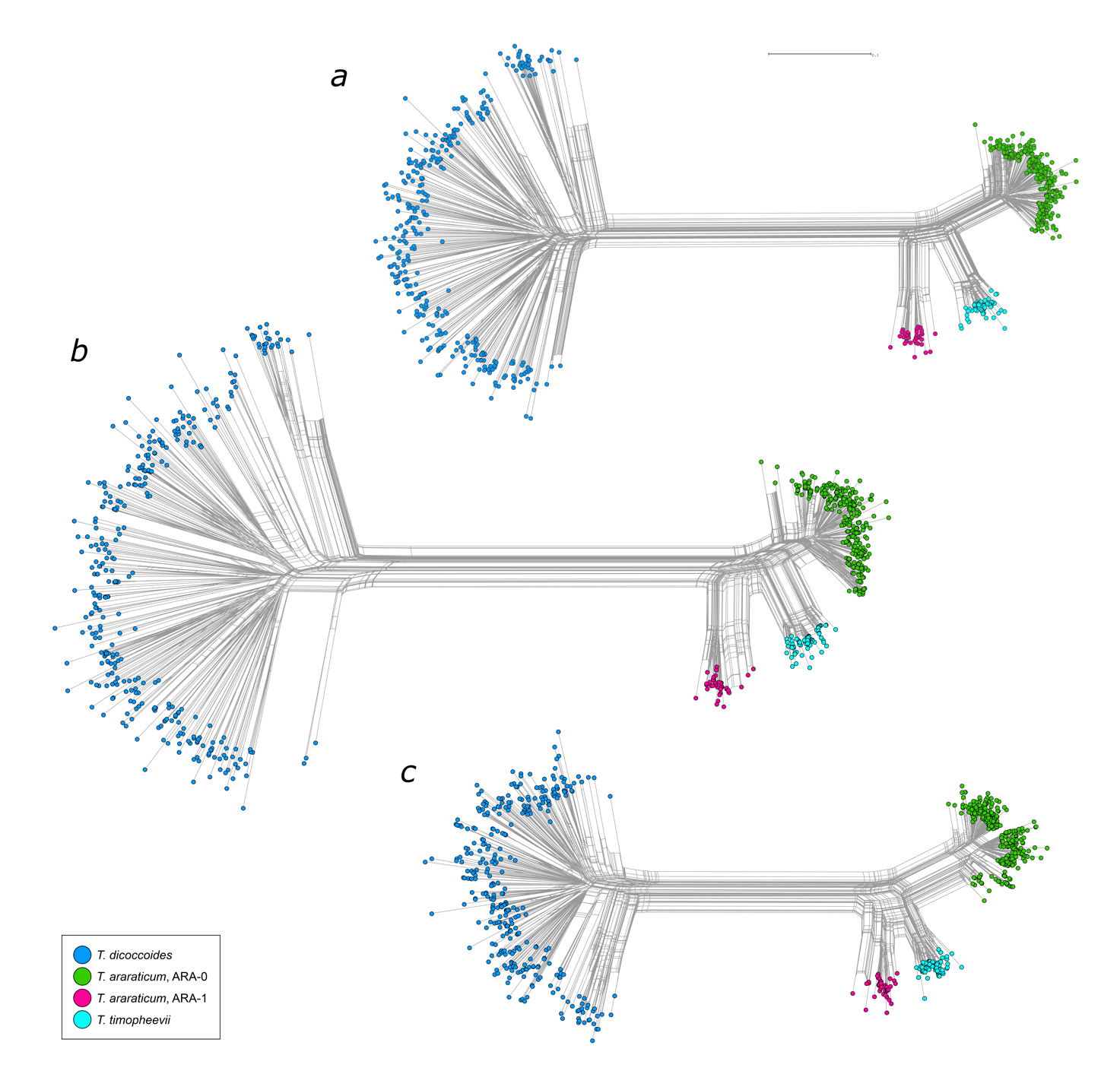

**NeighborNet phylogenetic networks constructed for BBAA- and GGA^t^A^t^-genome wheat accessions based on Dice genetic distances and SSAP markers.** **(a)** combined set of 656 SSAP markers (*BARE-1* and *Jeli* retrotransposon insertions); **(b)** based on 401 markers of *BARE-1* retrotransposon insertions only; **(c)** based on 255 markers of *Jeli* retrotransposon insertions only. According to our previous data (Konovalov et al. 2010) *Jeli* represents mostly the A genome, while *BARE-1* is distributed between A- and B/G-genome chromosomes.

**Supplementary Figure S5**

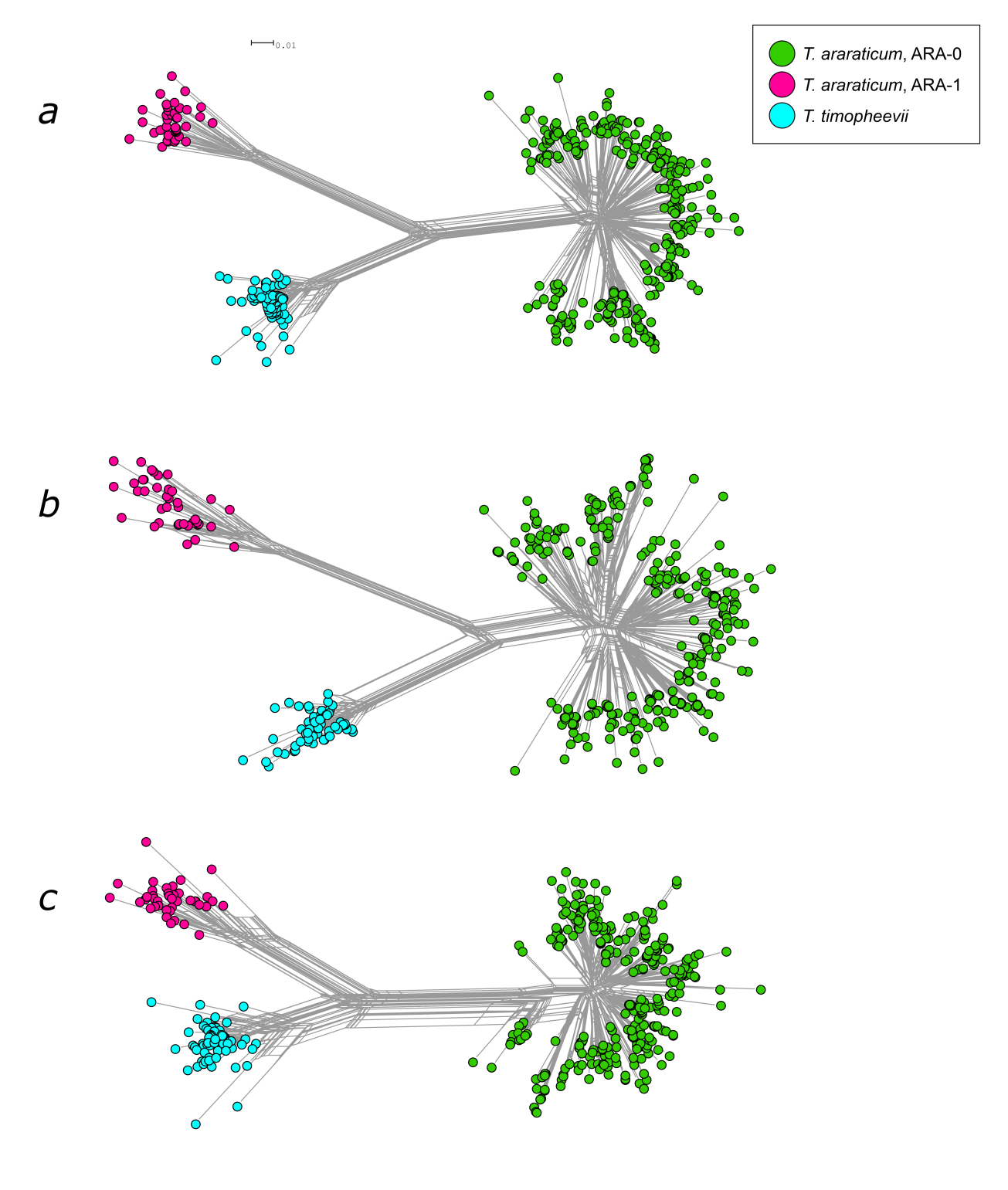

**NeighborNet phylogenetic networks constructed for GGA^t^A^t^-genome wheat accessions only based on Dice genetic distances and SSAP markers**. **(a)** combined set of 656 SSAP markers (*BARE-1* and *Jeli* retrotransposon insertions); **(b)** based on 401 markers of *BARE-1* retrotransposon insertions only; **(c)** based on 255 markers of *Jeli* retrotransposon insertions only. Note: the difference in distances between TIM, ARA-0 and ARA-1 groups revealed by different retrotransposon types is possibly reflecting different patterns of A^t^ and G genome diversity generated during evolution. According to our previous data (Konovalov et al. 2010) *Jeli* represents mostly the A genome, while *BARE-1* is distributed between A- and B/G-genome chromosomes.
