## Supplementary material for "Genetic diversity, distribution and domestication history of the neglected GGA^t^A^t^ genepool of wheat": Supplementary_Figure S7.docx

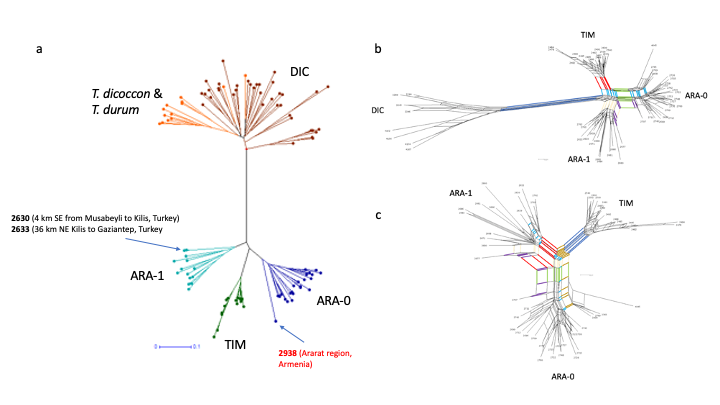

**Phylogenetic graphs constructed based on 146 polymorphic AFLP markers.** **(a)** Neighbor-Joining (NJ) tree for 103 genotypes based on Jaccard distances [Jaccard 1908, Perrier et al. 2003]; **(b)** NeighborNet planar graph based on Hamming distances for 59 genotypes. The following splits are highlighted: red split separating TIM; blue split separating DIC; green split separating ARA-0; creamy split separating all ARA-1; purple split – separating lines 2677 (IG 117891 from Syria; ARA-1) and 2707 (TRI 17419 from Iraq; ARA-0); light blue split – separating all TIM and four ARA-0 lines from others. **(c)** NeighborNet planar graph based on Hamming distances for 52 genotypes. The following splits are highlighted: red split – separating ARA-1; blue split – separating TIM; green split – separating ARA-0; purple split – separating lines 2677 (IG 117891 from Syria; ARA-1) and 2707 (TRI 17419 from Iraq; ARA-0); brown split – separating all TIM and four ARA-0 lines collected in Armenia and Azerbaijan; light blue split – separating all TIM, five ARA-1 and four ARA-0 lines from all other ARA-1 and all other ARA-0 lines; creamy split – separating TIM and all ARA-0 lines but line 2677 (IG 117891 from Syria; ARA-1) from all lines. See Supplementary Table S2 for more information.
