## Supplementary material for "Genetic diversity, distribution and domestication history of the neglected GGA^t^A^t^ genepool of wheat": Supplementary_Figure S9.docx

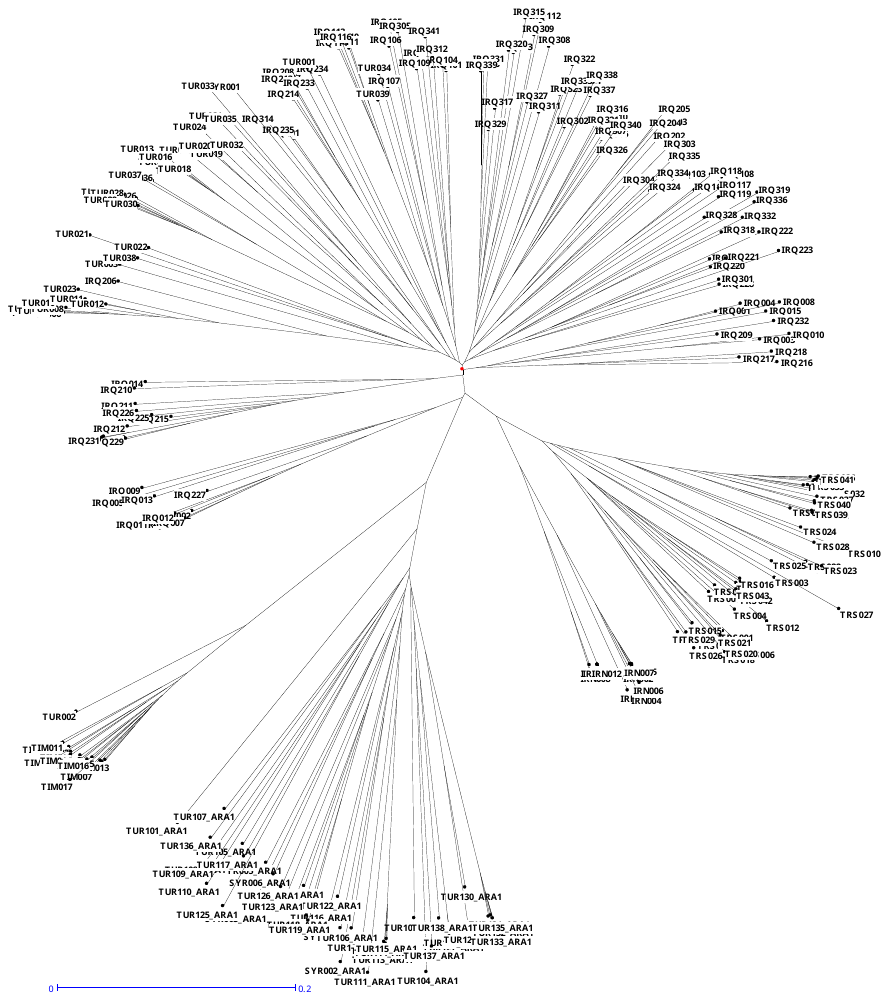

ARA-1

ARA-0

TIM

TUR002: TA1900

**Neighbor-Joining (NJ) tree for 265 genotypes** of *T. araraticum* and *T. timopheevii* using 379 C-banding markers based on Jaccard distances (Jaccard 1908; Perrier et al. 2003).
