## Supplementary material for "Genetic diversity, distribution and domestication history of the neglected GGA^t^A^t^ genepool of wheat": Supplementary_Material_S1.docx

**Regarding the first finds of *T. timopheevii***

The original *T. timopheevii* accession that was used as standard in species description was obtained by P.M. Zhukovsky from Shida Kartli (Inner Kartli, formerly Gori) region of Eastern Georgia (part adjacent to Likhi Range. However, the expedition of Dekaprelevich and Menabde (1932) failed to find *T. timopheevii* in this region.

Zhukovsky described *T. timopheevii* as a wild or weedy plant in his 1923 and 1928 papers (Zhukovsky 1923, 1928), but Stoletova (1924) noted that the rachis was not as brittle as in typical wild two-grained wheats and considered the plants found by Zhukovsky to be feral cultivated plants. She and others found *T. timopheevii* under cultivation in a restricted area of western Georgia (eastern part), mostly in the regions of Lechkhumi and Racha (Stoletova 1924–1925; Dekaprelevich and Menabde 1929, 1932; Menabde 1948, Dekaprelevich 1954). Zhukovsky's find was therefore just outside the range of twentieth century *T. timopheevii* cultivation.
