## Supplementary material for "Genetic diversity, distribution and domestication history of the neglected GGA^t^A^t^ genepool of wheat": Supplementary_Material_S3.docx

**I. Details about the SSAP protocol and analysis**

At the first step, 125 ng DNA was digested with *Taq*I restriction endonuclease (NEB) for 3h at 65°C in 40 μl volume containing 1x NEBuffer 4, 100 μg/ml BSA and 10U *Taq*I. Subsequent adapter ligation was performed in the same tubes or plates by adding 10 μl ligation mixtures for obtaining final concentrations of 1x NEBuffer 4, 100 μg/ml BSA, 50 pmol *Taq*I adaptor, 1 mM ATP and 3U T4 DNA Ligase (Invitrogen). The mix was incubated at 37°C for 6h followed by 65°C for 1h, and then diluted twice with 50 μl deionized water. Next, two subsequent rounds of PCR were performed. First, 5 μl of ligaton products were amplified in 25 μl mix with final concentrations of 1U HotStarTaq DNA polymerase (Qiagen), 1.5 mM MgCl_2_, 20 μM dNTPs each, 0.2 μM LTR primer and 1x PCR buffer. The PCR program consisted of enzyme activation for 14 min 30 sec at 95°C, 30 cycles of denaturation at 95°C for 30 s, annealing at 62°C for 1 min, an extension at 72°C for 2 min, followed by a final extension of 10 min at 72°C. Next, 15 μl of the first PCR product was amplified in 25 μl mix with final concentrations of 1U *Taq* DNA polymerase (Qiagen), 1.5 mM MgCl_2_, 200 μM dNTPs each, 1 μM LTR primer, 1 μM adapter primer and 1x PCR buffer. The second-round PCR program consisted of 35 cycles of denaturation at 95°C for 30 s, annealing at 62°C for 1 min, an extension at 72°C for 2 min, followed by a final extension of 10 min at 72°C. The LTR primer sequences used were 5'-labeled with TET or HEX. PCR products were separated on MegaBACE 1000 capillary electrophoresis system (GE Healthcare) according to manufacturer’s recommendations. Electropherograms obtained with nine primer combinations (using 3’-selective bases ACG, AG, CAG, GAC, CTG for *Jeli* LTR primers and CAGT, TGAC, AGTC and ACTG for *BARE-1* LTR primers) were scored automatically by MegaBACE proprietary software with complete manual check.

**II. Details about the AFLP protocol and analysis**

Initially, eight genotypes, namely 2484, 2495 (all *T. timopheevii*), 2660, 2683, 2671, 2677 (all *T. araraticum*), and 3313, 3354 (all *T. dicoccoides*, Supplementary Table S2), were screened with 40 AFLP primer combinations in order to select most promising primer combinations for the analysis. Only combinations that yielded a high number of polymorphic fragments between the eight genotypes but also reliably scoreable DNA markers were considered. In total, six AFLP primer combinations (EACA-MAAG, EACA-MACC, EACC-MAAA, EACC-MACG, EAGA-MAGT, EAGC-MACG) were used to screen the collection of 104 lines. Restriction-ligation mixtures were performed in a total volume of 11 μl containing 10X T4 DNA Ligase buffer, 0.5 M NaCl, 1 mg/μl BSA, 5 pmol *EcoR*I adaptor, 50 pmol *Mse*I adaptor, 10 μ/μl *Eco*RI enzyme, 5 μ/ μl T4 DNA Ligase enzyme, and 50–100 ng of DNA. Restriction-ligation mixtures were then incubated at 37°C for 8 h. The pre-selective amplification was performed according to Zabeau and Vos (1993), with modifications according to Altıntaş et al. (2008). Subsequently, the selective amplification was carried out in a final volume of 20 μl containing 10X PCR buffer, 25 mM MgCl_2_, 2.5 mM dNTP, 5 pmol *Eco*RI, 5 pmol *Mse*I, 1 μl *Taq*, and pre-amplified DNA. The PCR program consisted of 20 cycles of denaturation at 94°C for 30 s, annealing at 56°C for 30 s, an extension at 72°C for 2 min, followed by a final extension of 30 min at 60°C. Ten microliters of the AFLP selective amplification product were mixed with 10 μl of loading buffer, denatured at 94 °C for 5 min, and placed on ice. The LI-COR 4300 DNA Analyzer was used for AFLP analysis. After a pre-run of electrophoresis at 40 W for 15 min on LI-COR, 1 μl of the amplified product was loaded onto a 6 % denaturing polyacrylamide gel (19:1) with 0.5× TBE buffer and run at 40 W until the loading dye reached the bottom of the gel. Autoradiographs were scored for the absence (0) or presence (1) of AFLP bands manually and independently at least twice.

**III. Details about the C-banding and FISH protocols and analyses**

*Passportization in the C-banding analysis*

The idiogram (Fig. 1) contained 131 bands distributed among fourteen A^t^ and G-genome chromosomes, however, only 96 informative C-bands were considered for the construction of chromosomal passports. The remaining 35 bands were excluded from the analysis because they were either invariable in size or inconsistent and difficult to score. In contrast to our previous report (Badaeva et al. 2015), the size of C-bands was estimated using a 6-grade scale – from “0” for C- bands which were absent at a particular position, to “5” for very large bands (over 1 µm). In addition to the band size, we considered the pericentromeric inversion often observed on the chromosome 4G. The absence of inversion was encoded as “2”, the presence, as “3”. NeighborNet planar graph between all 265 wheat genotypes was constructed based on the karyotype of each line using SplitsTree 4.15.1 (Huson and Bryant 2006). C-bands were equally weighted and coded for each individual at all loci using the size estimated.

*Fluorescence in situ hybridization (FISH)*

The probes were labeled with Fluorescein and biotin and detected using anti-Fluorescein- Fluorescein/ Oregon green, rabbit IgG fraction, Alexa Fluor 488 conjugate (Molecular Probes, USA) and streptavidin-Cy-3 (Amersham Pharmacia Biotech, USA) according to (Zoshchuk et al. 2007). The slides were counter-stained with DAPI (4′,6-diamidino-2-phenylindole) in Vectashield mounting media (Vector laboratories, Peterborough, UK) and analyzed on a Zeiss Imager D-1 microscope. Selected metaphase cells were captured with an AxioCam HRm digital camera using software AxioVision, version 4.6.3. Images were processed in Adobe Photoshop, version CS6 (Adobe Systems, Edinburgh, UK).
