## Supplementary material for "Genetic diversity, distribution and domestication history of the neglected GGA^t^A^t^ genepool of wheat": Supplementary_Material_S21.docx

**Region-specific chromosomal rearrangements**

*Transcaucasia* (**Supplementary Figure S14**):

Fifteen variants of chromosomal rearrangements were identified in *T. araraticum* from Transcaucasia. Most variants belonged to translocation lineages identified earlier by Badaeva et al. (1990, 1994). The translocation T2A^t^:4G gave rise to three variants: a novel translocation T2A^t^:4G + T6A^t^:7A^t^, T2A^t^:4G + T4A^t^:7A^t^ and T2A^t^:6A^t^:4G (Fig. 6). Hybridization of either one of two latter genotypes with a genotype carrying T1G:2G-2 + T4G:6G resulted in formation of a unique karyotype, which possessed six pairs of translocated chromosomes T2A^t^:6A^t^:4G:6G + T1G:2G-2 (Fig. 4). The second lineage also started from T2A^t^:4G followed by T2A^t^:4G + T4A^t^:7A^t^, as in Armenia, however the subsequent steps were different. A tertiary translocation involved chromosome 7G resulting in a triple translocation T2A^t^:4G:7G + T4A^t^:7A^t^, which significantly dominated in Nakhichevan (21 accessions). This translocation gave rise to a quadruple translocation T2A^t^:4G:6G:7G + T4A^t^:7A^t^. The third, minor translocation lineage started from T1G:2G-2. Hybridization of this genotype with another one, carrying the T3G:4G-2 led to the formation of a novel variant of double translocation T1G:2G-2 + T3G:4G-2. Translocations T1G:5G, T4G:5G, T2G:7G and novel translocation T6A^t^:7G were found in 1–2 accessions each.

*Iraq* **(Supplementary Figures S15-S17**)

Thirty-three local populations from Iraq harbored the highest diversity of chromosomal rearrangements: altogether 34 variants were identified (Fig. 6; Supplementary Table S2; Supplementary Table S13). Translocation variants were usually restricted to one or few neighboring populations, which, in turn, may contain a mix of accessions with normal karyotype and one to several translocation variants. According to the geographical location, populations were divided into three groups: (i) Dahuk-Amadiyah, (ii) Erbil (Shaqlawa), and (iii) Sulaymaniyah.

The Dahuk group contained the lowest rate of accessions with rearranged karyotypes (24.3%) and five translocation variants were assigned (Fig. 6). Among them the translocation T1G:3G was found in four, T3G:4G-2 in two accessions, and translocations T2G:6G-1, T1G:2G-1, T4G:5G-2, and T5A^t^:7G+T4G:5G-3 – in a single accession each (Figs. 4, 6; Supplementary Figure S15; Supplementary Table S2), usually intermixed with karyotypically normal accessions. Some populations from Dahuk did not possess translocated accessions at all.

The Erbil group (Supplementary Figure S15; Supplementary Figure S16) included 65.1% karyotypically rearranged accessions. Chromosomal rearrangements were represented by 17 variants, among them three inversions, 11 single and three double translocations (Fig. 6). Translocations T2G:4G:6G, T6G:7G, T1G:4G-2, were frequent and found in 13, 13, and seven accessions respectively. Translocations T5A^t^:3G, T2G:4G, the paracentric inversion of 2A^t^ and pericentric inversion *per*Inv6G, were identified each in three accessions. Seven other translocations variants were unique (Fig. 6; Supplementary Table S2; Supplementary Table S13). Double translocations could prevail over the original single translocation (T2G:4G:6G >> T2G:6G), could be less frequent (e.g., T5A^t^:3G +T4G:5G-1 << T5A^t^:3G) or the original and derived translocations could be equally rare (T3G:6G-1 + T4G:5G-2 = T4G:5G-2). Some populations included accessions with normal karyotypes in a mix with one to four translocation variants. Other populations contained only accessions with translocated karyotypes belonging to one or several translocation variants.

The Sulaymaniyah group (Supplementary Figure S17) also contained a high ratio (61%) of translocated accessions. Chromosomal rearrangements were represented by 16 variants with frequencies ranging from eight accessions (T5A^t^:1G), four (inv7A^t^-1), three (T4A^t^:4G), two (T3G:7G and its derivative T3G:7G +T6A^t^:6G) to one accession each (10 variants). Some populations possessed only translocated accessions belonging to one or several translocation variants, the other contained predominantly karyotypically normal accessions.

*Iran* (**Supplementary Figure S18 g1–g18)**

Two populations represented the Iranian group. Accessions carrying the pericentric inversion of the 7A^t^ chromosome, either alone (11 accessions) or in combination with different secondary translocations dominated in Iran: three accessions with T1G:5G-2 and three with the novel translocation T6A^t^:6G. In contrast to previous studies (Badaeva et al. 1994, Mitrofanova et al. 2016), five accessions all originating from Sar Dasht (Khuzestan) lacked this inversion. Two of them harbored normal karyotypes and three carried novel translocations: T3A^t^:6A^t^ (two accessions, Table S3) and T7A^t^:2G (IG 113298). Altogether, 91% *T. araraticum* accessions from Iran possessed only rearranged karyotypes.

*Syria* (**Supplementary Figure S18 h1–h9)**

Only one of the six Syrian *T. araraticum* harbored the normal karyotype, and it was assigned to ARA-0. Three types of chromosomal rearrangements were identified among the remaining five accessions, all belonged to ARA-1. Among them, translocations T1G:3G and T2G:5G-1 were also found as in other countries but T3G:6G-2 was unique to Syria.

*Turkey* (**Supplementary Figure S19; Supplementary Figure S20**)

Forty percent of Turkish accessions carried chromosomal rearrangements assigned to 25 variants (Fig. 6). Altogether, we studied 33 populations, represented by one to 15 genotypes. According to geographical coordinates of the collection sites, populations were divided into two major geographical groups, which coincided with the chromosomal groups: (i) west of the Euphrates river (Adiyaman, Gaziantep, Kilis, Kahramanmaraş), and (ii) east of the Euphrates river (Batman, Diyarbakır, Elazığ, Mardin, Siirt, Tunceli) (Fig. 3). Noteworthy, the Eastern group consisted exclusively of ARA-0 accessions, whereas ARA-1 accessions significantly dominated in the Western group (95.5%).

Accessions from the Eastern group (Supplementary Figure S19) carried mostly normal karyotypes, and eight variants of chromosomal rearrangements (Fig. 6; Supplementary Table S13) were identified in nine accessions (25.6%). One variant, T2G:4S (Siirt) was also found in Iraq, whereas other translocations were unique to this region. Interestingly, two of the four A^t^:A^t^ genome translocations identified in *T. araraticum*, namely T2A^t^:6A^t^ and T5A^t^:6A^t^ (Fig. 5; Supplementary Table S13), were detected in Eastern Turkey. Two variants were multiple translocations: T3G:4G:7G and T2A^t^:7G + T6A^t^:5G:7G, however, their preceding single translocations were not found. In most cases accessions with translocated karyotypes appeared in small frequencies among karyotypically normal accessions.

The Western group was represented by 44 accessions, of them 42 belonged to the ARA-1 lineage (Supplementary Figure S20) and only two accessions were assigned to ARA-0 (accessions KU-1943 and KU-1986 from Kahramanmaraş (Supplementary Figure S19). Eighteen variants of chromosomal rearrangements were identified in 24 ARA-1 (54.5%) accessions, among them four (T1G:3G, T4G:7G, T4G:6G-1, and T1G:5G-1) were also found in ARA-0 accessions from other geographical regions (Supplementary Table S13). Most translocations were rare and occurred usually in a mix with karyotypically normal genotypes. The translocation T3A^t^:7A^t^-1 was identified in three accessions collected from Kilis and it gave rise to the double translocation T3A^t^:7A^t^-1 +T4A^t^:7G (Figs. 5, 6).
