## Supplementary material for "Genetic diversity, distribution and domestication history of the neglected GGA^t^A^t^ genepool of wheat": Supplementary_Material_S25.docx

**Additional findings of the FISH analysis**

The number of Spelt-1 sites in *T. araraticum* ranged between two and seven per haploid genome, 4.72 on average (Supplementary Table S26). Over 95% of *T. araraticum* and all *T. timopheevii* genotypes carried Spelt-1 sites on 4GL and 5GL. About 75% of *T. araraticum* and all *T. timopheevii* possessed Spelt-1 site on the satellite of chromosome 6A^t^S, but the signal size was highly polymorphic. Spelt-1 signals on 1GL, 2GL and 6GS were frequent in both *T. araraticum* and *T. timopheevii*. Frequencies of signals in ‘rare’ positions were lower and ranged from 6.9% (7GL, 6GL) to 33.7% (2A^t^L).

The patterns of Spelt-1 and Spelt-52 repeats varied across geographic regions and between chromosomal groups. Genotypes lacking the Spelt-1 locus on 6A^t^S originated mainly from northeastern Iraq (Dahuk, Sulaymaniyah), and Eastern Turkey (two regions). The Spelt-1 locus on 2A^t^L was absent in Transcaucasia and Erbil (Iraq). It was rare in Turkey, present in approximately half of the genotypes from Iran, Sulaymaniyah (Iraq), and Syria, and always present in *T. araraticum* from Dahuk (Iraq) and in *T. timopheevii*. The Spelt-1 signal on 6GS occurred in Transcaucasia, ARA-1 genotypes from Turkey and in about ¼ of *T. timopheevii*. It was rarely found in Iraq and was absent from the Iranian group. The Spelt-1 signal on 2GL varied significantly in size (Supplementary Table S24). Usually, it was present in high frequencies in *T. araraticum* from Erbil, Iraq (62.5%), and less frequently in Iran and Turkey (33.3%). This locus was rare in Transcaucasian ARA-0, while all *T. timopheevii* carried a large Spelt-1 site on 2GL. The medium-to-large signal on 3GL occurred mainly in Iran and Sulaymaniyah (Iraq). The Spelt-1 signal on 6GL was found in only six genotypes, five from Sulaymaniyah (Iraq) and one from Syria. One very large Spelt-1 signal on 7GS was observed in several ARA-0 genotypes from northeastern Iraq and in few ARA-1, while the signal on 7GL was found in the ARA-1 group only (Supplementary Figure S23). Spelt-1 signals at these ‘rare’ positions were not observed in *T. timopheevii.* The ARA-1 lineage was characterized by a higher number of Spelt-1 sites (5.38 in average) compared to ARA-0 (4.52) due to a more frequent occurrence of loci at 6A^t^S, 1GL, 6GS, and 7GL (Supplementary Table S25, Supplementary Table S26), whereas the loci 2A^t^L and 2GL were less frequent. The frequency of Spelt-52 sites in *T. araraticum* varied from 6.9% on 2GS to 94.3% on 2A^t^S. A small Spelt-52 signal on 1GL was present in 43.7% of *T. araraticum* genotypes. It was missing in Transcaucasia and Turkey, occurred rarely in southern and central Iraq, but was rather frequent is Iran, northern and eastern Iraq, and in the ARA-1 group (the frequency was twice higher in ARA-1 than in ARA-0). A large Spelt-52 site on 6GL was detected in 86% of *T. araraticum* genotypes, including all ARA-0 and eight ARA-1. It was absent from 12 ARA-1 genotypes (47.6%) and all *T. timopheevii*. Thus, absence of the Spelt-52 site on 6GL was specific for *T. timopheevii.* Some of the ARA-1 genotypes, instead of a 6GL site acquired a new Spelt-52 site on 2GS (Supplementary Figure S23), and in two ARA-1 genotypes the emergence of this locus was accompanied with a significant decrease of the Spelt-52 signal on 6GL. Thus, a new Spelt-52 site probably emerged due to reciprocal translocation between chromosomes 6GL and 2GS.
