## Supplementary material for "Genetic diversity, distribution and domestication history of the neglected GGA^t^A^t^ genepool of wheat": Supplementary_Material_S27.docx

**Information extracted and translated from Dorofeev et al. (1979) ^[1]^, regarding the cultivation area and geographical distribution of *Triticum timopheevii*, and the "Zanduri" wheat complex.** Translation by H. Knüpffer and colleagues^[2]^, February 2020

Apparently, *T. timopheevii* has never been found in cultivation outside Georgia.

**[Chapter on *T. monococcum*]**

<295> […] *T.* *monococcum* is considered by Palmova (1935) as a dry-steppe ecotype. However, domesticated einkorn originally from the Zanduri complex of western Georgia, such as highland European forms, is undoubtedly a mesophytic type. Menabde (1948) identified the domesticated einkorn of western Georgia as a special ecotype. All forms of domesticated einkorn are highland types but not high-mountain types. As Zhukovsky has explained (1950, 1964), in ancient irrigated agricultural regions of Mesopotamia, it was unable to compete with emmer and barley and, for this reason, became localized in the colder mountain regions where there is an abundance of natural precipitation.

Cultivation of domesticated einkorn in Georgia reached 900–1,600 m, in Chechnya and Ingush 1,500 m, and in Spain 750 m. According to Davidyan et al. (1971), in Yugoslavia its cultivation descends to 80 m. The domesticated einkorn is not demanding as to growth conditions. It can be cultivated on poor barren mountain soils. However, under good agricultural conditions, it gives excellent yields.

At present, *T.* *monococcum* occurs as an admixture with *T.* *dicoccon.* Mustafaev (1961) collected accessions in the fields of Nagorno-Karabakh and Nakhichevan. In western Georgia, domesticated einkorn formed part of the Zanduri complex, which also included *T.* *timopheevii* and *T.* *zhukovskyi*. […]

<296>[…] *Triticum* *monococcum* is a component of the Georgian Zanduri complex, which is remarkable for its resistance to fungal diseases. The resistance to such infection is higher in domesticated einkorn than in wild einkorn. […]

**[Chapter on *T. timopheevii*]**

<313> […] **History.** In the 8^th^ century CE, the Zanduri complex was a community of two species, *T.* *timopheevii* and *T.* *monococcum* (Menabde <314> 1948). Early on, *Triticum monococcum* took the predominant position. Later, *T.* *timopheevii* came to form the main part of the complex. The components of the Zanduri complex had their own vernacular names: “Chelta Zanduri” (Timofeev’s wheat) and “Gvatsa Zanduri” (domesticated einkorn).

**Geographical distribution.** Timofeev’s wheat is a narrowly local wheat species. Zhukovsky (1923) found it in 1922 in western Georgia. Stoletova (1925) specified the places of its cultivation. The Georgian triticologists Dekaprelevich and Menabde noted in a number of their works that the northern limit of Timofeev’s wheat cultivation centered around the towns of Tsageri, Orbeli, Lailashi, and Oni; its southern limit centered around Mekvena and Dgnorisa. Geographically, its cultivation was south of the Lechkhumi Range in the Caucasus Mountains in the present-day Ambrolauri, Oni, and Tsageri districts. Timofeev’s wheat is no longer cultivated there.

Ecologically, Timofeev’s wheat flourishes in the foothills of western Georgia (400–800 m) in a humid, cool climate.

[…]

Timofeev’s wheat in western Georgia was probably introduced from northeastern Turkey (Vavilov 1935). Menabde and Ericzjan (1942) associated the origin of *T.* *timopheevii* with the region of the ancient kingdom of Urartu, whence immigrant ancestors of modern-day Georgians introduced it into western Georgia. Certainly, the possibility of introduction of Timofeev’s wheat into Georgia from the south should not be categorically rejected.

[…]

Science does not yet possess sufficient information to decisively solve the question of the place of origin of *T.* *timopheevii.* While discussing the exceptional resistance of this species to fungal diseases, Vavilov (1964) emphasized that it could have developed under the warm, humid climatic conditions of western Georgia. The place of origin of Timofeev’s wheat is thought to be just there by Abesadze (1929) and Menabde (1948).

[…]

**[Chapter on *T. zhukovskyi*]**

<318> **Origin.** Zhukovsky’s wheat was found in Georgia in the Zanduri complex (Menabde and Ericzjan 1958, 1960).

[…]

Experiments by Tavrin (1963), and Upadhya and Swaminathan (1965) have shown that Zhukovsky’s wheat is allohexaploid, possessing the genomes of *T.* *monococcum* and *T.* *timopheevii.* As early as 1959, Bowden depicted the genome formula of Zhukovsky’s wheat as the sum of the genomes of these species. This path for an origin of Zhukovsky’s wheat has been confirmed by the similarity of morphological features of its spike to spikes of amphidiploids and F_1_ hybrids obtained from both spontaneous and controlled hybridization of *T.* *timopheevii* and *T.* *monococcum* by Kostov (1936), Kandelaki (1946), Upadhya and Swaminathan (1963), and Tavrin (1963).

[…]

<319> Immunochemical and electrophoretic analyses of caryopsis gliadins indicate the participation of *T.* *monococcum* and *T.* *timopheevii* in the formation of Zhukovsky’s wheat (Konarev et al. 1971, Aniol 1973, Konarev et al. 1976). Hence, Zhukovsky’s wheat originated from a spontaneous hybridization between the diploid and the tetraploid components of the Zanduri complex. […]
