## Supplementary material for "Genetic diversity, distribution and domestication history of the neglected GGA^t^A^t^ genepool of wheat": Supplementary_Material_S28.docx

**Was the cultivation range of *T. timopheevii* (*s.str.*) wider in the recent past?**

**Materials and Methods:**

Based on the Chapter on *T. timopheevii* of Dorofeev et al. (1979) – see Supplementary Material SM19, the recent past distribution range of *T. timopheevii* s.str. has been described as: **“***The Georgian triticologists Dekaprelevich and Menabde noted in a number of their works that the northern limit of Timofeev’s wheat cultivation* ***centered around the towns of Tsageri, Orbeli, Lailashi, and Oni; its southern limit centered around Mekvena and Dgnorisa. Geographically, its cultivation was south of the Lechkhumi Range in the Caucasus Mountains in the present-day Ambrolauri, Oni, and Tsageri districts.*** *Timofeev’s wheat is no longer cultivated there. Ecologically, Timofeev’s wheat flourishes in the foothills of western Georgia (400–800 m) in a humid, cool climate.”*

Based on Supplementary Table S2, the following detailed collection site information is available for *T. timopheevii* samples:

| **Species** | **Subspecies** | **ACC NUMB** | **Collection site** |
| --- | --- | --- | --- |
| *T. timopheevii* | *timopheevii* | IG 45097 | Tsageri. Region of town Kutaisi, Georgia |
| *T. timopheevii* | *timopheevii* | PI 341802 | Khachuri, Shida Kartli region, Georgia |
| *T. timopheevii* | *timopheevii* | PI 343447 | Khachuri, Shida Kartli region, Georgia |
| *T. timopheevii* | *viticulosum* | PI 349054 | Khachuri, Shida Kartli region, Georgia |
| *T. timopheevii* | *viticulosum* | PI 352506 | Khachuri, Shida Kartli region, Georgia |
| *T. timopheevii* | *typicum* | PI 352508 | Khachuri, Shida Kartli region, Georgia |
| *T. timopheevii* | *viticulosum* | PI 352510 | Khachuri, Shida Kartli region, Georgia |
| *T. timopheevii* | *timopheevii* | PI 418585 | Khachuri, Shida Kartli region, Georgia |

Additional collection site details for *T. timopheevii* samples were obtained from Dr. E.V. Zuev, the head of the Department of Wheat Genetic Resources, VIR, St-Petersburg, Russia:

| **Species** | **ACC NUMB** | **Collection site** |
| --- | --- | --- |
| *T. timopheevii* | K-30922 | Tsageri, 45 km to N from t.Kutaisi |
| *T. timopheevii* | K-31684 | Tsageri, v.Acaras, 60 km to N from t.Kutaisi |
| *T. timopheevii* | K-35914 | Tsageri, 45 km to N from t.Kutaisi |
| *T. timopheevii* | K-35915 | Tsageri, 52 km to N from t.Kutaisi |
| *T. timopheevii* | K-35916 | Tsageri, 52 km to N from t.Kutaisi |

The following collecting site information is known for two *T. timopheevii* samples reported presumably to be collected in Turkey:

| **Species** | **Subspecies** | **ACC NUMB** | **Collection site** |
| --- | --- | --- | --- |
| *T. timopheevii* | *viticulosum* Zhuk. | PI 119442 | Market, Araç, near Kastamonu, Turkey |
| *T. timopheevii* | *viticulosum* Zhuk. | TA1900 | 32 km S of Denizli, near Kahramanmaraş, Turkey |

The geo-referenced collection sites are as follows:

| **Georgia** | **City/village** | Geographic coordinates |
| --- | --- | --- |
| North | Tsageri/ Tsagera | 42.64745404707501, 42.771396142564484 |
|  | Orbeli | 42.632624005949765, 42.83203989860203 |
|  | Lailashi | 42.60731908705502, 42.85927891794966 |
| East | Oni | 42.584715963856326, 43.44372862743692 |
| South | Mekvena | 42.47752421737309, 42.75505276321019 |
|  | Dgnorisa | 42.46290842016359, 42.81663889859892 |
| **Turkey** |  |  |
|  | Araç | 41.24328549392879, 33.32765934598587 |
|  | 32 km S of Denizli, near Kahramanmaraş | 37.425770, 29.091797 |

*Simulation approach:*

56 sampling points from Georgia and 52 from Turkey were simulated around the eight collecting sites (6 from Georgia, 2 from Turkey) to capture the regional agro-ecological variation.

We collected 19 potential drivers of crop diversity and altitude from the WorldClim version 2 database (Fick and Hijmans, 2017), freely available at http://www.worldclim.org, and downloadable at 2.5 arc-min spatial resolution.

Principal components analysis and *k*-means clustering (number of clusters equal to 2) were conducted using the bioclimatic variables for the sampling points in both countries.

Data analysis was done using R v3.3.1 (R Core Team, 2016).

*Results:*

Several bioclimatic variables showed significant differences between sampling sites in Georgia and in Turkey, more specifically for mean diurnal range, temperature seasonality, mean temperature of driest quarter and mean annual precipitation:

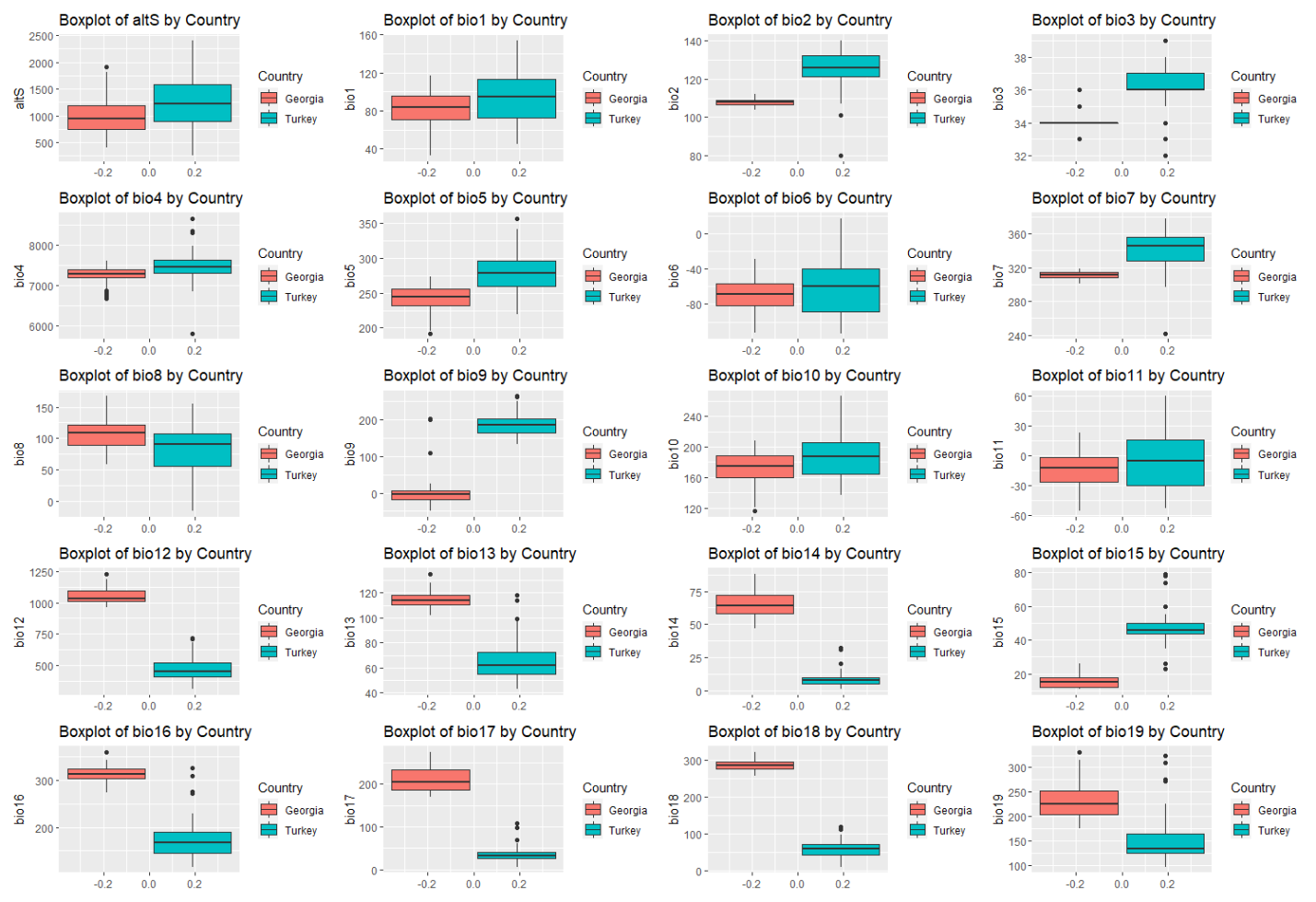

BIO1 = Annual Mean Temperature

BIO2 = Mean Diurnal Range (Mean of monthly (max temp - min temp))

BIO3 = Isothermality (BIO2/BIO7) (×100)

BIO4 = Temperature Seasonality (standard deviation ×100)

BIO5 = Max Temperature of Warmest Month

BIO6 = Min Temperature of Coldest Month

BIO7 = Temperature Annual Range (BIO5-BIO6)

BIO8 = Mean Temperature of Wettest Quarter

BIO9 = Mean Temperature of Driest Quarter

BIO10 = Mean Temperature of Warmest Quarter

BIO11 = Mean Temperature of Coldest Quarter

BIO12 = Annual Precipitation

BIO13 = Precipitation of Wettest Month

BIO14 = Precipitation of Driest Month

BIO15 = Precipitation Seasonality (Coefficient of Variation)

BIO16 = Precipitation of Wettest Quarter

BIO17 = Precipitation of Driest Quarter

BIO18 = Precipitation of Warmest Quarter

BIO19 = Precipitation of Coldest Quarter

The first two components from PCA analysis explained 80.3% of the total climatic variability. Plotting PC1 versus PC2 revealed a clear distinction between sampling points from the two countries:

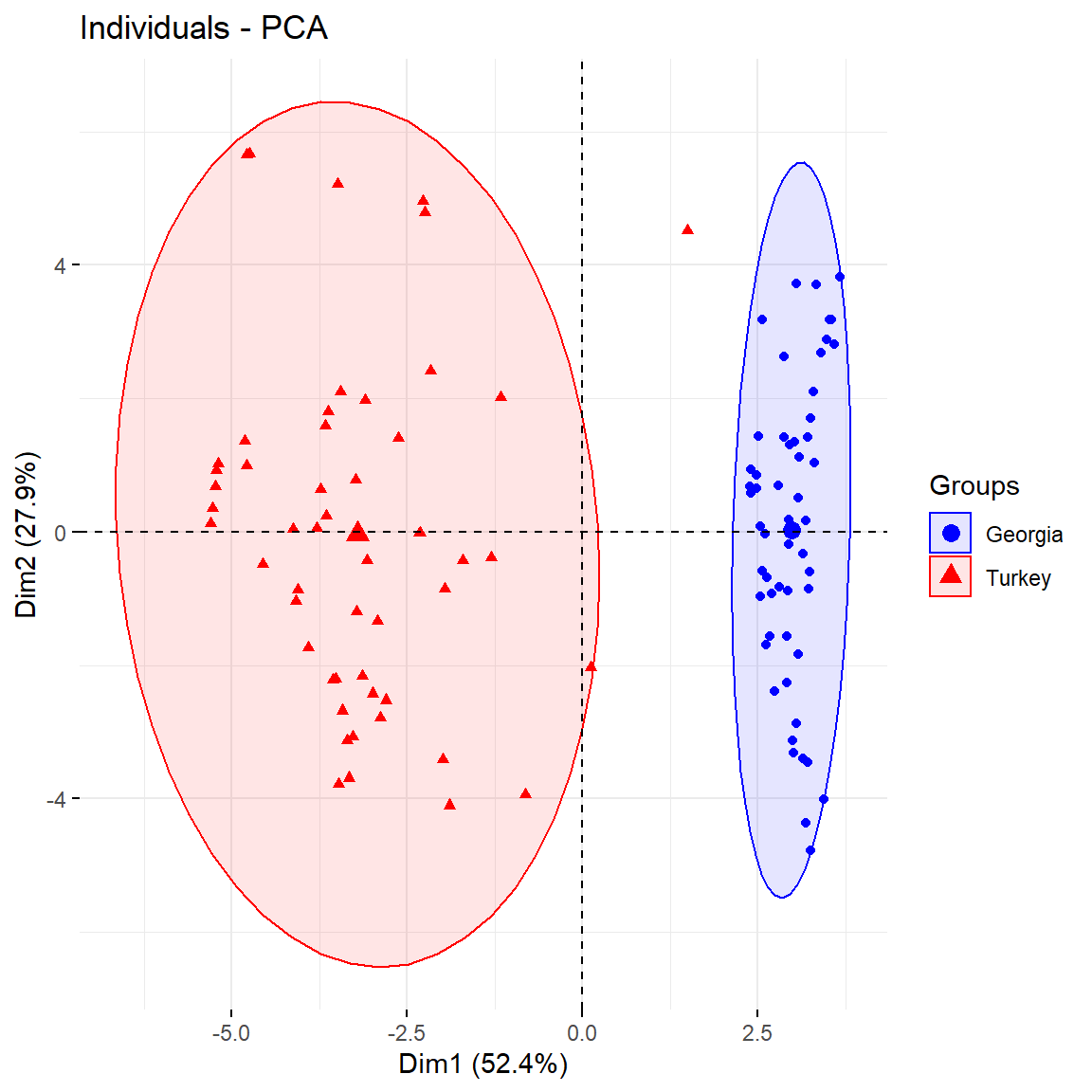

At *k* = 2, all sampling sites from Georgia were assigned to one cluster, and all Turkish sites to the second cluster (but except one site near Samsun in Turkey that was assigned to the Georgian cluster):

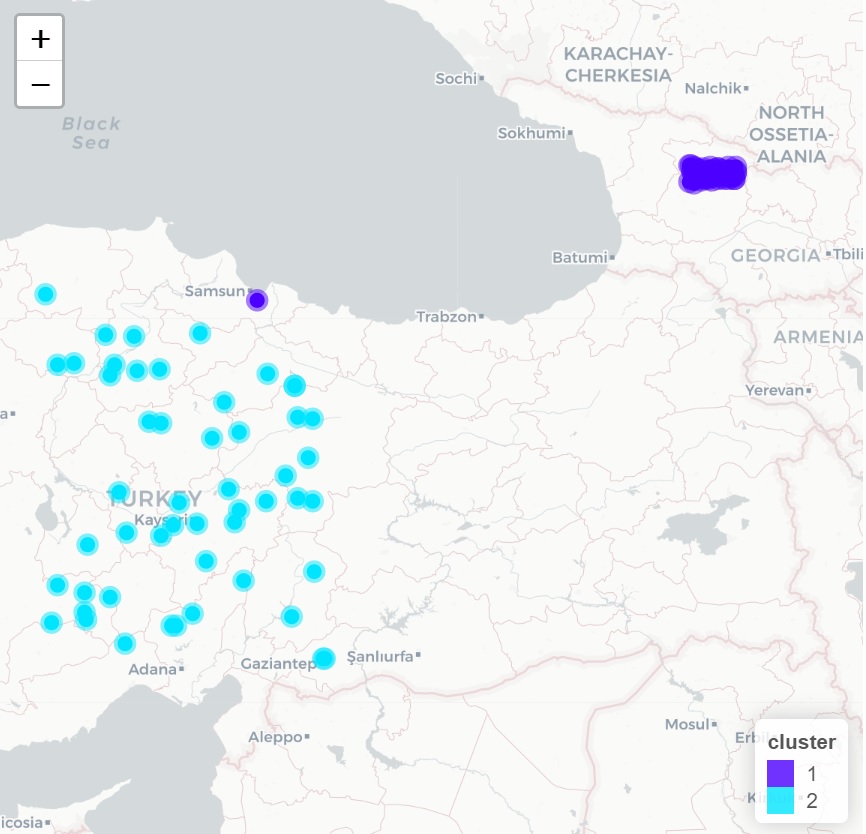

*Conclusion:*

Based on bioclimatic variables and our simulation of regional agro-ecological variation, we predict that *T. timopheevii* *s.str.* is maladapted to the climate outside Western Georgia.
