## Supplementary material for "Genetic diversity, distribution and domestication history of the neglected GGA^t^A^t^ genepool of wheat": Supplementary_TableS6.docx

**Supplementary Table S6**

**Table S6a. Summary statistics of genetic variation between four tetraploid wheat groups based on 401 *BARE-1*, 255 *Jeli* and the 656 markers combined.**

|  | ***BARE-1*** | | | | | |
| --- | --- | --- | --- | --- | --- | --- |
| **Species** | ***Na*** | ***Ne*** | ***I*** | ***He*** | ***uHe*** | ***%P*** |
| *T. timopheevii* (TIM) | 0.506 ± 0.039 | 1.032 ± 0.006 | 0.039 ± 0.008 | 0.023 ± 0.004 | 0.023 ± 0.004 | 17.21 |
| *T. araraticum* (ARA-1) | 0.454 ± 0.037 | 1.061 ± 0.010 | 0.058 ± 0.008 | 0.037 ± 0.006 | 0.038 ± 0.006 | 15.71 |
| *T. araraticum* (ARA-0) | 1.090 ± 0.048 | 1.077 ± 0.009 | 0.092 ± 0.008 | 0.053 ± 0.005 | 0.053 ± 0.005 | 50.87 |
| *T. dicoccoides* (DIC) | 1.860 ± 0.025 | 1.255 ± 0.014 | 0.277 ± 0.011 | 0.168 ± 0.008 | 0.168 ± 0.008 | 92.52 |
|  | ***Jeli*** | | | | | |
| *T. timopheevii* (TIM) | 0.773 ± 0.053 | 1.071 ± 0.011 | 0.078 ± 0.011 | 0.048 ± 0.007 | 0.048 ± 0.007 | 26.27 |
| *T. araraticum* (ARA-1) | 0.733 ± 0.053 | 1.113 ± 0.015 | 0.110 ± 0.013 | 0.071 ± 0.009 | 0.072 ± 0.009 | 25.88 |
| *T. araraticum* (ARA-0) | 0.973 ± 0.058 | 1.112 ± 0.015 | 0.117 ± 0.013 | 0.072 ± 0.008 | 0.072 ± 0.009 | 41.57 |
| *T. dicoccoides* (DIC) | 1.761 ± 0.039 | 1.312 ± 0.021 | 0.305 ± 0.015 | 0.193 ± 0.011 | 0.193 ± 0.011 | 86.27 |
|  | ***BARE-1* + *Jeli*** | | | | | |
| *T. timopheevii* (TIM) | 0.610 ± 0.032 | 1.047 ± 0.006 | 0.054 ± 0.006 | 0.032 ± 0.004 | 0.033 ± 0.004 | 20.73 |
| *T. araraticum* (ARA-1*)* | 0.563 ± 0.031 | 1.081 ± 0.009 | 0.078 ± 0.007 | 0.050 ± 0.005 | 0.051 ± 0.005 | 19.66 |
| *T. araraticum* (ARA-0) | 1.044 ± 0.037 | 1.090 ± 0.008 | 0.102 ± 0.007 | 0.060 ± 0.005 | 0.061 ± 0.005 | 47.26 |
| *T. dicoccoides* (DIC) | 1.822 ± 0.022 | 1.277 ± 0.012 | 0.288 ± 0.009 | 0.178 ± 0.006 | 0.178 ± 0.006 | 90.09 |

*Na*: number of different alleles, *Ne*: number of effective alleles, *I*: Shannon’s information index, *He*: expected heterozygosity, *μHe*: unbiased expected heterozygosity, %P: percentage of polymorphic loci

**Table S6b. Nei's genetic distance between four tetraploid wheat groups based on 401 *BARE-1*, 255 *Jeli* and the 656 markers combined.**

|  | *BARE-1* | | |  | *Jeli* | | |  | *BARE-1* + *Jeli* | | |
| --- | --- | --- | --- | --- | --- | --- | --- | --- | --- | --- | --- |
| Species | ARA-0 | ARA-1 | DIC |  | ARA-0 | ARA-1 | DIC |  | ARA-0 | ARA-1 | (DIC) |
| *T. araraticum* (ARA-0) | ---- |  |  |  | ----- |  |  |  | ----- |  |  |
| *T. araraticum* (ARA-1) | 0.071 | ----- |  |  | 0.132 | ----- |  |  | 0.094 | ----- |  |
| *T. dicoccoides* (DIC) | 0.234 | 0.224 | ----- |  | 0.298 | 0.280 | ----- |  | 0.258 | 0.245 | ----- |
| *T. timopheevii* (TIM) | 0.058 | 0.090 | 0.274 |  | 0.119 | 0.091 | 0.320 |  | 0.081 | 0.091 | 0.292 |
