## Supplementary material for "Genetic diversity, distribution and domestication history of the neglected GGA^t^A^t^ genepool of wheat": Supplementary_TableS8.docx

**Supplementary Table S8**

**Table S8a.** **Summary statistics of genetic variation between 103 *Triticum* genotypes based on 146 polymorphic AFLP markers.**

| **Species** | ***Na*** | ***Ne*** | ***I*** | ***He*** | ***uHe*** | ***%P*** |
| --- | --- | --- | --- | --- | --- | --- |
| *T. dicoccon* | 1.103±0.071 | 1.237±0.028 | 0.213±0.023 | 0.141±0.016 | 0.151±0.017 | 41.78 |
| *T. durum* | 0.767±0.062 | 1.121±0.023 | 0.105±0.018 | 0.071±0.013 | 0.082±0.015 | 19.18 |
| *T. timopheevii* (TIM) | 0.712±0.068 | 1.136±0.024 | 0.118±0.019 | 0.079±0.013 | 0.082±0.014 | 23.29 |
| *T. araraticum* (ARA-1) | 0.801±0.072 | 1.169±0.026 | 0.147±0.021 | 0.098±0.014 | 0.102±0.015 | 29.45 |
| *T. araraticum* (ARA-0) | 0.973±0.074 | 1.187±0.027 | 0.167±0.021 | 0.110±0.015 | 0.113±0.015 | 38.36 |
| *T. dicoccoides* (DIC) | 1.678±0.053 | 1.374±0.030 | 0.343±0.022 | 0.224±0.016 | 0.227±0.016 | 77.40 |

*Na*: number of different alleles, *Ne*: number of effective alleles, *I*: Shannon’s information index, *He*: expected heterozygosity, *μHe*: unbiased expected heterozygosity, %p: percentage of polymorphic loci

**Table S8b.** **Nei's genetic distance between 103 *Triticum* genotypes based on 146 polymorphic AFLP markers.**

| **Species** | *T. dicoccon* | *T. durum* | *T. timopheevii* (TIM) | *T. araraticum* (ARA-1) | *T. araraticum* (ARA-0) |
| --- | --- | --- | --- | --- | --- |
| *T. durum* | 0.072 |  |  |  |  |
| *T. timopheevii* (TIM) | 0.573 | 0.711 |  |  |  |
| *T. araraticum* (ARA-1) | 0.469 | 0.597 | 0.108 |  |  |
| *T. araraticum* (ARA-0) | 0.528 | 0.673 | 0.150 | 0.092 |  |
| *T. dicoccoides* (DIC) | 0.074 | 0.141 | 0.440 | 0.363 | 0.405 |
