## Supplementary material for "Genetic diversity, distribution and domestication history of the neglected GGA^t^A^t^ genepool of wheat": Supplementary_TableS10.docx

**Supplementary Table S10**

**Table S10a. Summary statistics of genetic variation between 265 genotypes based on 96 polymorphic C-bands considered for chromosomal passports construction.**

| **Species** | **Na** | **Ne** | **I** | **He** | **uHe** | **%P** |
| --- | --- | --- | --- | --- | --- | --- |
| *T. timopheevii* | 0.493±0.042 | 1.096±0.012 | 0.090±0.010 | 0.058±0.007 | 0.060±0.007 | 20.58 |
| *T. araraticum* (ARA-1) | 1.491±0.045 | 1.236±0.015 | 0.252±0.012 | 0.154±0.008 | 0.156±0.008 | 74.41 |
| *T. araraticum* (ARA-0) | 1.631±0.040 | 1.217±0.015 | 0.233±0.012 | 0.142±0.008 | 0.142±0.008 | 81.53 |

Na: number of different alleles, Ne: number of effective alleles, I: Shannon’s information index, He: expected heterozygosity, *μHe*: unbiased expected heterozygosity, %p: percentage of polymorphic loci

**Table S10b. Nei's genetic distance between 265 genotypes based on 96 polymorphic C-bands considered for chromosomal passports construction.**

| **Species** | *T. araraticum* (ARA-1) | *T. araraticum* (ARA-0) |
| --- | --- | --- |
| *T. timopheevii* | 0.098 | 0.113 |
| *T. araraticum* (ARA-1) | ----- | 0.036 |
