## Supplementary material for "Genetic diversity, distribution and domestication history of the neglected GGA^t^A^t^ genepool of wheat": Supplementary_TableS11.docx

**Supplementary Table S11**

**Table S11a.** **Summary statistics of genetic variation between 95 genotypes based on polymorphic Spelt-1 and Spelt-52 markers** (Supplementary Table S25).

| **Species** | **Na** | **Ne** | **I** | **He** | **uHe** | **%P** |
| --- | --- | --- | --- | --- | --- | --- |
| *T. timopheevii* | 0.754±0.128 | 1.170±0.039 | 0.165±0.032 | 0.106±0.022 | 0.113±0.023 | 36.84 |
| *T. araraticum* (ARA-1) | 1.544±0.112 | 1.120±0.033 | 0.241±0.027 | 0.142±0.019 | 0.145±0.019 | 77.19 |
| *T. araraticum* (ARA-0) | 1.789±0.082 | 1.160±0.027 | 0.211±0.023 | 0.117±0.016 | 0.118±0.016 | 89.47 |

Na: number of different alleles, Ne: number of effective alleles, I: Shannon’s information index, He: expected heterozygosity, *μHe*: unbiased expected heterozygosity, %p: percentage of polymorphic loci

**Table S11b.** **Nei's genetic distance between 95 genotypes based on polymorphic Spelt-1 and Spelt-52 markers** (Supplementary Table S25).

| **Species** | *T. araraticum* (ARA-1) | *T. araraticum* (ARA-0) |
| --- | --- | --- |
| *T. timopheevii* | 0.022 | 0.038 |
| *T. araraticum* (ARA-1) | ----- | 0.013 |
