## Supplementary material for "Genetic diversity, distribution and domestication history of the neglected GGA^t^A^t^ genepool of wheat": Supplementary_TableS13.docx

**Supplementary Table S13**

**Variants of chromosomal rearrangements in *T. araraticum*.** No – identification number of rearrangements (according to Fig. 4); variant of rearrangement (in brief); Structure of rearranged chromosomes; Chromosomal group; the # lines carrying rearrangement; Country of origin; Collection site; Accession #.

| **No** | **Variant of rearrangement** | **Structure of rearranged chromosomes** | **Group** | **# lines** | **Country of origin** | **Collection site** | **Accession #** |
| --- | --- | --- | --- | --- | --- | --- | --- |
| 1 | inv 2A^t^ | parInv2A^t^L | ARA-0 | 4 | Iraq | 1 km NE of Salahaddin | PI 503292; PI 538470; PI 427311; TRI 11939 |
| 2 | inv 3A^t^ | *per*Inv3A^t^ | ARA-1 | 2 | Turkey | 2 km West of Kavşak road Kuşçu | IG 116168 |
|  |  |  | ARA-1 |  |  | 36 km NE from Kilis to Gaziantep | 2634 |
| 3 | inv 4A^t^ | *per*Inv4A^t^ | ARA-1 | 1 | Turkey | between T. Karadut and Kizilkent Villages; 2 km before Kizilkent | IG 116169 |
| 4* | inv 7A^t^-1 | *per*iInv7A^t^-1 | ARA-0 | 16 | Iran | 40 km NW of Shahabad | PI 427368; PI 427369 |
|  |  |  | ARA-0 |  | Iran | 47 km NW from Eslamabad (Shahabad) to Sar-e Pol-e Zahab | PI 427365; PI 427366; TA1540; TA42; TA43; TA5; Cltr 17680 |
|  |  |  | ARA-0 |  | Iran | SE of Shahabad | TA 1543 |
|  |  |  | ARA-0 |  | Iraq | 43 km NW of Sulaymaniyah to Surdash | PI 427356; PI 427357b; TA1534; TA1535 |
|  |  |  | ARA-0 |  | Mistaken | Unknown | PI 427304b |
| 5 | inv 7A^t^-2 | *per*Inv7A^t^-2 | ARA-0 | 1 | Iraq | 1 km NE of Salahaddin | PI 427381 |
| 6 | inv 4G | *per*Inv4G | ARA-0 | 1 | Iraq | 11.4 km ENE from Koi Sanjaq to Ranya | KU-8705 |
| 7 | inv 5G | *per*Inv5G | ARA-1 | 1 | Turkey | 36 km NE from Kilis to Gaziantep | 2633 |
| 8 | inv 6G | *per*Inv6G | ARA-0 | 4 | Iraq | 2 km S from the position of 38.9 km from Rowanduz to Rayat | KU-8763 |
|  |  |  | ARA-0 |  | Iraq | 4.8 km NE from Shaqlawa to Rowanduz | KU-8774 |
|  |  |  | ARA-0 |  | Iraq | 52.4 km SW from Sulaymaniyah to Surdash | KU-8657 |
|  |  |  | ARA-0 |  | Iraq | Shaqlawa | KU-8802 |
| 9* | T1G:2G-1 | T1GS:2GS + T1GL:2GL | ARA-0 | 1 | Iraq | 6 km E of Suara Tuka | PI 427349 |
| 10* | T1G:2G-2 | T1GS:2GL + T2GS:1GL | ARA-0 | 8 | Armenia | 8 km W of Garni (Erevan-Garni) | TA2892 |
|  |  |  | ARA-0 |  | Armenia | Abovian reg., between vil. Garni and Azavan | K-61656 |
|  |  |  | ARA-0 |  | Armenia | Abovian reg., Shor-Bulag | i-059443 |
|  |  |  | ARA-0 |  | Armenia | Aragatsotn reg., Geghadir village | k-58667 |
|  |  |  | ARA-0 |  | Armenia | E of Erevan, forestry station | K-61654b; TA1566 |
|  |  |  | ARA-0 |  | Azerb. | Nakhchivan, E of Aznaburt, Mt Dash Aghl | K-28244b; K-28280b |
| 11* | T1G:3G | T1GS:3GL + T3GS:1GL | ARA-0 | 6 | Iraq | 22.4 km W from Amadiyah to Bamarni | KU-8907 |
|  |  |  | ARA-0 |  | Iraq | 3-4 km E of Suara-Tuka, Mazorka Gorge | PI 427353; TA27; TA902 |
|  |  |  | ARA-1 |  | Syria | 4 km N of St. Simeon; road to Afrin | IG 117895c |
|  |  |  | ARA-1 |  | Turkey | 1 km before Sarıbuğday road from Yavuzeli | IG 116176 |
| 12* | T1G:4G-1 | T1GS:4GL + T1GL:4GS | ARA-0 | 1 | Iraq | 58.3 km NW of As-Sulaymaniyah to Surdash | TA860 |
| 13* | T1G:4G-2 | T1GS:4GS + T4GL:1GL | ARA-0 | 7 | Iraq | 21 km S of Harir, between Rowanduz and Shaqlawa | TA1520, *different from* PI 538492 |
|  |  |  | ARA-0 |  | Iraq | 7 km NE of Shaqlawa | PI 427399; PI 427400; PI 538488; TA14; TA16; TA1517; |
| 14* | T1G:5G-1 | T1GS:5GL + T5GS:1GL | ARA-0 | 3 | Armenia | near Erevan-Shorbulag, towards Erevan – Shorbulag | TA2890 |
|  |  |  | ARA-0 |  | Azerb. | Akhsuinskyi pass | i-082532b |
|  |  |  | ARA-1 |  | Turkey | 1 km North of Akçaburç | IG 116165 |
| 15 | T1G:5G-3 | T1GL.1GS-5GS + T5GL.5GS-1GS | ARA-0 | 1 | Turkey | Oak forest | TRI 17028 |
| 16 | T1G:6G | T1GS:6GS + T1GL:6GL | ARA-0 | 1 | Iraq | 10.6 km ENE from Koi Sanjaq to Ranya | KU-8710 |
| 17* | T2A^t^:4G | T2A^t^S:4GS + T2A^t^L:4GL | ARA-0 | 6 | Armenia | 8 km W of Garni (Erevan-Garni) | TA2894 |
|  |  |  | ARA-0 |  | Armenia | near Erevan-Shorbulag | K-30258b |
|  |  |  | ARA-0 |  | Azerb. | Nakhchivan region | K-31627a |
|  |  |  | ARA-0 |  | Azerb. | Nakhchivan, E of Aznaburt, Mt Dash Aghl | K-61655; TRI 16598 |
| 18 | T2A^t^:5G | T2A^t^S:5GL + T5GS:2A^t^L | ARA-0 | 1 | Iraq | 53 km NW from Sulaymaniyah to Dukan Dam | KU-8682 |
| 19* | T2A^t^:6A^t^ | T2A^t^S:6A^t^S + T2A^t^L:6A^t^L | ARA-0 | 2 | Turkey | Tunceli, 17 km S of Tunceli on road from Elazığ to Erzincan | CItr 17677; PI 427440 |
| 20* | T2G:4G | T2GS:4GS + T2GL:4GL | ARA-0 | 6 | Iraq | 4 km NE of Shaqlawa, Forest or woodland | PI 427334a; PI 538431; TA23; TA888 |
|  |  |  | ARA-0 |  | Turkey | 6 km east of Eruh on the Eruh-Sirnak road. Habitat: Steep, south facing slope in canyon; hard limestone with oak trees | PI 596290 |
|  |  |  | ARA-0 |  | Turkey | 6 km W of Uzumluk village on Siirt-Eruh road | PI 560873 |
| 21 | T2G:5G-1 | T2GL.2GS-5GL + T5GS.5GL-2GS | ARA-1 | 3 | Syria | 2 km W of St. Simeon; left of the road to Basota | IG 117891 |
|  |  |  | ARA-1 |  | Syria | 4 km N of St. Simeon; road to Afrin | IG 117895b |
|  |  |  | ARA-1 |  | Turkey | 2 km SW from Sarıbuğday | IG 116177b |
| 22 | T2G:5G-2 | T2GL.2GS-5GS + T5GL.5GS-2GS | ARA-1 | 1 | Turkey | Cibeker Village or Bağbaşı (old village name) | IG 116164 |
| 23 | T2G:5G-3 | T2GS.2GL-5GL + T5GS.5GL-2GL | ARA-1 | 1 | Turkey | 63 km SE from Türkoğlu (SW of Karadağ) | 3121, 3127 |
| 24* | T2G:6G-1 | T2GS:6GL + T6GS:2GL | ARA-0 | 1 | Iraq | 15.3 km ENE from Dohuk to Amadiyah | KU-8824A |
| 25 | T2G:6G-2 | T2GS:6GS + T2GL:6GL | ARA-1 | 2 | Turkey | 45 km SE of Maras (Maras – Gaziantep) | KU-1950; KU-1966 |
| 26* | T2G:7G | T2GS:7GS + T2GL:7GL | ARA-0 | 2 | Armenia | Abovian reg. | K-61657a, K-61656c |
| 27 | T3A^t^:6A^t^ | T3A^t^S.3A^t^L-6A^t^L + T6A^t^S.6A^t^L-3A^t^L | ARA-0 | 2 | Iran | Sar Dasht | IG 113294; IG 113295 |
| 28 | T3A^t^:7A^t^-1 | T3A^t^S:7A^t^L+7A^t^S:3A^t^L | ARA-1 | 4 | Turkey | 45 km NE from Kilis to Gaziantep | 2631; 2639; 2641; 2644 |
| 29* | T3G:4G-1 | T3GS:4GS + T3GL:4GL | ARA-0 | 1 | Iraq | 17.9 km W from Shaqlawa to Arbil, NE slope of Pirman Dagh | TA872 |
| 30 | T3G:4G-2 | T3GS.3GL-4GL + T4GS.4GL-3GL | ARA-0 | 2 | Iraq | 13.4 km W from Amadiyah to Bamarni | KU-8877; KU-8878 |
| 31 | T3G:4G-3 | T3GS:4GL + T4GS:3GL | ARA-0 | 1 | Azerb. | Nakhchivan, mt. Ali-Chapan South slope | TRI 7389a |
| 32 | T3G:5G | T3GL.3GS-5GS+T5GS.5GL-3GS | ARA-1 | 1 | Turkey | 45 km SE of Maras (Maras – Gaziantep) | KU-1984B |
| 33 | T3G:6G-1 | T3GS:6GS + T3GL:6GL | ARA-0 | 1 | Iraq | 43 km NW of Sulaymaniyah to Surdash | PI 427359 |
| 34 | T3G:6G-2 | T3GL.3GS-6GL + T6GS.6GL-3GS | ARA-1 | 2 | Syria | 2 km S of Kafr Nabil | IG 119456; IG 117895a |
| 35* | T3G:7G | T3GS:7GS + T3GL:7GL | ARA-0 | 2 | Iraq | 16 km E of Sulaymaniyah to Chuarta | TA1537 |
|  |  |  | ARA-0 |  | Iraq | 52.4 km SW from Sulaymaniyah to Surdash | KU-8601 |
| 36 | T4A^t^:3G | T4A^t^S:3GL + T4A^t^L:3GS | ARA-0 | 2 | Iraq | 14 km S from Sulaymaniyah to Qara Dagh, roadside refreshment house, NE slope of Shakh I Basranan | KU-8545; KU-8561 |
| 37 | T4A^t^:4G-1 | T4A^t^S:4GL+ T4A^t^L:4GS | ARA-0 | 1 | Turkey | 45 km SE of Maras (Maras – Gaziantep) | KU-1986 |
| 38 | T4A^t^:4G-2 | Т4A^t^L.4A^t^S-4GS + T4GL.4GS-4AS | ARA-0 | 3 | Iraq | 13.2 km S from Sulaymaniyah to Qara Dagh | KU-8497 |
|  |  |  | ARA-0 |  | Iraq | 14 km S from Sulaymaniyah to Qara Dagh, roadside refreshment house, NE slope of Shakh I Basranan | PI 427364; PI 538514 |
| 39* | T4G:5G-1 | T4GS:5GL + T5GS:4GL | ARA-0 | 2 | Armenia | unknown | TA1563 |
|  |  |  | ARA-0 |  | Azerb. | Shemakhinskyi reg. | K-61658 (i-059450) |
| 40 | T4G:5G-2 | T4GS:5GS + T4GL:5GL | ARA-0 | 1 | Iraq | 13 km W of Shaqlawa | PI 427346 |
| 41* | T4G:6G-1 | T4GS:6GL + T6GS:4GL | ARA-0 | 3 | Iraq | 14 km S from Sulaymaniyah to Qara Dagh, roadside refreshment house, NE slope of Shakh I Basranan | KU-8567 |
|  |  |  | ARA-0 |  | Iraq | 4 km NE of Shaqlawa, Forest or woodland | PI 427329 |
|  |  |  | ARA-1 |  | Turkey | Maras; Rocky basaltic area; 40 km S of Maras to Gizi Antep | TA1008 |
| 42 | T4G:7G-1 | T4GS:7GL + T7GS:4GL | ARA-0 | 1 | Turkey | 12 km E of Silvan (Diyarbakir – Malabadi) | KU-1938 |
| 43 | T4G:7G-2 | T4GS:4GL-7GL + T7GS.7GL-4GL | ARA-1 | 1 | Turkey | 3 km from Gelinbuğday village | PI 656869a |
| 44* | T5A^t^:1G | T5A^t^L.5A^t^S-1GS + T1GL.1GS-5A^t^S | ARA-0 | 8 | Iraq | 19 km E of Sulaymaniyah to Chuarta, field margin | PI 427363; PI 538511; PI 538512; TA170; TA35; TA171; TA172; TA173 |
| 45* | T5A^t^:3G | T3GS:5A^t^L + T5A^t^S:3GL | ARA-0 | 6 | Iraq | 7 km NE of Shaqlawa | PI 427339 |
|  |  |  | ARA-0 |  | Iraq | 7.1 km NE from Shaqlawa to Rowanduz | TA941 |
|  |  |  | ARA-0 |  | Iraq | Unknown | K-40123; TRI 17419 |
|  |  |  | ARA-0 |  | Mist. | Unknown | PI 352265, PI 355452 |
| 46 | T5A^t^:6A^t^ | T5A^t^S-6A^t^S-:6A^t^L + T5A^t^L.5A^t^S-6A^t^S | ARA-0 | 1 | Turkey | 17.3 km E from Silvan to Bitlis | KU-8923 |
| 47* | T5A^t^:6G | T5A^t^S:6GS + T5A^t^L:6GL | ARA-1 | 1 | Turkey | 45 km SE of Maras (Maras – Gaziantep) | 1022/86 |
| 48 | T6A^t^:7G | T6A^t^S:7GL + T7GS:6A^t^L | ARA-0 | 1 | Armenia | unknown | TRI 11564c |
| 49* | T6G:7G | T6GS:7GL + T7GS:6GL | ARA-0 | 13 | Iraq | 1 km NE of Salahaddin | TA7; TA8; TA20; TA21 |
|  |  |  | ARA-0 |  | Iraq | 2 km NW of Salahaddin | PI 427322; PI 427385; PI 427386; TA145; TA146; TA28; TA9 |
|  |  |  | ARA-0 |  | Iraq | 13 km W of Shaqlawa | PI 538445 |
|  |  |  | ARA-0 |  | Iraq | 17.9 km W from Shaqlawa to Arbil, NE slope of Pirman Dagh | KU-8719 |
| 50 | T7A^t^:2G | T7A^t^S.7A^t^L:2GL + T2GS.2GL-7A^t^L | ARA-0 | 1 | Iran | Sar Dasht | IG 113298 |
| 51 | T7A^t^:4G | T7A^t^S:4GL + T7A^t^L:4GS | ARA-0 | 1 | Iraq | 4 km NE of Shaqlawa, Forest or woodland | PI 427398 |
| 52 | T7A^t^:6G | T6GS:7AtL + T6GL:7AS | ARA-1 | 1 | Turkey | 45 km SE of Maras (Maras – Gaziantep) | KU-1982 |
| 53 | inv 7A^t^ + T1G:5G-2 | T1GS.1GL-5GL + inv 7A^t^ | ARA-0 | 3 | Iran | unknown | Ami-5; TA1575; i-0108435 |
| 54 | inv7A^t^ + T6A^t^:6G | *per*inv7A^t^ +T6A^t^S:6GS +T6A^t^L:6GL | ARA-0 | 3 | Iran | 12.2 km NW from Karand to Qasri Shirin | KU-8944; KU-8945; KU-8946 |
| 55 | T1G:2G-2 + T3G:4G | T1GS:2GL + T2GS:1GL + T3GS:4GL + 4GS:T3GL | ARA-0 | 1 | Azerb. | Nakhchivan, mt. Ali-Chapan South slope | TRI 7389b |
| 56* | T1G:2G-2 +T4G:6G-2 | T1GS:2GL + T2GS:1GL + T4GS:6GS + 4GL:T6GL | ARA-0 | 1 | Armenia | 8 km W of Garni (Erevan-Garni) | TA2893 |
| 57* | T2A^t^:4G + T4A^t^:7A^t^ | T2AtS:4GS + T2AtL:4GL + T4AtS:7AtS + T4AtL:7AtL | ARA-0 | 4 | Armenia | Environs of Kurbalanya; near Erevan | TRI 11945a; TRI 14269 |
|  |  |  | ARA-0 |  | Azerb. | Nakhchivan, E to Aznaburt, between Dash Aghl and Andok Uchak | K-58669b |
|  |  |  | ARA-0 |  | Armenia | Abovian reg. | K-61657a |
| 58* | T2A^t^:6A^t^:4G | T2A^t^S:6A^t^L + T4GS:6A^t^S + T2A^t^L:4GL | ARA-0 | 1 | Armenia | Ararat region, between vil. Areni and Chiva | TRI 16599 |
|  |  |  | ARA-0 |  | Armenia | Ekhednadzor reg., near Arani village | Guk-1 |
|  |  |  | ARA-0 |  | Armenia | unknown | K-59942; TRI 11564b; 2638, ARM33 |
|  |  |  | ARA-0 | 5 | Armenia | Ararat region, Chimankend vil. | i-062604 |
| 59 | T2A^t^:4G + T6A^t^:7A^t^ | T2A^t^S:4GS + T2A^t^L:4GL + T6A^t^S.6A^t^L-7A^t^L + T7A^t^L.7A^t^S-6A^t^L | ARA-0 | 1 | Armenia | Kotayk, Abovian reg | K-61657b |
| 60* | T2G:4G:6G | T2GS:6GL + T4GS:6GS + T2GL:4GL | ARA-0 | 13 | Iraq | 21 km S of Harir, between Rowanduz and Shaqlawa | PI 427343; PI 427406; PI 427407; PI 538490; PI 538491; PI 538492; T18; TA40; TA41 |
|  |  |  | ARA-0 |  | Iraq | 4 km NE of Shaqlawa, Forest or woodland | PI 427334b; TA11 |
|  |  |  | ARA-0 |  | Iraq | 7 km NE of Shaqlawa | TA17; TA947 |
| 61 | T3A^t^:7A^t^-1 +T4A^t^:7G | T3A^t^S:7A^t^L+7A^t^S:3A^t^L + T4A^t^S:7GS + T4A^t^L:7GL | ARA-1 | 1 | Turkey | 45 km NE from Kilis to Gaziantep | 2643 |
| 62 | T3A^t^:7A^t^-1+ T2G:6G | T3A^t^S:7A^t^L + 7A^t^S:3A^t^L + T2GS:6GS + T2GL:6GL | ARA-1 | 1 | Turkey | 45 km NE from Kilis to Gaziantep | 2642 |
| 63 | T3A^t^:7A^t^-2 + T4G:6G | T3A^t^S:7A^t^L + 7A^t^S:3A^t^L + T4GS:6GS + T4GL:6GL | ARA-1 | 1 | Turkey | Adiyaman, 4 km north of Yarpuzlu, | IG 46434 |
| 64 | T3G:4G:7G | T4GS:3GL + T7GL:4GL +T3GS:7GS | ARA-0 | 1 | Turkey | 18 km W Kurtalan | IG 46246 |
| 65* | T3G:6G-1 + T4G:5G-2 | T3GS:6GS + T3GL:6GL + T4GS:5GS + T4GL:5GL | ARA-0 | 3 | Iraq | 19.1 km W from Shaqlawa to Arbil, SW slope of Pirman Dagh | KU-8713 |
| 66 | T3G:7G +T6A^t^:6G | T3GS:7GS + T3GL:7GL + T6A^t^L.6A^t^S-6GS + T6GL.6GS-6A^t^S | ARA-0 |  | Iraq | 16 km E of Sulaymaniyah to Chuarta | PI 427361; PI 538509 |
| 67* | T4A^t^:4G-2 + T2G:6G-1 | T2GS:6GL + T6GS:2GL | ARA-0 | 1 | Iraq | 13.2 km S from Sulaymaniyah to Qara Dagh | KU-8460 |
| 68 | T4A^t^:6G + T2G:4G | T4A^t^L.4A^t^S-6GS+T6GL.6GS-4A^t^S + T2GS:4GL + T4GS:2GL | ARA-0 | 1 | Iraq | 41 km NW of As-Sulaymaniyah to Surdash, Forest or woodland, h=780. 35.815586N, 45.09613E | PI 538504a |
| 69* | T5A^t^:3G + T4G:5G-1 | T3GS:5A^t^L + T5A^t^S:3GL+ T4GS:5GL + T5GS:4GL | ARA-0 | 1 | Iraq | 7 km NE of Shaqlawa | TA934 |
| 70* | T5A^t^:7G + T4G:5G-3 | T5A^t^S:7GS + T5A^t^L:7GL + T5GL-4GS.4GL | ARA-0 | 1 | Iraq | 4.4 km NW from Amadiyah, Mazorka Gorge | TA972 |
| 71 | T6G:6A^t^ + T2G:4G-2 | T2GS:4GL + T4GS:2GL + T6GL.6GS-6A^t^S(sat) + T6A^t^L.6A^t^S-6GS(sat) | ARA-0 | 1 | Iraq | 53 km NW from Sulaymaniyah to Dukan Dam | KU-8686 |
| 72* | T2A^t^:4G:7G + T4A^t^:7A^t^ | T2A^t^S:4GS + T2A^t^L:7GS + T4GL:7GL + T4A^t^S:7A^t^S + T4A^t^L:7A^t^L | ARA-0 | 18 | Azerb. | Nakhchivan | PI 352264; TRI 11367; TRI 17218 |
|  |  |  | ARA-0 |  | Azerb. | Nakhchivan, Aznaburt, mt. Kabakh-Tape | PI 418579; K-30212 |
|  |  |  | ARA-0 |  | Azerb. | Nakhchivan, Aznaburt, mt. Vash Konja | K-28283 |
|  |  |  | ARA-0 |  | Azerb. | Nakhchivan, E of Aznaburt, Mt Dash Aghl | PI 427303 |
|  |  |  | ARA-0 |  | Azerb. | Nakhchivan, E to Aznaburt, between Dash Aghl and Andok Uchak | K-58669c; TRI 11365 |
|  |  |  | ARA-0 |  | Azerb. | Nakhchivan, mt. Ali-Chapan South slope | TRI 11362; TRI 11947; TRI 17417; TRI 6912; TRI 7388 |
|  |  |  | ARA-0 |  | Azerb. | Nakhchivan, near Aznaburt Ishanakan | K-31122; K-30268 |
|  |  |  | ARA-0 |  | Azerb. | Nakhchivan, Sary-Aghl | K-30234 |
| 73 | T2A^t^:7G + T6A^t^:5G:6G | T2A^t^S-7GS.7GL + T2A^t^L.2AtS-7GS + T5GS.5GL-6A^t^L + T6GS.6GL-5GL + T6A^t^S.6A^t^L-6GL | ARA-0 | 1 | Turkey | 5 km E Kozluk junction | IG 46247 |
| 74 | T3G:7G:7A^t^ +T6A^t^:6G | T3GS:7A^t^L + T7GS:7A^t^S + T3GL:7GL + T6A^t^L.6A^t^S-6GS + T6GL.6GS-6A^t^S | ARA-0 | 1 | Iraq | 16 km E of Sulaymaniyah to Chuarta | PI 427360 |
| 75* | T2A^t^:4G:6G:7G + T4A^t^:7A^t^ | T2A^t^S:6GL + T4GS:6GS + T2A^t^L:7GS + T4GL:7GL + T4A^t^S:7A^t^S + T4A^t^L:7A^t^L | ARA-0 | 1 | Azerb. | Nakhchivan region | TRI 16597 |
| 76* | T2A^t^:6A^t^:4G:6G + T1G:2G-1 | T2A^t^S:6A^t^L + T4GS: 6A^t^S + T2A^t^L:6GS + T4GL:6GL + T1GS:2GL + T2GS:1GL | ARA-0 | 2 | Armenia | Ararat region, between vil. Areni and Chiva | K-59940b |
|  |  |  | ARA-0 |  | Armenia | Ekhednadzor reg., between Agavnadzori vil. and Ekhegnadzor city | K-61659 |
