## Supplementary material for "Genetic diversity, distribution and domestication history of the neglected GGA^t^A^t^ genepool of wheat": Supplementary_TableS24.docx

**Supplementary Table S24**

**Distribution of Spelt-1 and Spelt-52 loci over chromosomes of 87 *T. araraticum,* seven *T. timopheevii* and one *T. zhukovskyi* genotypes.** Accession number; Number of Spelt-1 signals (per haploid genome); position of Spelt-1 loci on the A^t^ or G-genome chromosomes; position of Spelt-52 loci on the A^t^ or G-genome chromosomes; Chromosomal group. The signal size at the respective chromosomal position is assessed according to signal intensity: 1 – very weak; 2 – weak, 3 – medium, 4 – large, 5 – very large. Empty boxes indicate the absence of signal at the respective position.

|  |  | Spelt-1 | | | | | | | | | | | Spelt-52 | | | |  |
| --- | --- | --- | --- | --- | --- | --- | --- | --- | --- | --- | --- | --- | --- | --- | --- | --- | --- |
| Accession # | Total Spelt1 (per 1x) | 2AL | 6AS | 1GL | 2GL | 3GL | 4GL | 5GL | 6GS | 6GL | 7GS | 7GL | 2AS | 1GS | 2GS | 6GL | Chr. group |
| Cltr 17680 | 5 | 2 | 4 |  |  | 1 | 3 | 3 |  |  |  |  | 2 | 1 |  | 5 | ARA-0 |
| IG 113294 | 3 |  | 4 |  |  | 5 |  | 1 |  |  |  |  | 1 |  |  | 4 | ARA-0 |
| IG 113298 | 3 |  | 4 |  |  |  | 3 | 4 |  |  |  |  | 1 |  |  | 5 | ARA-0 |
| KU-8944 | 6 | 2 | 3 |  | 1 | 5 | 3 | 3 |  |  |  |  | 2 | 2 |  | 5 | ARA-0 |
| TRI 11509 | 5 |  | 3 |  | 4 | 3 | 2 | 1 | 2 |  |  |  | 1 | 1 |  | 5 | ARA-0 |
| KU-8948 | 3 | 3 |  |  |  |  | 3 | 4 |  |  |  |  | 2 | 2 |  | 5 | ARA-0 |
| KU-8824A | 5 | 1 |  | 2 | 4 | 4 |  | 3 |  |  |  |  | 2 |  |  | 5 | ARA-0 |
| KU-8866 | 4 | 2 |  |  | 5 |  | 1 | 2 |  |  |  |  | 2 |  |  | 4 | ARA-0 |
| KU-8907 | 4 | 2 |  |  |  |  | 2 | 4 |  |  | 5 |  | 2 |  |  | 5 | ARA-0 |
| PI 427350 | 6 | 2 | 3 |  |  |  | 3 | 2 | 3 |  | 5 |  | 2 |  |  | 4 | ARA-0 |
| PI 427416 | 4 | 1 |  |  |  |  | 3 | 4 |  |  | 5 |  | 1 | 2 |  | 5 | ARA-0 |
| PI 427417 | 4 | 3 |  |  |  |  | 3 | 4 |  |  | 5 |  | 1 | 2 |  | 4 | ARA-0 |
| PI 538518 | 5 | 1 |  |  |  | 3 | 1 | 3 |  |  | 3 |  | 2 | 2 |  | 4 | ARA-0 |
| TRI 11508 | 5 | 1 |  |  |  | 3 | 1 | 2 |  |  | 3 |  | 1 | 2 |  | 4 | ARA-0 |
| KU-8713 | 3 |  | 2 |  |  |  | 2 | 2 |  |  |  |  | 2 |  |  | 5 | ARA-0 |
| KU-8739 | 3 |  | 5 |  |  |  | 4 | 4 |  |  |  |  | 1 |  |  | 4 | ARA-0 |
| KU-8774 | 3 |  | 4 |  |  |  | 3 | 3 |  |  |  |  | 2 |  |  | 5 | ARA-0 |
| PI 427380 | 6 |  | 4 |  | 5 | 3 | 2 | 1 | 1 |  |  |  | 2 |  |  | 5 | ARA-0 |
| PI 427381 | 6 | 1 | 3 | 2 | 5 |  | 2 | 2 |  |  |  |  | 2 |  |  | 5 | ARA-0 |
| PI 427386 | 5 | 2 | 4 |  | 5 |  | 2 | 3 |  |  |  |  | 2 |  |  | 5 | ARA-0 |
| PI 427392 | 4 |  | 3 |  | 5 |  | 2 | 2 |  |  |  |  | 1 |  |  | 5 | ARA-0 |
| PI 427398 | 3 |  | 4 |  |  |  | 2 | 2 |  |  |  |  | 2 | 1 |  | 5 | ARA-0 |
| PI 427403 | 5 |  | 2 |  | 5 |  | 2 | 2 | 3 |  |  |  | 1 |  |  | 5 | ARA-0 |
| PI 427407 | 4 |  | 3 |  | 5 |  | 2 | 3 |  |  |  |  | 1 |  |  | 5 | ARA-0 |
| PI 538461 | 4 |  | 3 |  | 5 |  | 1 | 2 |  |  |  |  | 2 |  |  | 5 | ARA-0 |
| PI 538516 | 5 |  | 2 |  | 5 | 2 | 2 | 2 |  |  |  |  | 2 | 2 |  | 5 | ARA-0 |
| TRI 11507 | 3 |  | 4 |  |  |  | 4 | 4 |  |  |  |  | 2 |  |  | 5 | ARA-0 |
| TRI 11939 | 6 |  | 4 |  | 5 | 2 | 2 | 1 | 1 |  |  |  | 1 |  |  | 5 | ARA-0 |
| TRI 17418 | 6 |  | 3 |  |  |  | 2 | 2 |  |  |  |  | 1 |  |  | 5 | ARA-0 |
| TRI 17419 | 5 |  | 3 |  | 4 |  | 2 | 3 | 3 |  |  |  | 1 |  |  | 5 | ARA-0 |
| KU-8451 | 4 | 2 |  |  |  | 3 | 2 | 3 |  |  |  |  | 2 | 2 |  | 5 | ARA-0 |
| KU-8545 | 2 |  |  |  |  |  | 4 | 4 |  |  |  |  | 2 | 2 |  | **4** | ARA-0 |
| KU-8567 | 6 | 1 | 3 |  |  | 4 | 3 | 3 |  |  | 5 |  | 2 |  |  | 5 | ARA-0 |
| KU-8595 | 4 | 1 | 2 |  |  | 1 | 3 | 2 |  |  |  |  | 2 | 2 |  | 5 | ARA-0 |
| KU-8596 | 5 |  | 2 |  | 5 |  | 2 | 2 |  | 4 |  |  | 2 | 2 |  | 4 | ARA-0 |
| KU-8602 | 4 | 2 |  |  |  | 2 | 4 | 4 |  |  |  |  | 2 | 2 |  | 5 | ARA-0 |
| KU-8619 | 6 | 1 | 3 |  |  | 1 | 3 | 2 |  | 4 |  |  | 2 | 2 |  | 4 | ARA-0 |
| KU-8620 | 6 | 1 | 1 |  | 4 |  | 2 | 2 |  | 4 |  |  | 1 | 2 |  | 3 | ARA-0 |
| KU-8640 | 3 | 2 |  |  |  |  | 3 | 3 |  |  |  |  | 2 | 2 |  | 5 | ARA-0 |
| KU-8682 | **5** | 1 |  |  |  |  | 1 | 2 |  | 4 | **5** |  | 2 | 1 |  | 2 | ARA-0 |
| PI 427364 | 3 |  |  |  |  | 5 | 3 | 3 |  |  |  |  | 1 |  |  | 3 | ARA-0 |
| PI 538456 | 4 |  | 2 |  | 5 |  | 3 | 3 |  |  |  |  | 2 | 2 |  | 5 | ARA-0 |
| PI 538505 | 5 |  |  | 2 | 5 |  | 3 | 3 |  | 5 |  |  | 1 | 1 |  | 4 | ARA-0 |
| PI 538508 | 2 |  |  |  |  |  | 3 | 3 |  |  |  |  | 2 | 1 |  | 4 | ARA-0 |
| PI 538512 | 3 |  |  |  |  |  | 2 | 3 |  |  | 5 |  | 2 | 1 |  | 5 | ARA-0 |
| PI 538514 | 3 |  |  |  | 5 |  | 3 | 3 |  |  |  |  | 1 |  |  | 4 | ARA-0 |
| IG 117895 | 5 | 1 | 5 |  |  |  | 2 | 3 |  | 5 |  |  | 1 | 2 |  | 3 | ARA-0 |
| IG 117895c | 6 |  | 3 | 2 | 5 |  | 1 | 2 |  |  |  | 4 | 1 | 2 |  | 4 | ARA-1 |
| IG 119456 | 8 |  | 3 | 3 | 5 |  | 3 | 3 | 1 |  | 5 | 4 | 2 | 2 |  | 5 | ARA-1 |
| IG 117891 | 6 | 1 | 4 | 2 |  |  | 2 | 2 | 1 |  |  |  |  | 2 |  | 4 | ARA-1 |
| K-58667 | 5 |  | 5 | 3 |  |  | 4 | 3 | 3 |  |  |  | 1 |  |  | 5 | ARA-0 |
| K-61659 | 6 | 1 | 3 | 3 |  |  | 3 | 3 | 2 |  |  |  | 1 |  |  | 5 | ARA-0 |
| K-61654 | 5 |  | 5 | 3 |  |  | 3 | 3 | 2 |  |  |  | 1 |  |  | 4 | ARA-0 |
| K-61657a | 5 |  | 5 | 3 |  |  | 3 | 3 | 2 |  |  |  | 1 |  |  | 5 | ARA-0 |
| K-61659 | 5 |  | 5 | 3 |  |  | 3 | 3 | 1 |  |  |  | 1 |  |  | 5 | ARA-0 |
| KU-1901 | 5 |  | 4 | 3 |  |  | 3 | 3 | 1 |  |  |  | 1 |  |  | 5 | ARA-0 |
| TRI 16599 | 5 |  | 3 | 3 |  |  | 3 | 3 | 1 |  |  |  | 1 |  |  | 5 | ARA-0 |
| K-61658A | 4 |  | 5 | 2 |  |  | 2 | 3 |  |  |  |  |  |  |  | 4 | ARA-0 |
| K-61658B | 4 |  | 5 | 2 |  |  | 2 | 3 |  |  |  |  |  |  |  | 4 | ARA-0 |
| TRI 11356 | 5 |  | 3 | 3 |  |  | 3 | 3 | 2 |  |  |  | 1 |  |  | 5 | ARA-0 |
| TRI 11365 | 5 |  | 3 | 3 |  |  | 3 | 3 | 2 |  |  |  | 1 |  |  | 5 | ARA-0 |
| KU-1930 | 7 | 2 |  | 3 |  | 1 | 3 | 3 | 2 |  |  |  | 2 |  |  | 5 | ARA-0 |
| KU-1986 | 5 |  |  | 2 | 5 |  | 2 | 2 | 2 |  |  |  | 2 |  |  | 3 | ARA-0 |
| KU-8909 | 5 |  | 2 | 3 | 5 |  | 1 | 2 |  |  |  |  | 2 |  |  | 5 | ARA-0 |
| KU-8910 | 6 |  | 2 | 3 | 4 |  | 2 | 2 | 2 |  |  |  | 2 |  |  | 4 | ARA-0 |
| KU-8917 | 5 | 1 | 5 | 3 |  |  | 2 | 3 |  |  |  |  | 2 |  |  | 5 | ARA-0 |
| KU-8926 | 6 | 2 | 4 | 2 | 2 |  | 2 | 3 |  |  |  |  | 2 |  |  | 4 | ARA-0 |
| KU-8939 | 5 |  | 3 |  | 5 |  | 1 | 2 | 4 |  |  |  | 2 |  |  | 5 | ARA-0 |
| PI 560872 | 3 |  |  | 3 |  |  | 2 | 3 |  |  |  |  | 2 |  |  | 5 | ARA-0 |
| 3121 | 5 |  | 2 | 2 | 3 |  | 2 |  |  |  |  | 4 | 2 |  |  | 5 | ARA-1 |
| **3127** | 5 |  | 3 | 3 |  |  | 3 | 3 |  |  |  | 4 | 1 |  | 5 |  | ARA-1 |
| 2630 | 5 |  | 4 | 2 |  |  | 2 | 2 | 2 |  |  |  | 2 | 2 |  |  | ARA-1 |
| 2631 | 5 |  | 4 | 2 |  |  | 2 | 2 | 2 |  |  |  | 1 | 2 |  |  | ARA-1 |
| 2633 | 5 |  | 3 | 2 |  |  | 2 | 2 | 3 |  |  |  | 2 | 2 |  | 4 | ARA-1 |
| 2634 | 5 |  |  | 2 | 4 |  | 2 | 2 | 4 |  |  |  |  |  | 5 | 2 | ARA-1 |
| IG 116164 | 5 |  | 5 | 2 |  |  | 2 | 3 | 2 |  |  |  | 1 |  | 5 |  | ARA-1 |
| IG 116165 | 7 | 2 | 3 | 2 | 5 |  | 2 | 2 | 2 |  |  |  | 2 | 2 |  |  | ARA-1 |
| IG 116168 | 6 |  | 3 | 2 | 2 |  | 2 | 2 | 2 |  |  |  | 2 | 2 |  | 4 | ARA-1 |
| IG 116169 | 5 |  | 4 | 2 |  |  | 2 | 2 | 2 |  |  |  | 1 | 1 |  |  | ARA-1 |
| IG 116170 | 5 |  | 4 | 3 |  |  | 2 | 2 | 1 |  |  |  | 1 | 2 |  |  | ARA-1 |
| IG46434 | 5 |  | 2 | 2 |  |  | 2 | 2 | 3 |  |  |  | 3 | 2 |  |  | ARA-1 |
| KU-1950 | 6 | 1 | 5 | 2 |  |  | 2 | 2 | 1 |  |  |  |  | 1 | 2 | 5 | ARA-1 |
| KU-1960 | 5 | 1 | 2 |  |  |  |  | 1 | 2 |  | 4 |  | 2 |  |  | 3 | ARA-1 |
| KU-1982 | 4 |  | 2 |  |  |  | 3 | 3 |  |  | 5 |  | 2 | 1 |  | 5 | ARA-1 |
| KU-1984B | 6 |  | 5 | 2 |  |  | 2 | 3 | 1 |  |  | 4 | 2 | 2 | 5 |  | ARA-1 |
| PI 654340 | 5 |  | 5 | 2 |  |  | 2 | 3 | 1 |  |  |  | 1 |  | 5 |  | ARA-1 |
| TA1008 | 4 |  | 5 | 2 |  |  | 3 | 2 |  |  |  |  | 1 | 1 |  | 3 | ARA-1 |
| KU-107-4 | 7 | 2 | 4 | 2 | 4 |  | 2 | 2 | 1 |  |  |  | 2 | 2 |  |  | TIM |
| TRI 3433 | 6 | 1 | 2 | 2 | 2 |  | 2 | 3 |  |  |  |  | 2 | 2 |  |  | TIM |
| KU-1818 | 7 | 1 | 4 | 2 | 4 |  | 2 | 2 |  |  |  |  | 1 | 2 |  |  | TIM |
| KU-1819 | 6 | 1 | 3 | 3 | 3 |  | 3 | 1 |  |  |  |  | 2 | 2 |  |  | TIM |
| KU-1820 | 7 | 1 | 4 | 2 | 4 |  | 2 | 3 | 1 |  |  |  | 2 | 2 |  |  | TIM |
| KU-1821 | 6 | 1 | 4 | 2 | 4 |  | 3 | 3 |  |  |  |  | 1 | 2 |  |  | TIM |
| TRI 5352 | 6 | 1 | 5 | 2 | 4 |  | 2 | 2 |  |  |  |  | 1 | 2 |  |  | TIM |
| K-43063 (Zhuk) | 6 | 2 | 4 | 2 | 3 |  | 2 | 2 |  |  |  |  |  | 2 |  |  | TIM |
