## Supplementary material for "Genetic diversity, distribution and domestication history of the neglected GGA^t^A^t^ genepool of wheat": Supplementary_TableS26.docx

**Supplementary Table S26**

**Distribution of Spelt-1 and Spelt-52 loci in geographic/chromosomal groups in comparison with *T. timopheevii.*** Region/ Country of origin; # – number of genotypes per region/ country; frequencies of occurrence of Spelt-1 and Spelt-52 signals in particular positions are given in absolute values. Group – chromosomal group of genotypes in particular region defined by C-banding analysis. The frequencies of Spelt-1 and Spelt-52 loci for total *T. araraticum* (ARA) in comparison with *T. timopheevii* (TIM) groups are given in percentage.

|  |  |  | Spelt-1 | | | | | | | | | | | Spelt-52 | | | |  |
| --- | --- | --- | --- | --- | --- | --- | --- | --- | --- | --- | --- | --- | --- | --- | --- | --- | --- | --- |
| **Region/country of origin** | **#** | **Average Spelt-1 (per 1x)** | **2AL** | **6AS** | **1GL** | **2GL** | **3GL** | **4GL** | **5GL** | **6GS** | **6GL** | **7GS** | **7GL** | **2AS** | **1GS** | **2GS** | **6GL** | **Group** |
| Transcaucasia | 11 | 4.82 | 1 | 11 | 11 | 0 | 0 | 11 | 11 | 9 | 0 | 0 | 0 | 9 | 0 | 0 | 11 | ARA-0 |
| Iran | 6 | 4.17 | 3 | 5 | 0 | 2 | 4 | 5 | 6 | 1 | 0 | 0 | 0 | 6 | 4 | 0 | 6 | ARA-0 |
| Iraq | 40 | 4.33 | 18 | 23 | 3 | 17 | 12 | 39 | 40 | 5 | 5 | 3 | 0 | 40 | 19 | 0 | 40 | ARA-0 |
| *Dahuk* | 8 | 4.63 | 8 | 1 | 1 | 2 | 3 | 7 | 8 | 1 | 0 | 6 | 0 | 8 | 4 | 0 | 8 | ARA-0 |
| *Erbil* | 16 | 4.44 | 2 | 16 | 1 | 10 | 3 | 16 | 16 | 4 | 0 | 0 | 0 | 16 | 2 | 0 | 16 | ARA-0 |
| *Sulaymaniyah* | 16 | 4.06 | 8 | 6 | 1 | 5 | 5 | 16 | 16 | 0 | 5 | 3 | 0 | 16 | 14 | 0 | 16 | ARA-0 |
| Turkey | 24 | 5.19 | 6 | 22 | 23 | 9 | 1 | 25 | 25 | 18 | 0 | 2 | 3 | 24 | 12 | 6 | 16 | ARA-0 |
| *Eastern* | 8 | 5.25 | 3 | 5 | 7 | 5 | 1 | 8 | 8 | 4 | 0 | 0 | 0 | 8 | 0 | 0 | 8 | ARA-0 |
| *Western* | 18 | 5.17 | 3 | 17 | 16 | 4 | 0 | 17 | 17 | 14 | 0 | 2 | 3 | 16 | 12 | 6 | 8 | ARA-1 |
| Syria | 4 | 6.25 | 2 | 4 | 3 | 2 | 0 | 4 | 4 | 2 | 1 | 1 | 2 | 3 | 4 | 0 | 4 | ARA-1 |
| Total for *T. araraticum* (%): | 87 | 4.72 | 33.7 | 74.7 | 46.0 | 34.5 | 19.5 | 95.4 | 97.7 | 40.2 | 6.9 | 13.8 | 6.9 | 94.3 | 43.7 | 6.9 | 86.0 |  |
| Total for ARA-0 | 66 | 4.52 | 37.8 | 68.2 | 31.8 | 36.4 | 25.6 | 97 | 100 | 28.8 | 9.1 | 13.6 | 0 | 97 | 36.4 | 0 | 100 | ARA-0 |
| Total for ARA-1 | 21 | 5.38 | 19.0 | 95.2 | 90.5 | 28.6 | 0 | 95.4 | 95.4 | 76.2 | 0 | 14.3 | 23.8 | 85.7 | 71.4 | 28.7 | 52.4 | ARA-1 |
| Total for *T. timopheevii* incl. *T. zhukovskyi* (%): | 8 | 6.38 | 100 | 100 | 100 | 100 | 0 | 100 | 100 | 25 | 0 | 0 | 0 | 100 | 100 | 0 | 0 | TIM |
