## Supplementary figures and images for "Genetic diversity, distribution and domestication history of the neglected GGA^t^A^t^ genepool of wheat"

### Fig_S4.tiff

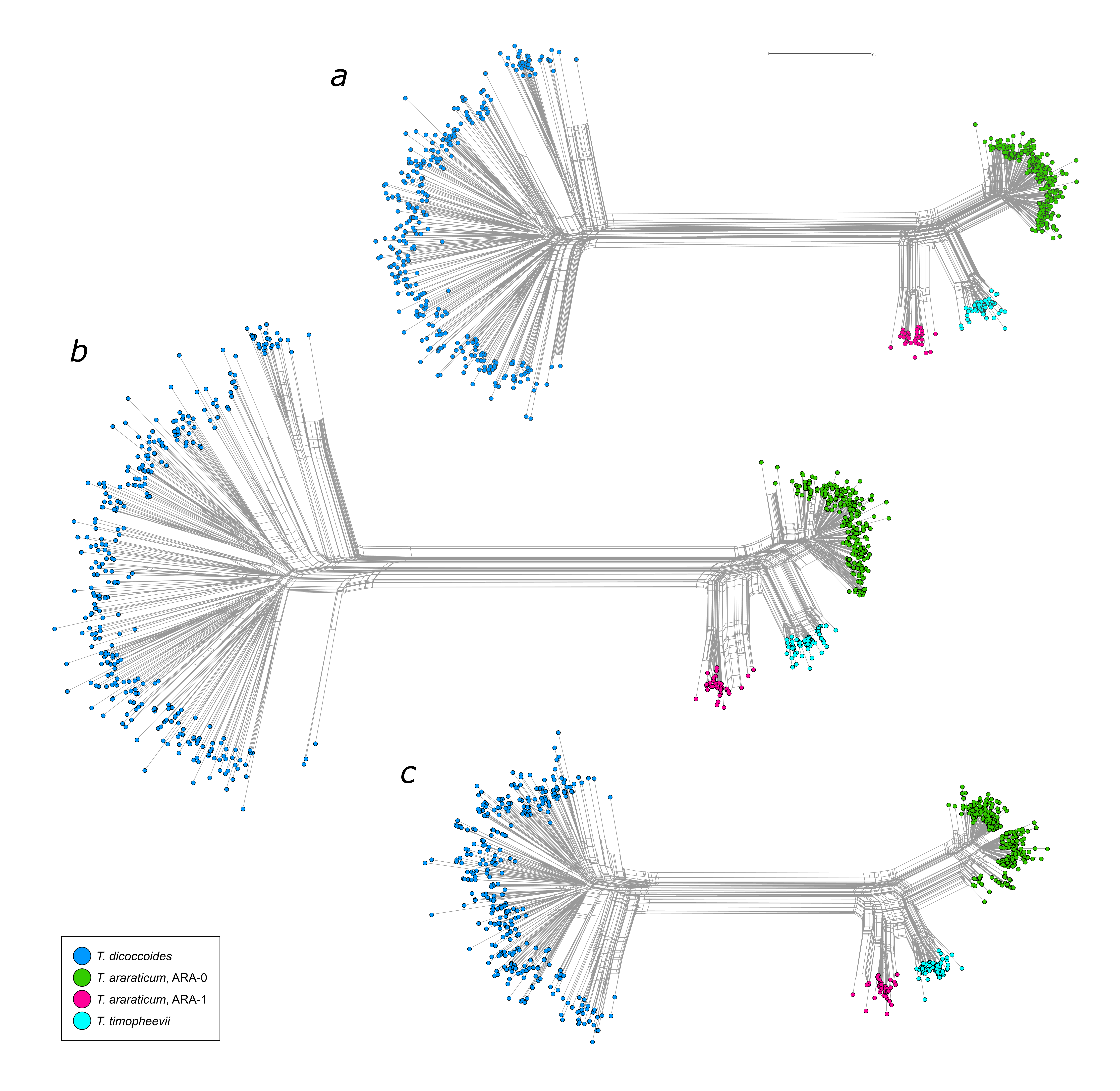

### Fig_S7.png

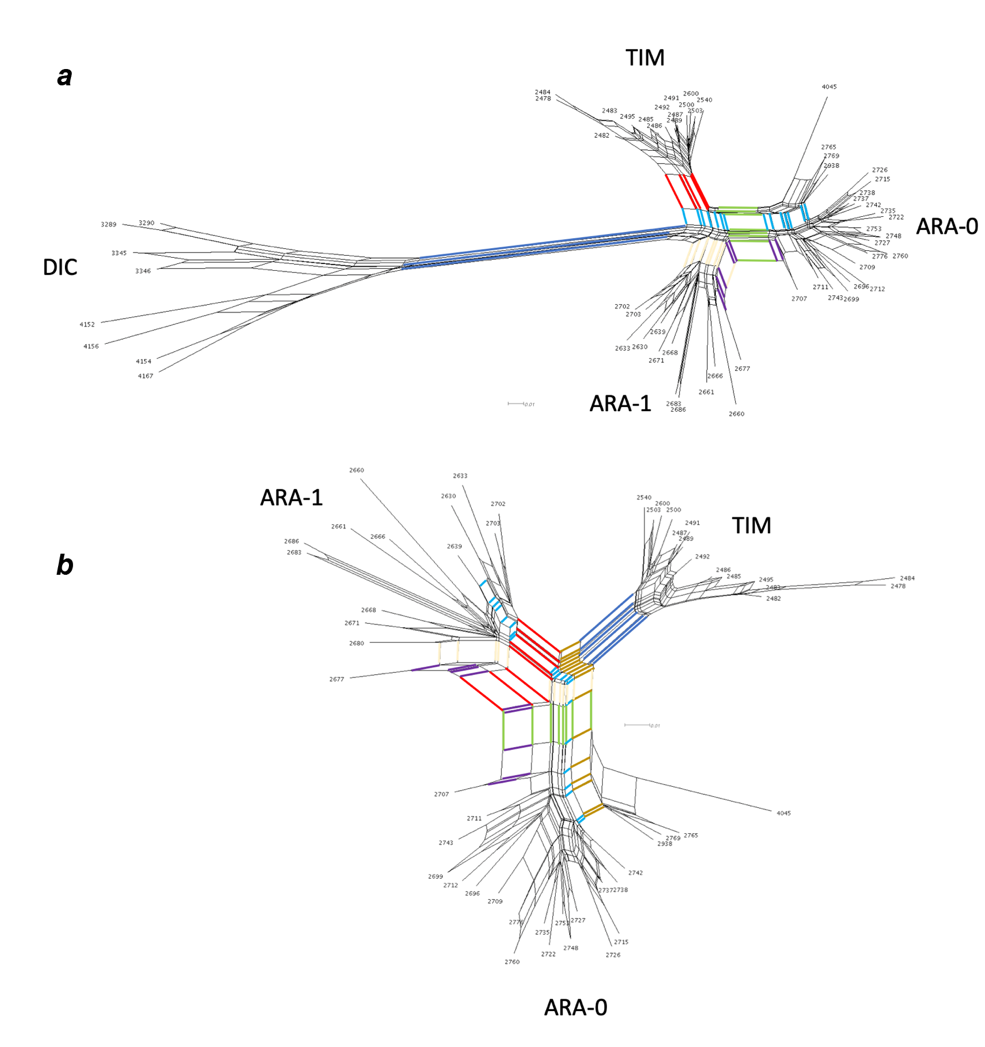

### Fig_S12.tif

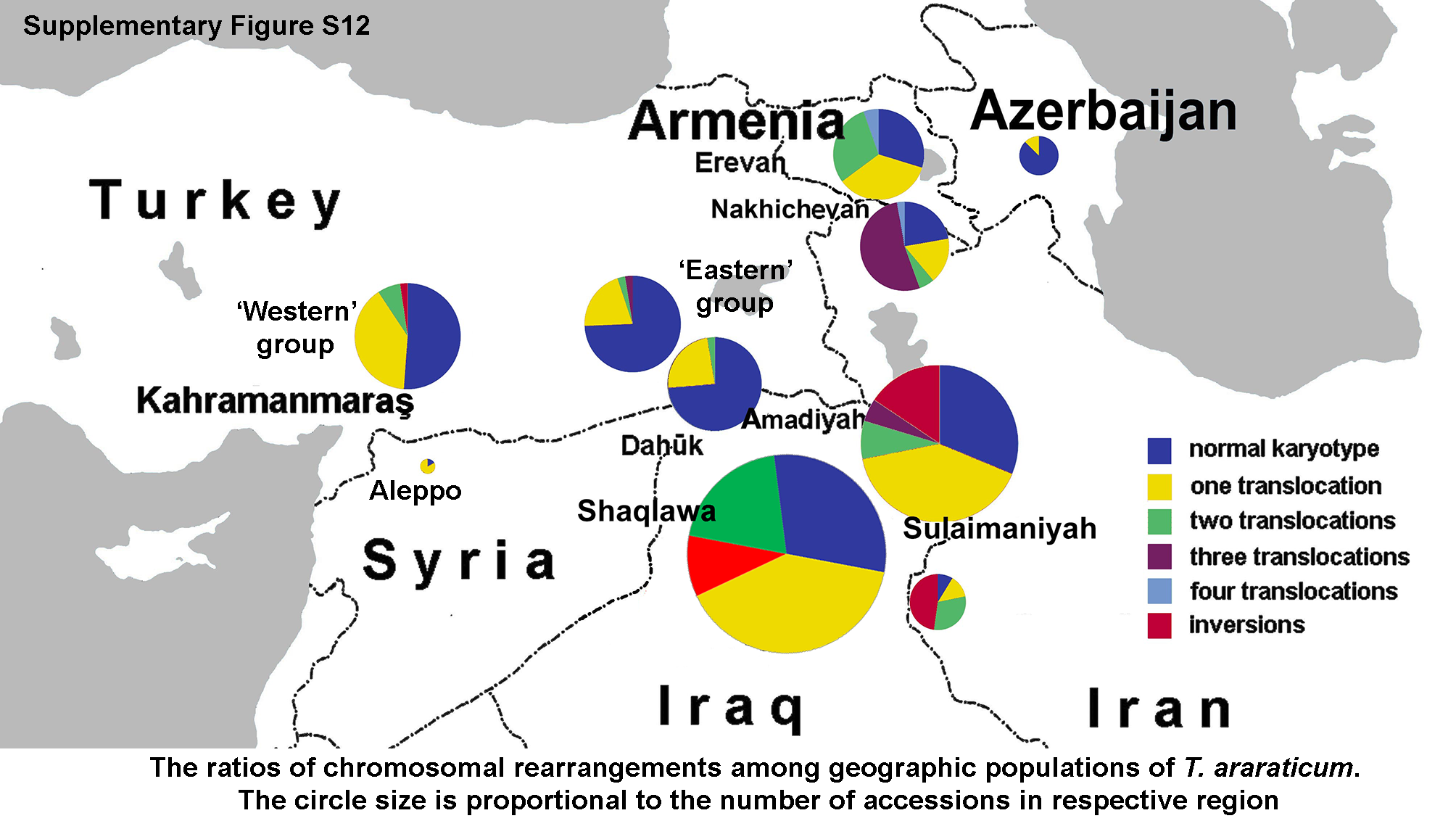

### Fig_S14.tif

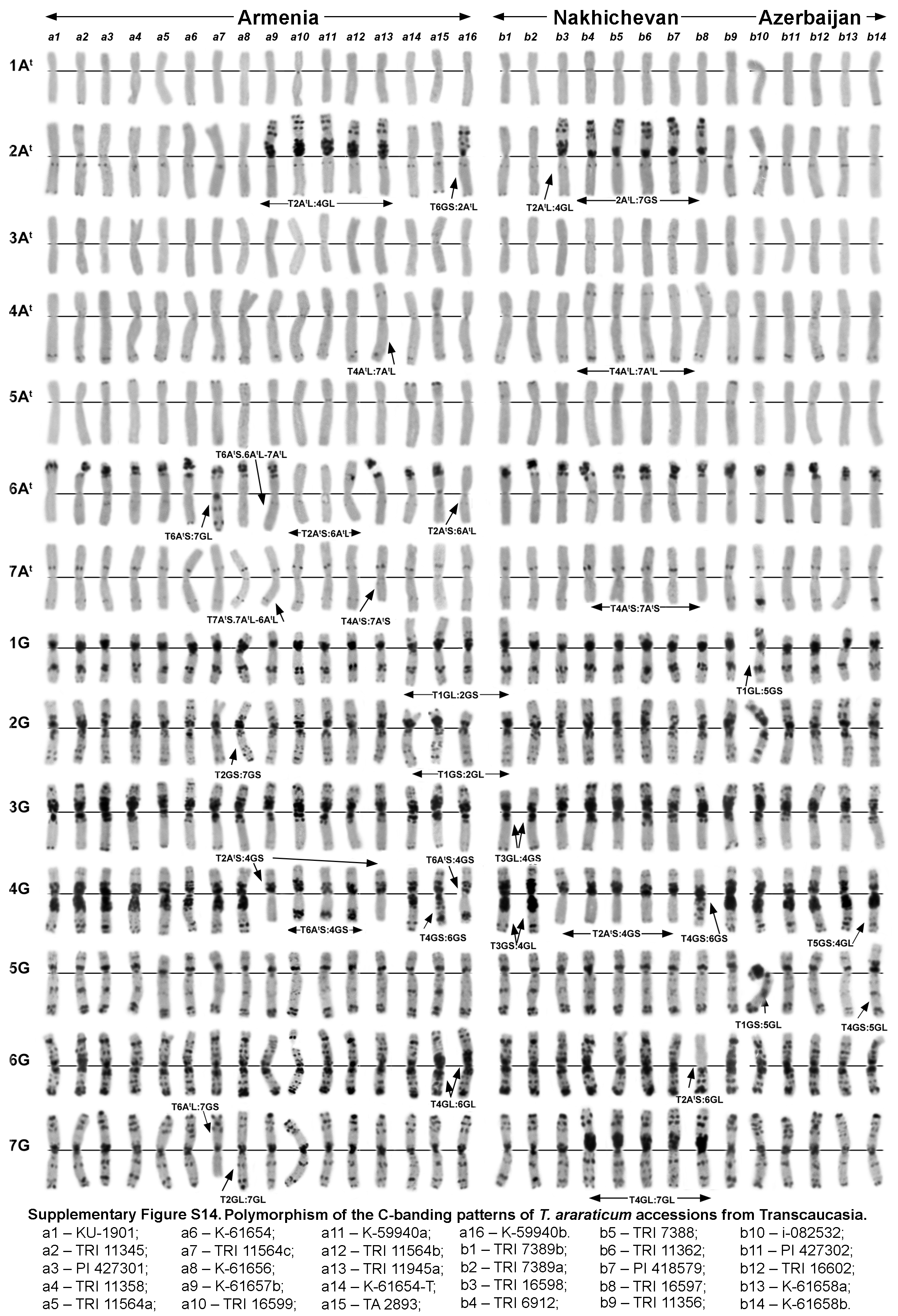

### Fig_S15.tif

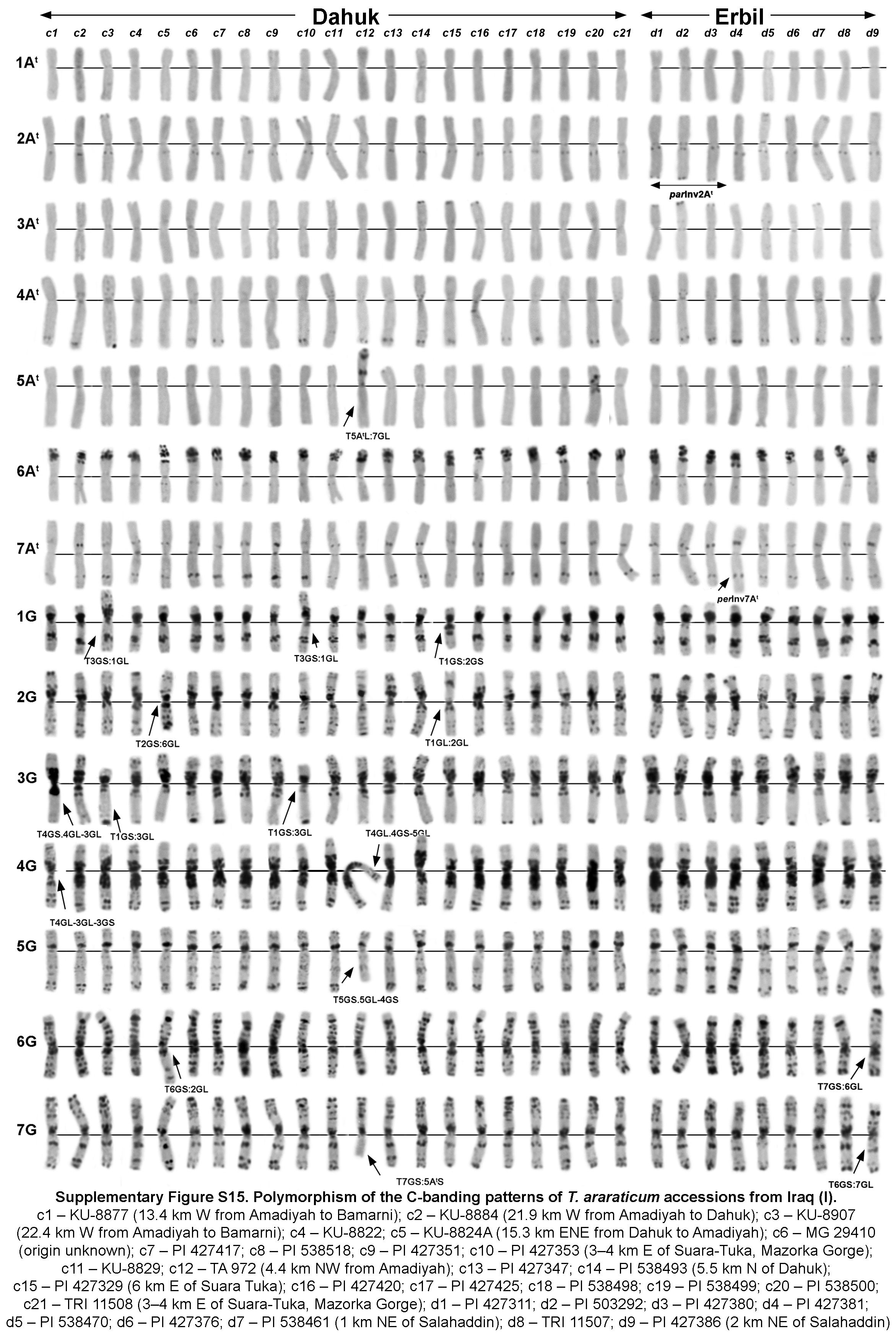

### Fig_S16.tif

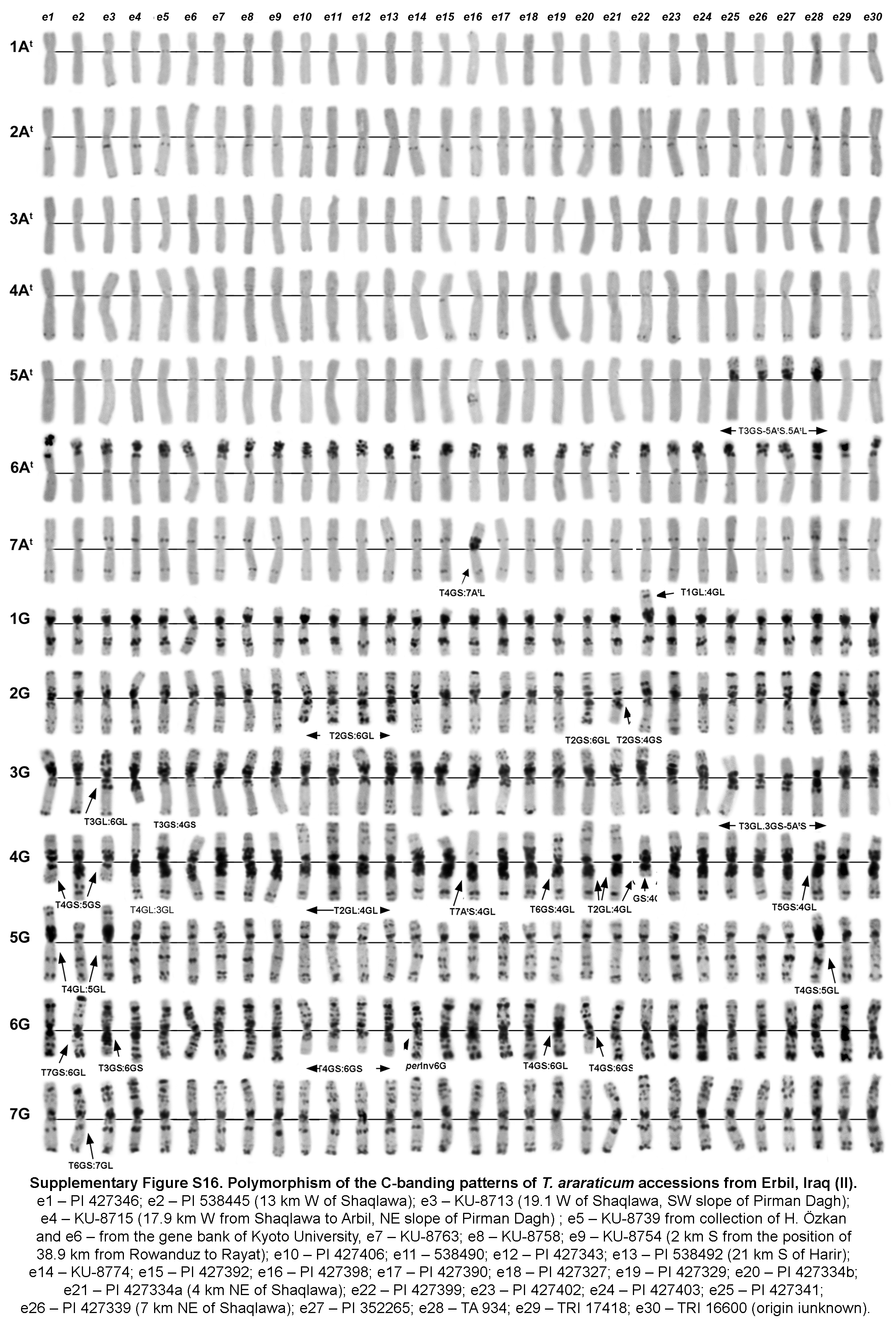

### Fig_S17.tif

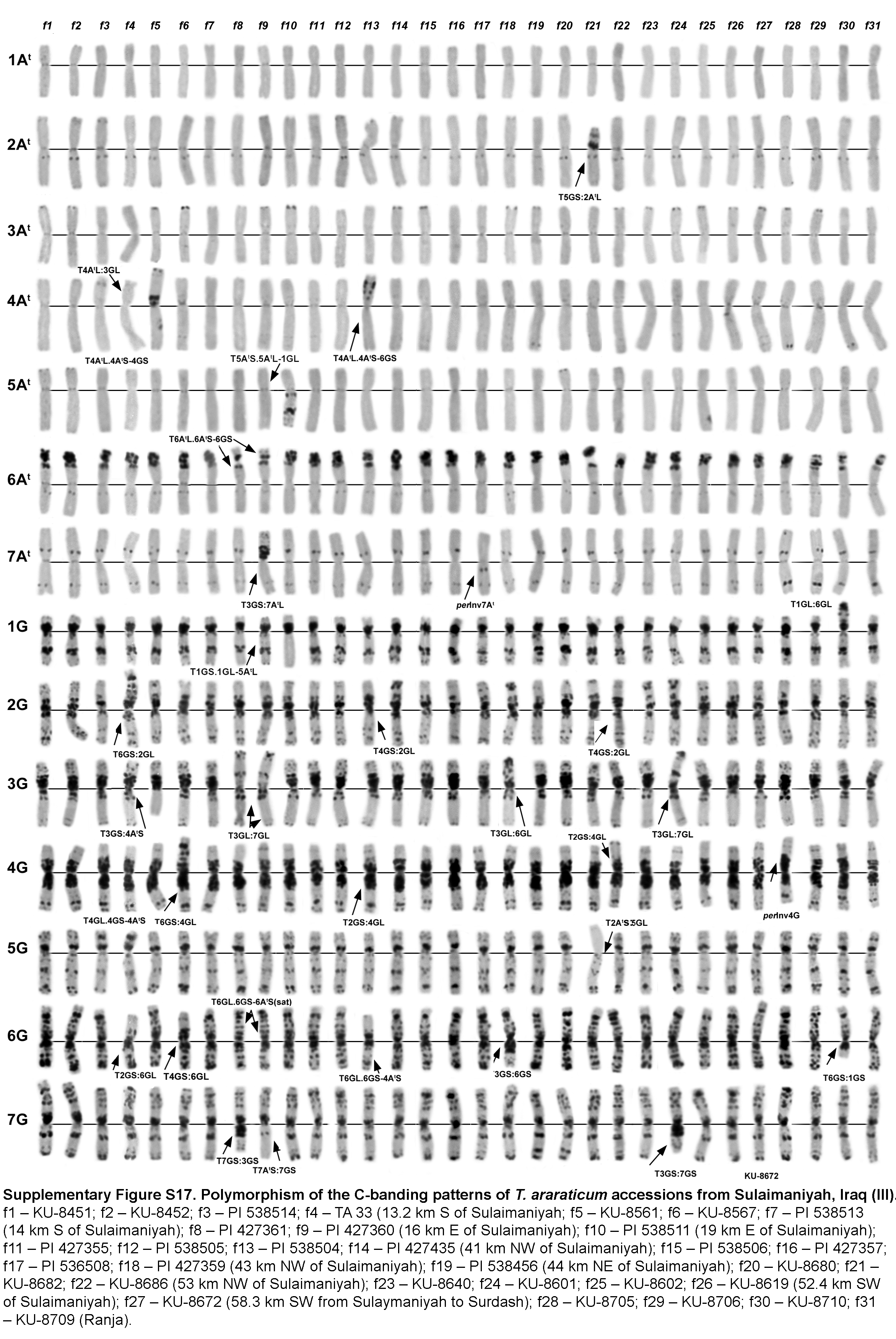

### Fig_S18.tif

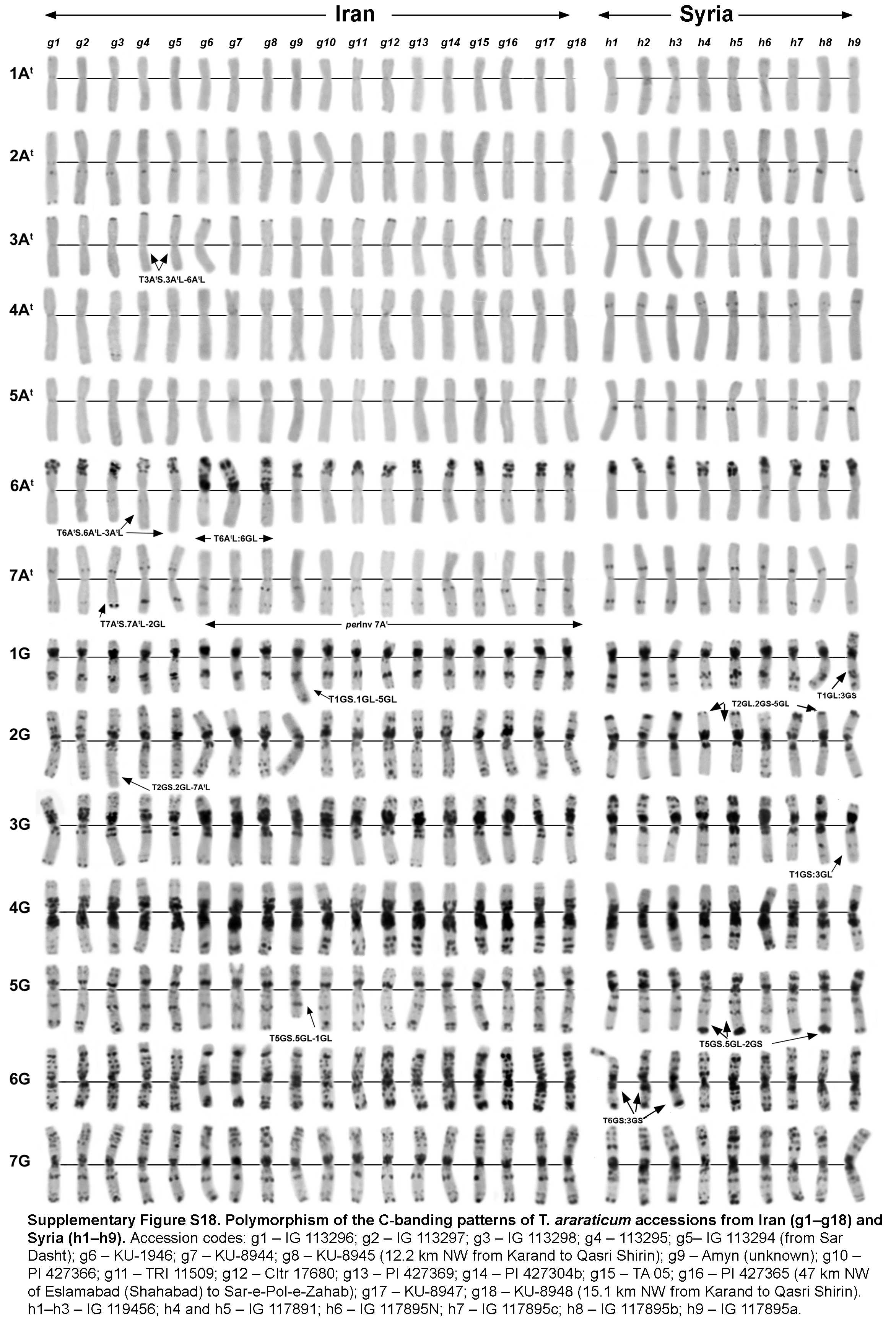

### Fig_S19.tif

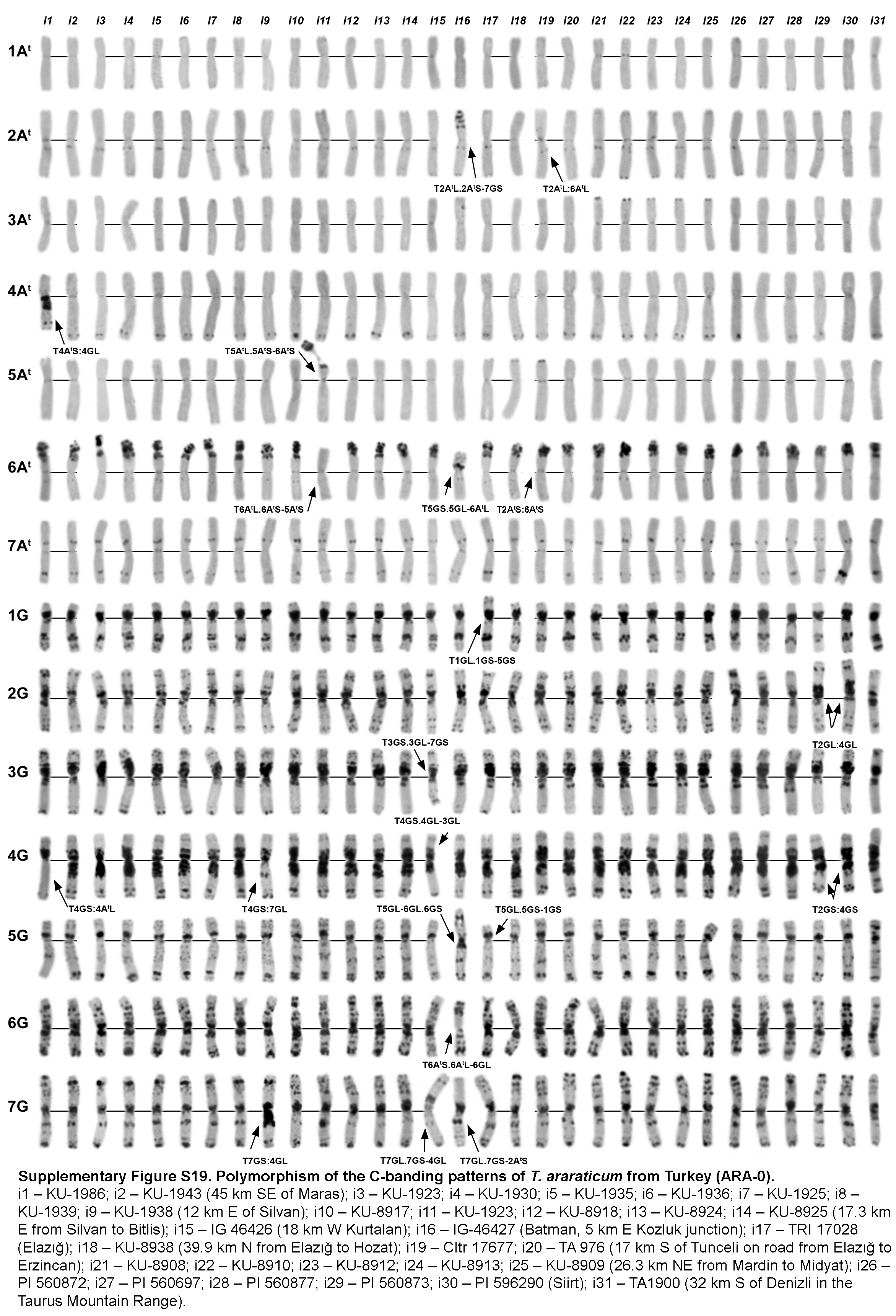

### Fig_S20.tif

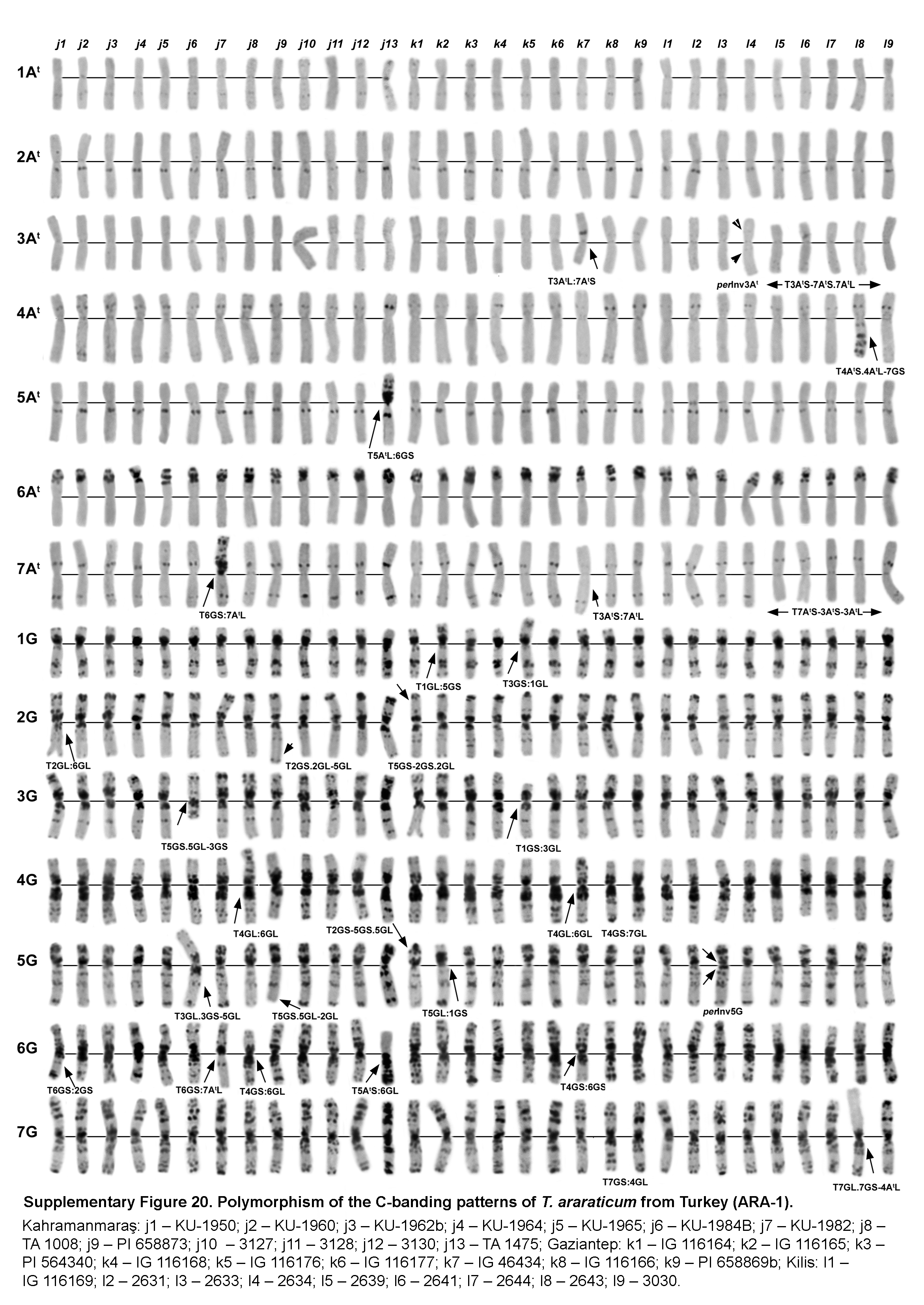

### Fig_S22.tif

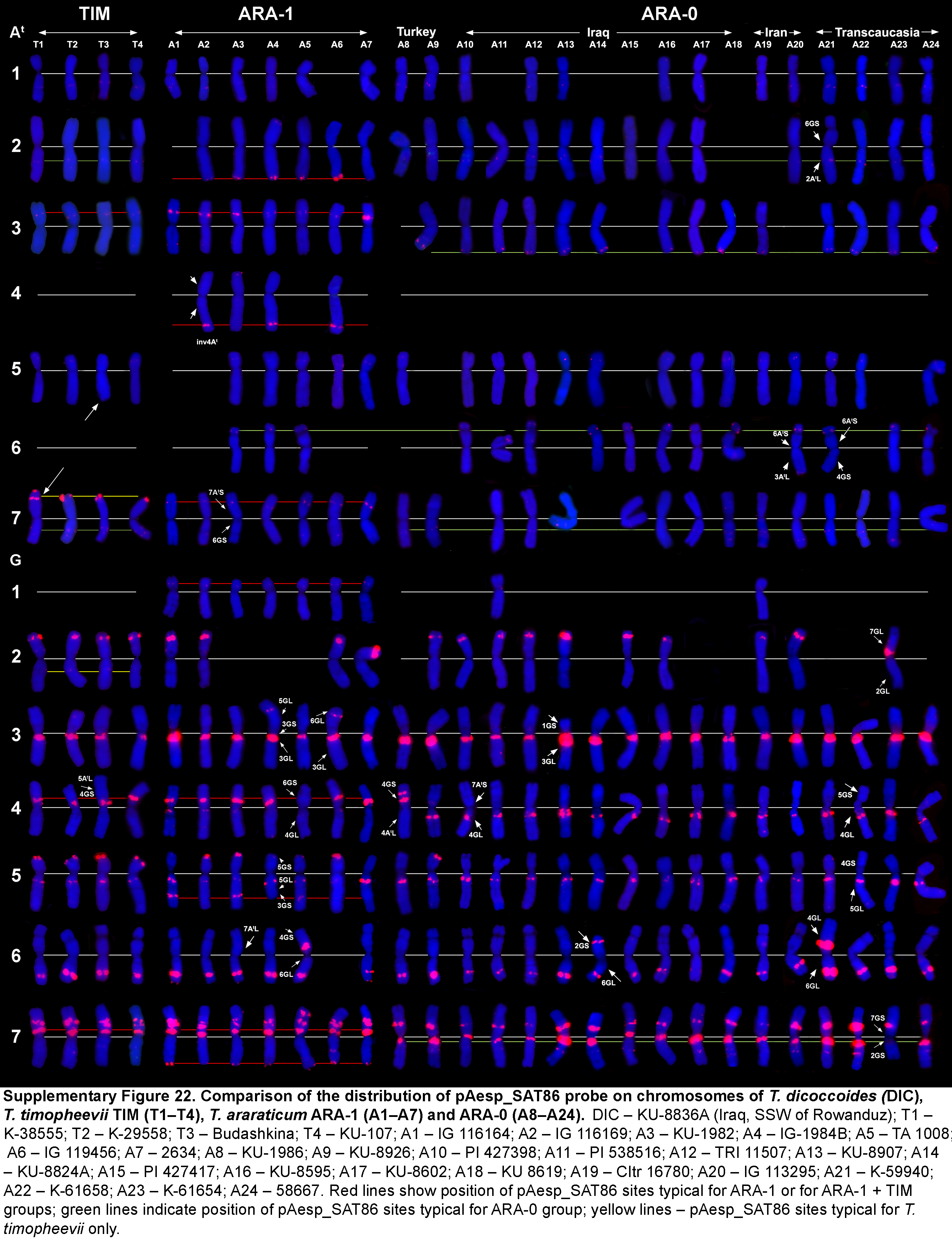

### Fig_S23.tif

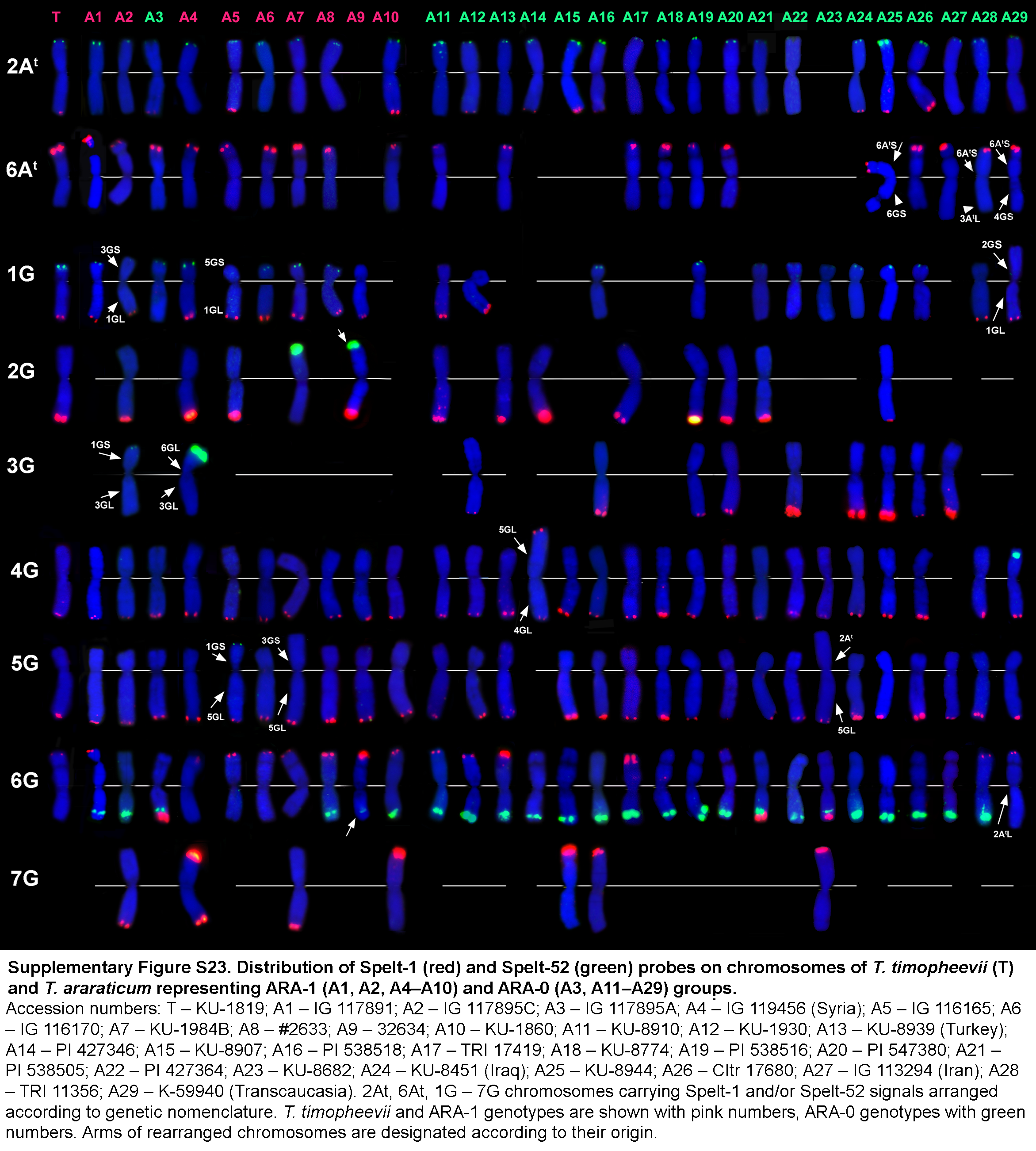
